# Iron□Flavonoid□Peroxidase Signaling Triggers Seed□Coat Remodeling for Rapid Germination

**DOI:** 10.64898/2026.06.01.729177

**Authors:** Deon Mandebere, Ömer Faruk Turan, Mary Lou Guerinot, Seckin Eroglu

## Abstract

Seed germination and survival depend on a sustained water supply. We previously observed that iron (Fe) supply accelerates germination in tomato. Using *Arabidopsis thaliana*, we show here that increased Fe in the spermosphere acts as an extrinsic signal that increases seed coat permeability. Fe triggers structural remodeling of the cuticle covering the endosperm, as evidenced by loss of Auramine O staining in the testa-ruptured seeds. Fe-triggered germination phenotypes are compromised in the flavonoid (*tt4* and *tt7*) and *prx17* mutants, and in peroxidase inhibitor-treated wild-type seeds. We hypothesize that PRX17-driven peroxidase activity is governed by a maternal repressor-derepressor mechanism in which seed coat flavonoids constitutively repress enzyme activity. This repression is relieved by chelation of Fe by flavonoids, allowing PRX17 to remodel the cuticle and accelerate germination. To test this model, we examined the interaction between Fe and peroxidases with quercetin, a major seed coat flavonoid. Upon imbibition, quercetin is deglycosylated and inhibits PRX17 activity. Formation of Fe-quercetin complexes at the chalaza reduces the level of active quercetin, thereby restoring PRX17 activity in a dose-dependent manner. This study suggests that elevated Fe levels in oxygen-deprived, submerged soils prime seeds for rapid germination, a novel control mechanism mediated by seed coat biochemistry.

## Introduction

The transition from a quiescent seed to a germinating seedling is a critical decision in the life cycle of a plant^1^. Consequently, germination timing is tightly regulated by internal and environmental cues that promote or inhibit this process. These factors shift the balance between the antagonistic hormones abscisic acid (ABA), which represses germination, and gibberellins (GA), which promote it^2^. While the endosperm serves as a primary site for integrating hormonal signals^3^, the seed coat (testa) has traditionally been viewed as a static physical barrier that limits water uptake and oxygen permeability through its lipid and phenolic composition^4^. However, recent evidence demonstrates that the seed coat—despite consisting of non-living maternal tissue at maturity—employs sophisticated regulatory mechanisms^5^, including activation of stored peroxidases that facilitate mucilage release^6^.

While ABA-GA interactions fine-tune timing, the seed coat’s hydrophobic barriers function as a major determinant of germination by controlling permeability. In species with physical dormancy, these hydrophobic layers—comprising suberin, cutin, and waxes—can effectively block germination for decades^7^. The biosynthetic pathways of these polymers are well-characterized, involving peroxidases for the polymerization of aromatics and acyltransferases for the esterification of fatty acids to form the functional hydrophobic layers^8,9^. However, the exact physiological processes required to loosen or degrade these barriers and restore permeability during germination are largely unknown.

Class III peroxidases are apoplastic enzymes that accept electrons from phenolic and other substrates to generate radicals. These radicals can then crosslink and polymerize cell wall components, as in lignin and suberin synthesis ^10^ or attack otherwise stable macromolecules, for example through hydroxyl radical-mediated polysaccharide cleavage^11^. Given that germinating seeds accumulate a diverse array of peroxidase isoforms^12^ and phenolic reserves^13^, peroxidases likely participate in multiple physiological processes. To date, only a small subset of peroxidases have been characterized^6,14^. Whether specific peroxidases are involved in barrier remodeling during germination remains unclear. Intriguingly, seed coats exhibit both elevated peroxidase activity^15^ and compromised permeability^16^ when flavonoid metabolism is disrupted.

Flavonoids are among the most abundant phenolic compounds in seeds. Within this group, quercetin predominates in the seed coat, yet it accumulates primarily as an inactive rhamnosylated conjugate, quercitrin. Understanding the biological function of quercitrin has been hampered by significant methodological barriers. For example, the classical histochemical stain DPBA fails to detect quercetin in the seed coat^15^, and there are currently no practical means to track the *in situ* conversion of quercitrin to its active aglycone form. Moreover, this conversion may require a specific rhamnosidase; however, plant-specific rhamnosidases are unknown^17^. Thus, the biological role of quercetin in the seed coat still awaits clarification.

Seeds are a major sink for metal micronutrients. Historically, research on micronutrient accumulation in dicot seeds has focused primarily on the embryo, considering these metals solely as nutrients for seedling establishment^18–21^. Consequently, the spatial distribution and physiological function of metal pools outside the embryo have been largely neglected^22^. However, given that transition metals are essential cofactors for a wide range of biological processes including enzymatic catalysis^23^, and seed coats are known to harbor various active enzymes^5^, non-embryonic metal pools may also participate in the regulation of germination.

Recently, we showed that iron (Fe) in tomato seeds preferentially localized to the chalaza, the primary site of water entry. We demonstrated that mobilization of internal Fe stores or external Fe supplementation significantly accelerates germination^24^. In the current study, we dissect the molecular mechanism underlying Fe-induced germination using *Arabidopsis thaliana.* Its biphasic germination profile, rupture of the testa followed by rupture of the endosperm, enabled us to investigate the impact of Fe on physiological events preceding radicle protrusion. By combining mutant analysis, histochemical profiling, and pharmacological interventions, we show that seed coat Fe, flavonoids, and PRX17 orchestrate seed coat permeability and germination timing.

## Material & Methods

### Plant materials and growth conditions

All *Arabidopsis thaliana* lines and T-DNA insertion mutants used in this study were derived from the Columbia-0 (Col-0) background and obtained from the ABRC stock center (Ohio State University, USA) with the following accession numbers: *prx17-1,* SALK_003180C ; *prx17-2,* SALK_034684C; *tt4-11,* SALK_020583C*; tt7*-7, CS2105578*; vit1-1,* CS66013. Cvi-0 was obtained from NASC.

All chemicals were purchased from Sigma-Aldrich unless stated otherwise. Seeds were surface-sterilized with a solution of 70% (v/v) ethanol and 0.05% (v/v) Triton X-100. For sterilization, approximately 20 µL of seeds were placed in 2 mL Eppendorf tubes, and 1 mL of the solution was added to each tube. The tubes were agitated for 21 min at 110 rpm at room temperature. The sterilization solution was then discarded under a laminar flow hood, and seeds were washed twice with 100% ethanol and air-dried under sterile conditions overnight.

Seeds were sown on Petri dishes containing water agar (0.6% agar, buffered at pH 5.5 with MES) with or without the respective treatments. Plated seeds were stratified for 2 days at 4°C in a refrigerator and then grown under long-day conditions (16/8 h light/dark, 22°C/18°C) at a light intensity of 140 μmol photons m^-2^ s^-1^.

### Seed germination assay

Testa rupture and germination assays were performed on agar medium or filter paper. Sterilized seeds were sown on filter paper or 0.6% (w/v) agar medium buffered with 5 mM MES (pH 5.5) and supplemented with or without 100 µM methimazole, 250 µM Fe(III)-EDTA, 30 µM nicotianamine (NA) (Cayman), or 200 µM desferrioxamine (Des) (Cayman), as specified. At least 30 seeds per genotype were analyzed in three replicates. Germination was defined by radicle emergence and was recorded at different time points. For assessing testa rupture, 30 µM ABA was added to the medium. Testa rupture was defined as exposure of the white endosperm.

### Perls/DAB staining

Perls/DAB intensification was performed according to Roschzttardtz *et al.* (2009) with minor changes. Briefly, all seeds subjected to Perls/DAB staining were either non-imbibed or imbibed in distilled water for 12 h at 4°C. The chalazal and micropylar endosperm/seed coat regions were isolated by a cross-sectional cut using a surgical blade. The isolated caps were fixed with a solution containing methanol: chloroform: glacial acetic acid (6:3:1) for 1 hour under vacuum (500 mbar) as described by Brumbarova and Ivanov (2014). The fixing solution was removed, and seeds were washed 3 times for 1-2 min each with distilled water with gentle shaking. The samples were vacuum infiltrated with Perls solution containing equal volumes of 4% (v/v) HCl and 4% (w/v) K-ferrocyanide (Perls stain solution) for 1 h and washed with distilled water, followed by incubation in a methanol solution containing 0.01 M NaN_3_ and 0.3% (v/v) H_2_O_2_ for 1 h, and then washed with 0.1 M phosphate buffer (pH 7.0). The samples were incubated for 10 min in a 0.1 M phosphate buffer (pH 7.4) containing 0.025% (w/v) DAB, 0.005% (v/v) H_2_O_2_, and 0.005% (w/v) CoCl_2_ (intensification solution). The reaction was stopped by rinsing with distilled water. The caps were mounted on slides for examination under a light microscope and imaged.

### Synchrotron X-ray fluorescence (**μ**XRF)

Seeds were imbibed in distilled water for 24 h, the micropylar and chalazal parts were dissected, and the isolated endosperm caps were immediately transferred to a sample holder between two Ultralene foils and plunged into liquid nitrogen. The samples were maintained in liquid nitrogen during the transfer to the Scanning X-ray Microscope (SXM). Daiquiri software was used to acquire multiple XRF maps at ID21 X-ray Microscopy beamline, European Synchrotron Radiation Facility (Grenoble, France). Elemental localization and distribution data were collected at an excitation energy of 7.3 keV above the Fe K-edge, and a dwell time of 100 ms. Elemental data collected from the sample were fitted and analyzed, and images were generated using PyMCA 5.9.2 software.

### Histochemical stainings

Auramine O staining was performed according to the protocols described by De Giorgi et al. (2015), with minor modifications. Seeds were first incubated on 0.6% agar medium supplemented with or without 30 µM ABA or 30 µM ABA + 10 mM KNO_3_. After testa rupture, 30 seeds were collected and immediately incubated in a 0.005% (w/v) Auramine O solution for 1 h. After staining, the seeds were washed 3 times with phosphate-buffered saline (PBS) at pH 6.5 to remove excess dye and reduce background fluorescence. Stained seeds were mounted on glass slides and examined using a FLOID Cell Imaging Station (Fisher Scientific, Vantaa, Finland) equipped with a GFP filter at 488 nm/532 nm.

Toluidine blue staining was performed to assess seed coat permeability as described by De Giorgi et al. (2015), with minor modifications. Seeds were sown on 0.6% agar medium supplemented with or without 30 μM ABA or 250 µM FeCl_2_. Seeds were stratified for 2 days at 4°C. After stratification, the plates were transferred to a growth chamber under the conditions described above. When testa rupture was observed, about 30 seeds were placed in Eppendorf tubes containing 0.05% toluidine blue and incubated at room temperature for 6 h. Following incubation, seeds were washed with distilled water and embryos were isolated by placing the seeds between microscope slides and applying gentle pressure. The isolated embryos were photographed under a light microscope. All images were acquired using identical illumination and exposure settings to allow for quantitative comparison. Quantitative image analysis was performed using ImageJ (NIH, Bethesda, MD, USA). The “Color Deconvolution” plugin (H DAB vectors) was applied to separate the staining signal. The resulting blue channel images were inverted, and staining intensity was quantified by measuring the mean gray value across the entire embryo. Measurements were performed on at least 15 embryos per treatment.

Peroxidase staining of seeds was performed according to Linkies et al. (2010). Briefly, seeds were sown on Petri dishes containing 0.6% agar medium with 30 µM ABA, with or without 250 µM FeCl_2_. Seeds were collected at the testa rupture stage and incubated in 0.2% w/v TMB (3,5,3′,5′-tetramethylbenzidine-HCl) and 1 mM H_2_O_2_ in 20 mM phosphate buffer, pH 6.5. After approximately 5 minutes of incubation, when the blue coloration indicating peroxidase activity was visible, the seeds were removed from the solution, washed in phosphate buffer (pH 6.5) and imaged. Peroxidase activity was also observed using chloronaphthol, 4-chloro-1-naphthol. Seeds were incubated in freshly prepared staining solution consisting of 5 ml of 20 mM phosphate buffer (pH 6.5), chloronaphthol (30 mg ml^-1^), and 0.6 ml 3% H_2_O_2_ for 5 min. The reaction was stopped by removing the seeds from the staining solution followed by washing three times with phosphate buffer (pH 6.5) for at least 2 min each time.

For the detection of peroxidase activity *in vitro*, reactions were performed with TMB as the substrate. The reaction mixture contained 20 mM phosphate buffer (pH 6.5), 0.2% (w/v) TMB, 1 mM H_2_O_2_, and 0.001 g ml^-1^ horseradish peroxidase (HRP) with or without 100 μM salicylhydroxamic acid (SHAM), 100 μM quercetin, and 250 μM Fe(III)-EDTA. All reactions were incubated at room temperature for 5 min. The resulting colorimetric changes were imaged. The formation of a blue color indicated peroxidase-oxidized TMB. Absorbance was measured at 430 nm using a UV–VIS spectrophotometer.

### Flood simulation

Non-fertilized, top soil was collected from the campus of the Middle East Technical Universiy, Ankara, Turkey. After drying in an oven at 60°C for two days, the soil is sieved (<2mm) and homogenized. To facilitate anaerobic conditions and simulate microbial activity, soil was mixed with finely ground dried leaf litter collected from the same site (9:1, soil:leaf powder).

To simulate flooded conditions, 100 g soil was placed in 800 ml beaker following 200 ml of MES solution (2.5 mM at pH 5.5) addition. This solution is thoroughly mixed. The beaker was sealed airtight with Parafilm and then incubated in a growth chamber with a 16/8 h light/dark cycle at 22°C/18°C (day/night) and a light intensity of 140 μmol m^-2^ ^s-1^. For the prolonged flood, this mixture was incubated for three days, while for the control (acute flood) condition, the soil solution was prepared fresh. Sludges were placed into the wells of 24-well plates (n=4) to reach a height of 1.5 cm. Seeds were sown on the sludges and placed in a growth chamber, and germination rates were calculated.

To track the impact of prolonged flooding on Fe availability, oxidation-reduction potential (ORP) was monitored using an ORP meter (Milwaukee SE300 probe) as a proxy for Fe solubilization. The probe was left in the soil for 10 minutes until a stable *mV* reading was obtained. The impact of prolonged flooding on Fe levels was further determined colorimetrically using the ferrozine assay. Sludges described above were stirred vigorously for 5 minutes to ensure a uniform mixture, then allowed to settle for 5 minutes at room temperature. The supernatant was collected and filtered through a 0.25 µm syringe filter. Filtrate was mixed with ferrozine (200 µM final concentration), and the absorbance was measured at 562 nm using a SOIF UV-5100 UV-VIS spectrophotometer (Shanghai Metash, China).

## Results

### The seed coat contains a mobile chalazal Fe store

Previously, we observed that tomato seeds preferentially accumulated Fe in the chalaza and presented evidence that this Fe accumulation facilitates germination^24^. To reveal the genetic mechanisms underlying this phenomenon, we employed *Arabidopsis thaliana* as a model system. Although Fe-enriched sub-regions in Arabidopsis embryos are well documented^18,21,26,27^, specific Fe stores within the chalaza have not been previously reported. To investigate whether Fe accumulation in the chalaza is conserved in Arabidopsis, we used X-ray synchrotron tomography to examine intact, dry seeds. Consistent with previous reports^18^, we observed characteristic Fe accumulation patterns around the provascular strands of the Arabidopsis embryo (Figure 1A). We also observed a previously unreported ring-like accumulation of Fe in the chalazal seed coat, alongside dispersed Fe puncta. The signal intensity of the ring was similar to that of the provascular strands. Since provascular strands accumulate Mn^26^ in addition to Fe, we examined whether the accumulation of metals in the chalaza was specific for Fe. Mapping several metals in the ring, including Mn, showed that metal overaccumulation in this region was specific to Fe (Figure 1B).

**Figure 1:**
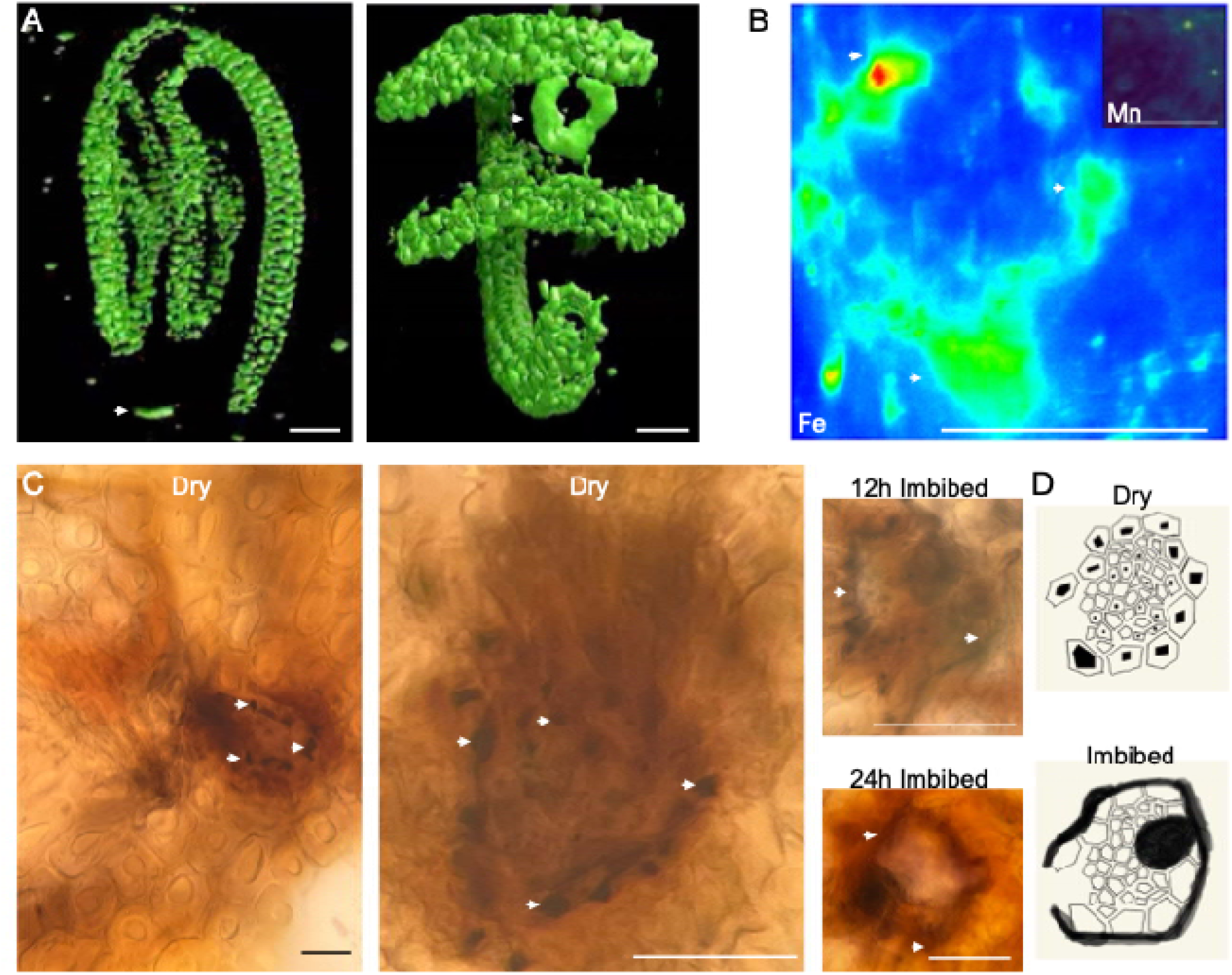
Specific accumulation and remobilization of Fe in the chalaza. A, Fe distribution in dry intact Arabidopsis seeds visualized by Synchrotron X-ray Fluorescence (SXRF) microscopy. Left, lateral view; right, chalazal view. Arrowheads indicate the specific enrichment of Fe in the chalaza. B, comparative elemental maps of Fe and manganese (Mn)(inset) in chalazal regions dissected from 24-hour-imbibed seeds analyzed by SXRF. Note that Mn does not show the specific accumulation pattern observed for Fe. C, histological localization of Fe in the chalaza using Perls/DAB staining. Images focus on the chalaza of the cross sections of seed coats obtained from dry or imbibed seeds. Black signals indicate Fe accumulation. Arrowheads point to the specific cellular localization of the stain. D, schematic of Fe accumulation in the chalaza of dry or imbibed seeds. Note that in dry seeds Fe accumulates as clear spots, but in imbibed seeds as a continuous line at the contours of the cells. All analyses were performed independently at least three times with a minimum of 5 samples per replicate; representative images are shown. Bar, 0.01mm.

Next, we investigated the fate of chalazal Fe upon imbibition, the earliest phase of seed germination. Perls/DAB histochemical staining of dry seeds resolved the SXRF signal as a highly ordered pattern, showing Fe deposits localized to the central lumens of epidermal cells surrounding the chalazal abscission zone (Figure 1C, D). Upon imbibition, Fe relocalized from the cell centers to the cell contours. These data suggest that Arabidopsis seeds possess a specific, mobile pool of Fe in the chalaza that is released into the extracellular space during the early stages of germination.

### Chalazal Fe controls seed coat permeability

Having established that chalazal Fe accumulation is a conserved feature in *Arabidopsis*, we investigated whether this specific pool contributes to germination kinetics similar to our findings in tomato^24^. In tomato seeds, mobilization of internal Fe stores correlated with germination speed^24^. Our results show that in Arabidopsis seeds, treatment with the Fe mobilizer nicotianamine (NA) significantly accelerated germination, reducing the time to 50% germination by approximately 3 hours. Conversely, the Fe immobilizer desferrioxamine (Des) did not delay germination (Figure 2A).

**Figure 2:**
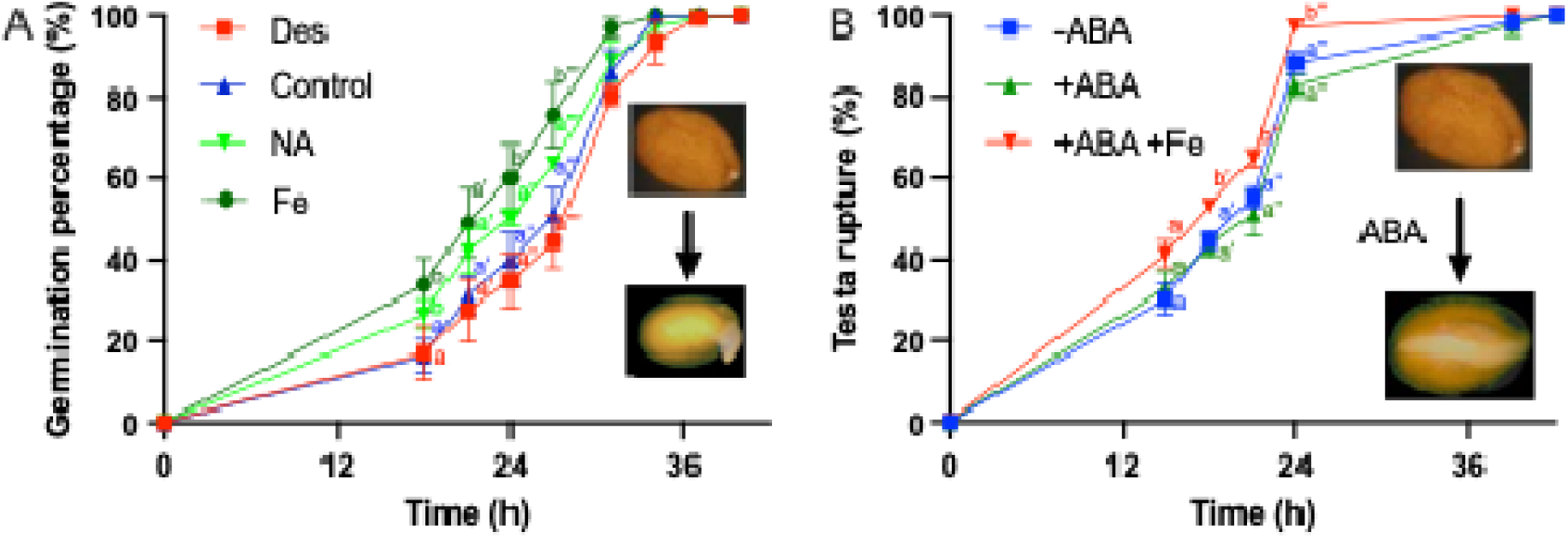
Mobilization of Fe stores or external Fe supplementation facilitates germination. A, impact of Fe chelators and external Fe on germination kinetics. Seeds were imbibed on filter paper wetted with MES buffer (pH 5.5) supplemented with the Fe mobilizer nicotianamine (NA, 5 µM), the Fe chelator desferrioxamine (Des, 200 µM), or Fe-EDTA (250 µM). B, impact of Fe on testa rupture in the presence of ABA. Seeds were imbibed with or without ABA (30 µM) and Fe-EDTA (250 µM) to uncouple seed-coat rupture from radicle protrusion. Insets show images defining the scored developmental stages of germination (radicle protrusion, left) and of testa rupture (seed coat split, right). Data represent means ± SEM (n = 4 replicates of > 30 seeds). Letters indicate significant differences between the means of the control (left) or +ABA control (right) and treatments at each time point (two-way ANOVA with Tukey’s post-hoc test, p ≤ 0.05).

Given that Des treatment significantly delays germination in tomato^24^ but it did not in Arabidopsis, we hypothesized that the lack of a significant effect in wild-type Arabidopsis might be due to a naturally lower abundance of chalazal Fe stores compared with tomato. To identify Arabidopsis mutant seeds with higher Fe accumulation in the chalaza, we revisited the literature for published Fe maps in Arabidopsis seeds and noticed that *vacuolar iron transporter 1* (*vit1*) mutants appeared to accumulate more Fe in the chalaza^18^. Consistent with our hypothesis, *vit1* seeds displayed a greater sensitivity to chelation: while NA still accelerated germination (∼2 hours), Des treatment caused a 2.5-hour delay (Figure S1). This amplified response in the high-Fe *vit1* background supports the model that germination kinetics are modulated by mobilizable chalazal Fe pools.

Similar to NA, germination speed is also induced by Fe supplementation (Figure 2A). All tested forms of Fe (FeCl_2_, FeCl_3_, Fe-EDTA, and Fe-PDMA) accelerated germination in a concentration-dependent manner (Figure S2). We also tested whether the impact on germination was specific to Fe and found that neither ZnCl_2_ nor MnCl_2_ increased germination speed (Figure S2). These experiments indicate that the response is specific to Fe. Next, we explored the stage at which Fe acts during germination. Considering that Fe relocalized soon after imbibition (Figure 1) we hypothesized that Fe functions prior to radicle protrusion. In Arabidopsis, germination proceeds in two steps: testa rupture followed by endosperm rupture (radicle protrusion)^262728^. By supplementing seeds with ABA, we blocked endosperm rupture, allowing us to isolate the testa rupture event. As expected^28^, while ABA prevented radicle emergence, it did not prevent the physical breakage of the seed coat. Strikingly, Fe supplementation accelerated the rupture of the testa (Figure 2B), confirming that Fe targets the mechanical integrity of the seed coat during the early phases of germination.

### Fe triggers rapid remodeling of the endosperm-associated cuticle

Previous studies have established that defects in the seed cuticle lead to increased permeability and testa rupture under ABA supplementation^29^. To determine if Fe-accelerated rupture proceeds via a similar mechanism, we assessed the impact of Fe on seed coat permeability and cuticle integrity. We first monitored permeability using a lipophilic dye, toluidine blue (TBO). In the absence of Fe, embryos displayed only a faint blue coloration, indicating limited dye uptake; whereas Fe supplementation resulted in increased coloration (Figure 3) This Fe-dependent increase in permeability was observed in both intact and testa-ruptured seeds (Figure 3B), confirming that Fe compromises the barrier function of the coat.

**Figure 3:**
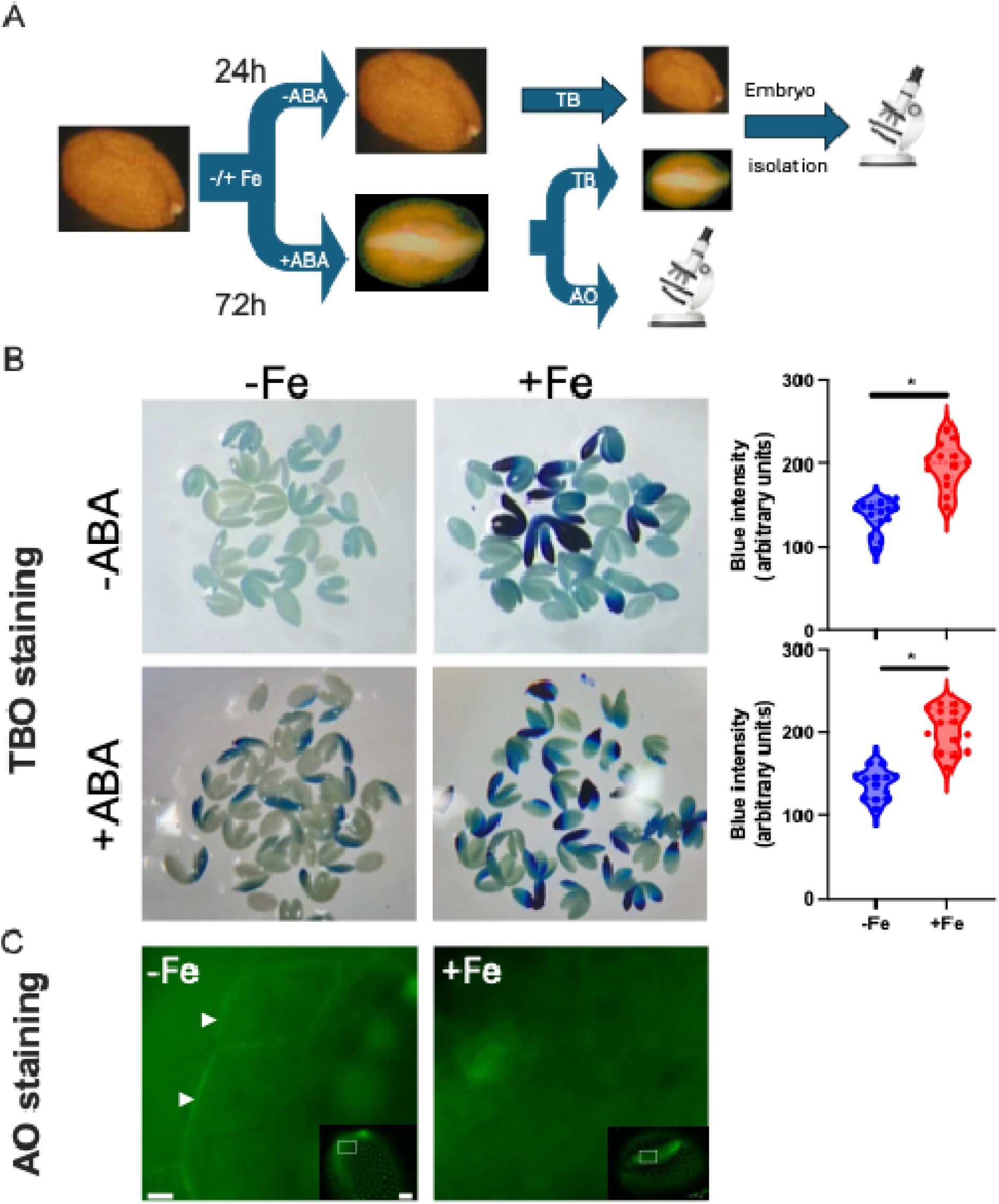
Fe increases seed-coat permeability and remodels the endosperm-associated cuticle. A, schematic illustration of the experimental design. Permeability was assessed in intact seeds (24 h, -ABA) and testa-ruptured seeds (72 h, +ABA) by monitoring the penetration of toluidine blue (TBO) dye into the isolated embryos. The testa-ruptured state also allowed for the direct microscopic examination of the cuticle. B, representative images and quantification of TBO-stained embryos isolated from seeds imbibed with or without 250 µM Fe-EDTA. Note the significantly higher stain intensity in Fe-treated embryos (+Fe) in both intact (-ABA) and testa-ruptured (+ABA) conditions, indicating increased permeability. Violin plots show the quantification of blue pixel intensity (n > 15 embryos). Asterisks indicate significant differences based on an unpaired t-test (p < 0.001). C, epifluorescence microscopy of auramine O (AO) stained testa-ruptured seeds focused on the exposed endosperm surface. In control seeds (-Fe), white arrowheads indicate the fluorescent signal defining the cutinized periclinal walls of the inner integument 1 (ii1) layer. This signal is undetectable in Fe-treated seeds. Insets show the whole ruptured seeds. Scale bar = 10 µm and 125 µm (insets).

To identify the specific barrier affected, we visualized the endosperm-associated cuticle using auramine O staining. In ruptured control seeds, the exposed endosperm surface is covered by the cutinized periclinal cell walls of the inner integument 1 (ii1) layer^30^. Consistent with previous reports^30^, auramine O strongly stained this layer in control seeds, defining the cellular contours of the ii1 layer (Figure 3C), in agreement with the previous observations^30^ . Strikingly, this specific auramine O signal was undetectable in Fe-imbibed seeds. This loss of staining suggests that Fe treatment triggers rapid structural remodeling of the endosperm-associated cuticle, thereby removing the primary physical restraint to germination.

### PRX17 mediates Fe-dependent cuticle remodeling

Because non-enzymatic scission of cell wall polymers can be driven by reactive oxygen species (ROS)^31^, we hypothesized that the observed cuticle remodeling during imbibition might be mediated by hydroxyl radicals being generated. A primary source of apoplastic ROS in seeds is the NADPH oxidase pathway mediated by respiratory burst oxidase homologues RBOHs^32^. The major RBOH in the endosperm is RBOHD.^33^ In *rbohD* mutants, Fe supplementation still accelerated germination (Figure S3), suggesting that the Fe-induced mechanism operates independently of this canonical ROS-generating pathway.

Class III peroxidases are alternative drivers of radical generation known to be activated during imbibition^34^. To visualize their activity, we used 3,3’,5,5’-tetramethylbenzidine (TMB) staining. In control seeds, we observed basal peroxidase activity localized to the ruptured endosperm surface and, notably, a distinct circular domain in the chalaza (Figure 4A). Supplementation with Fe-EDTA significantly intensified the TMB signal in both regions, suggesting that Fe acts as an activator of these seed coat peroxidases.

**Figure 4:**
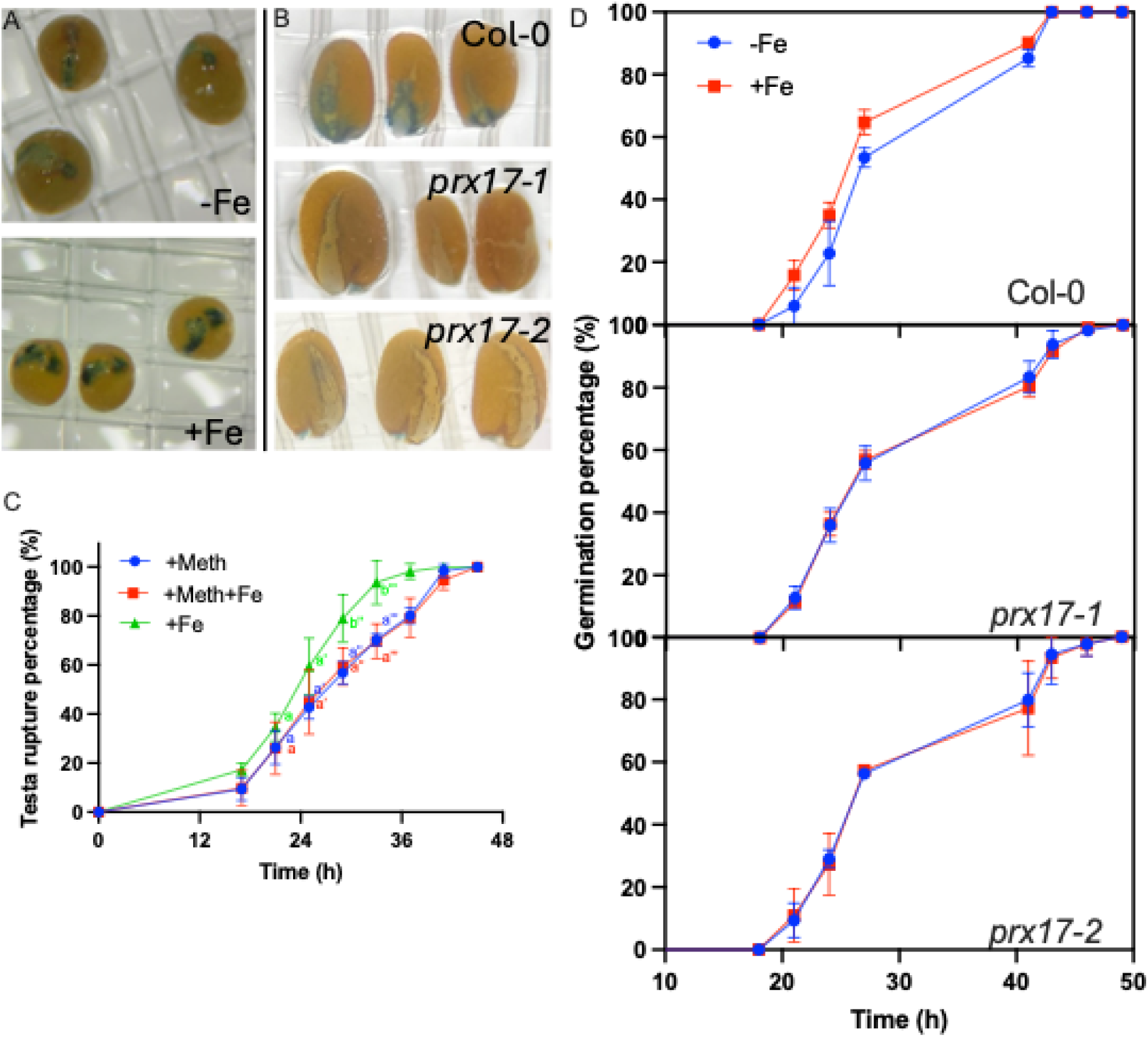
Fe-triggered germination requires PRX17. A, TMB staining of testa-ruptured seeds imbibed with or without 250 µM Fe-EDTA in the presence of 30 µM ABA. Blue coloration indicates peroxidase activity. B, TMB staining of wild-type (Col-0) and *prx17* mutant seeds imbibed with 250 µM Fe-EDTA. Representative images were selected from at least 15 seeds per genotype. C, Time course of testa rupture in the presence of the peroxidase inhibitor methimazole. Seeds were imbibed with 30 µM ABA and combinations of 250 µM Fe-EDTA and methimazole (Meth). Letters indicate statistically significant differences between the control (+Meth) and treatments at each time point (two-way ANOVA with Tukey’s post-hoc test, *p* < 0.05; n = 4 replicates of 30 seeds). D, germination time course of Col-0 and *prx17* mutants imbibed with or without 250 µM Fe-EDTA. Error bars represent ± SEM (n = 4). Two-way ANOVA revealed a significant difference between Fe treatments for Col-0 (*p* = 0.038), but not for *prx17-1* (*p* = 0.34) or *prx17-2* (*p* = 0.61).

To determine if the observed cuticle remodeling is causally linked to Fe-induced peroxidase activity, we treated seeds with the peroxidase inhibitor methimazole. In the presence of the inhibitor, the auramine O signal remained intact on the ii1 layer regardless of Fe supplementation (Figure S5). This confirms that peroxidase activity is required for the Fe-mediated loss of cuticle integrity.

Since auramine O is a broad-spectrum fluorescent probe that can also bind lignin and suberin^35^, we sought to verify that the lost signal represented cuticle (cutin) rather than other cell wall polymers. We stained seeds with basic fuchsin (lignin-specific) and nile red (suberin-specific) and compared the patterns to those obtained with auramine O. While basic fuchsin stained the ii1 layer covering the endosperm, this signal remained unchanged by Fe treatment, ruling out lignin remodeling. Conversely, nile red failed to stain the ii1 layer entirely, indicating a lack of significant suberin deposition in this region. Collectively, these data demonstrate that the Fe-triggered loss of auramine O signal corresponds specifically to the remodeling of the cuticle, and this process is dependent on Fe-activated peroxidases (Figure S4).

To pinpoint the specific peroxidase isoform responsible for this activity, we screened *Arabidopsis* T-DNA insertion lines of candidate class III peroxidase genes for a loss of the Fe-induced TMB signal during imbibition. The candidate genes predominantly included the ones reported to be active in the seed coat during development such as AT5G64100, AT5G39580, AT1G07890, AT5G64120, AT2G18980, AT3G52960, At3g28200, AT1G14540, At4g36430, and At2g22420^36^. This reverse genetics approach identified two independent alleles of *PEROXIDASE 17* (*PRX17*). Both *prx17-1* and *prx17-2,* which were shown to be loss of function mutants^37^, failed to display the characteristic TMB signal in the chalaza or ruptured endosperm, even in the presence of Fe supplementation (Figure 4B). We corroborated these findings using chloronaphthol, an alternative peroxidase substrate with distinct solubility properties. While wild-type seeds exhibited strong staining over the exposed endosperm upon Fe treatment, this activity was absent in the *prx17* mutants (Figure S6).

We next confirmed the functional necessity of the PRX17 enzyme. Treatment with the peroxidase inhibitor methimazole completely abolished the Fe-induced acceleration of germination in wild-type seeds (Figure 4C). Similarly, *prx17* mutants were insensitive to Fe supplementation, germinating at the same basal rate regardless of the Fe status (Figure 4D). Consistently, *prx17* mutants also failed to undergo Fe-triggered cuticle remodeling. Crucially, this defect was not due to a permanent structural alteration in the cuticle itself, as the auramine O signal could be successfully removed *in vitro* by treating *prx17* seeds with exogenous horseradish peroxidase (HRP) (Figure S7). Collectively, these data demonstrate that PRX17 is the essential enzyme linking Fe perception to cuticle remodeling.

### Fe chelates seed flavonoids

The increase in peroxidase activity following Fe supplementation was a surprising observation, given that it cannot be attributed to *de novo* biosynthesis, since the seed coat is dead and metabolism is suppressed by ABA. Since the seed coat is a well-known source of various germination inhibitors^38^, we suspected that the increase in peroxidase activity caused by Fe supplementation may neutralize peroxidase inhibitors. This reminded us that while we were examining the impact of various forms of Fe on germination (Figure S2), unchelated Fe (i.e., FeCl_2_ and FeCl_3_) formed black precipitates in the chalaza (Figure 5A). Revisiting our previous experimental results, we noticed that the pattern of this blackening spatially colocalized with regions of preferential Fe localization and peroxidase activity, forming a circle in the chalaza and a less apparent spot in the micropyle (Figure 5A). This phenomenon appears conserved; tomato seeds also exhibited Fe-dependent blackening that was reversible by EDTA washing (Figure S4). We hypothesized that this color change resulted from the chelation of specific peroxidase inhibitors by Fe.

**Figure 5:**
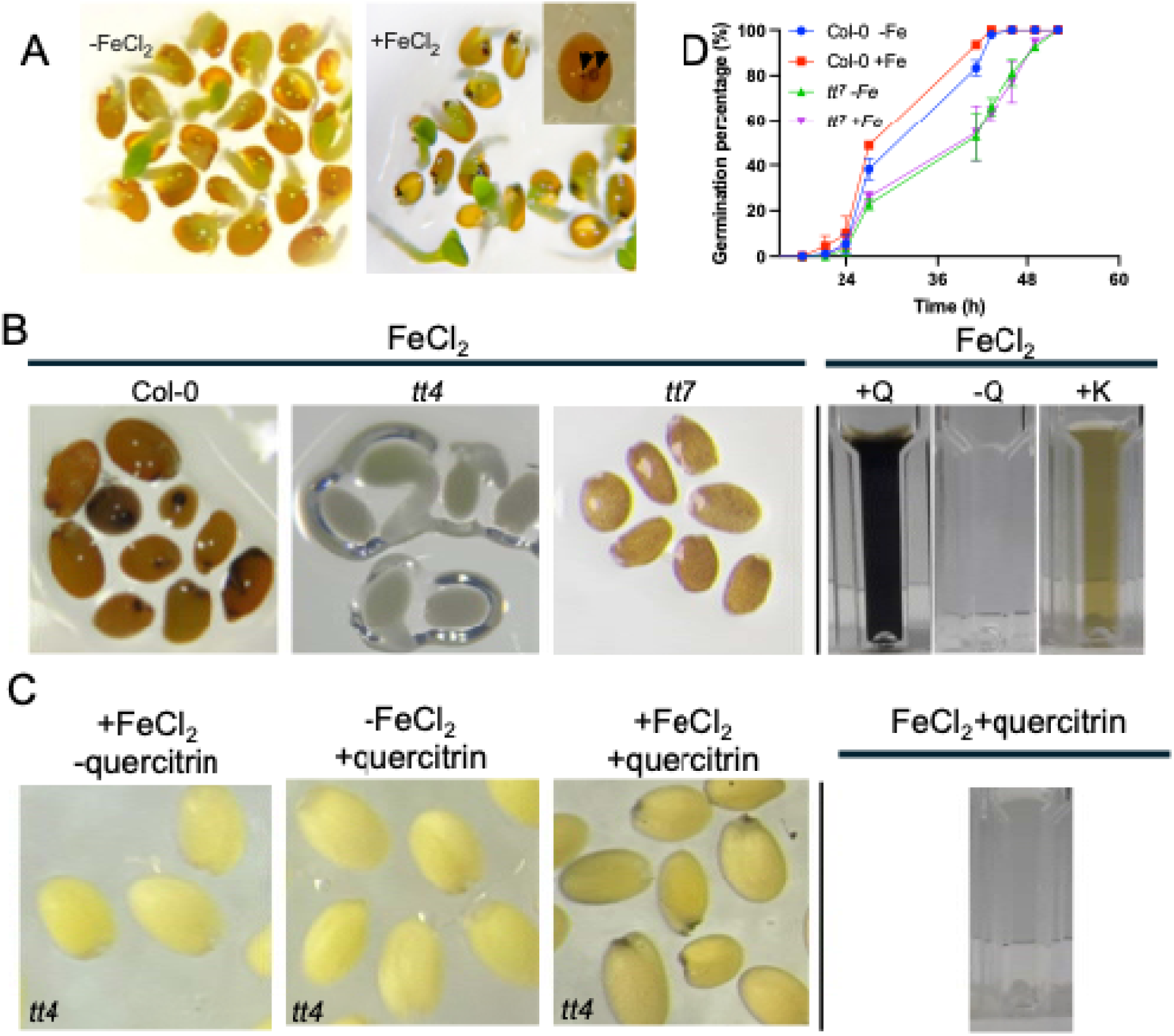
Flavonoids in the seed coat are essential for Fe-triggered blackening and germination. A, development of black coloration in the presence of FeCl_2_. Wild-type seeds were imbibed on filter paper with or without FeCl_2_. Inset, magnified view showing the specific coloration pattern (black arrowheads) localized to the chalaza and micropyle prior to testa rupture. The blackening was consistent between different seed batches. B, left, impact of flavonoid pathway mutations on FeCl_2_-dependent coat coloration. *tt4* (flavonoid-deficient) shows no color; *tt7* (lacking catechol-type flavonoids like quercetin/PAs and over-accumulating kaempferol) develops orange. Right, *in vitro* colorimetric assay of flavonoid-FeCl_2_ mixtures. +Q: quercetin + FeCl_2_ (black); -Q: buffer + FeCl_2_ (colorless); +K: kaempferol + FeCl_2_ (orange). Note the phenotypic resemblance between *tt7* seeds and the kaempferol standard. The coat coloration was consistent between different batches and observed on all seeds without exception. The medium contained 30 µM ABA to prevent radicle protrusion. C, left, *in situ* reconstitution of the blackening response. Seeds of *tt4* mutant plants were imbibed in FeCl_2_ with or without exogenous quercitrin (quercetin-rhamnoside). Note that blackening occurs only when both Fe and quercitrin are present. Right, *in vitro* mixture of quercitrin and FeCl_2_ remains colorless, indicating that the sugar moiety prevents chelation and must be removed *in vivo*. D, germination time course of Col-0 and *tt7* seeds supplemented with or without 250 µM Fe-EDTA. Error bars represent ± SEM (n = 4 independent replicates of 30 seeds). Two-way ANOVA revealed a significant main effect of Fe treatment only for Col-0 (*p* = 0.034), whereas *tt7* remains insensitive to Fe supplementation.

Flavonoids are abundant seed-coat metabolites known to chelate metals and inhibit oxidative enzymes^13,39^. To test their involvement in the modulation of peroxidase activity, we treated *transparent testa 4* (*tt4)* mutants—which are devoid of all flavonoids—with FeCl_2_. Seeds of *tt4* mutants failed to turn black upon treatment with Fe, confirming that flavonoids are the reactants (Figure 5B).

Flavonoids are a large class of specialized metabolites. To pinpoint the compound responsible for blackening, we screened Arabidopsis lines that carry a T-DNA mutation in various genes in the flavonoid pathway. This approach identified *transparent testa7* (*tt7)*, encoding flavonoid 3’-hydroxylase (F3’H), the enzyme catalyzing the biosynthesis of quercetin. Consequently, *tt7* seeds lack quercetin and proanthocyanidins but overaccumulate kaempferol^40^. Unlike wild-type seeds, *tt7* mutants developed a distinct orange-brown color in the presence of Fe. This is consistent with *in vitro* data showing that Fe forms black complexes with quercetin but orange complexes with kaempferol (Figure 5B). While we cannot rule out the contribution of proanthocyanidins to the blackening response, this spectral shift confirms that the reaction can depend on the catechol group present in quercetin and its derivatives.

However, a biochemical paradox remained: flavonoids including quercetin in the seed coat are often stored as inactive glycosides (e.g., quercitrin)^41^. We observed that, unlike the aglycone quercetin, quercitrin does not turn black when mixed with FeCl_2_ (Figure 5B), likely because the sugar moiety sterically hinders Fe chelation^42^. This implies that for *in vivo* blackening to occur, the sugar moiety must be removed.

To assess this bioconversion, we performed a reconstitution assay using quercetin as a representative substrate, given its abundance and structural suitability for Fe chelation. When *tt4* seeds were incubated with inactive quercitrin and FeCl_2_, they developed a strong black precipitate specifically in the chalaza (Figure 5C). This confirms that the chalazal seed coat possesses intrinsic enzymatic activity (presumably a rhamnosidase) capable of converting inactive glycosides into active, Fe-chelating aglycones upon imbibition. Interestingly, expression of the gene encoding the quercetin biosynthetic enzyme, FLS1, spatially and temporally overlaps with that of PRX17 according to publicly available transcriptome resources (Figure S8 and Figure S9).

### Fe and quercetin antagonistically regulate peroxidase activity

The precise spatial convergence of remobilized chalazal Fe, the site of quercetin activation indicated by blackening, and the domain of peroxidase activity (Figure S10) led us to hypothesize that a tripartite interaction between these components orchestrate germination kinetics.

To test this model, we reconstituted the regulatory module *in vitro* using horseradish peroxidase (HRP) and TMB. While HRP efficiently oxidized TMB to produce a distinct blue color, the addition of quercetin completely suppressed this enzymatic activity. Strikingly, subsequent titration of Fe into the inhibited mixture restored peroxidase activity in a dose-dependent manner (Figure 6). This confirms that Fe can functionally rescue peroxidase activity by chelating the inhibitory flavonoid quercetin.

**Figure 6:**
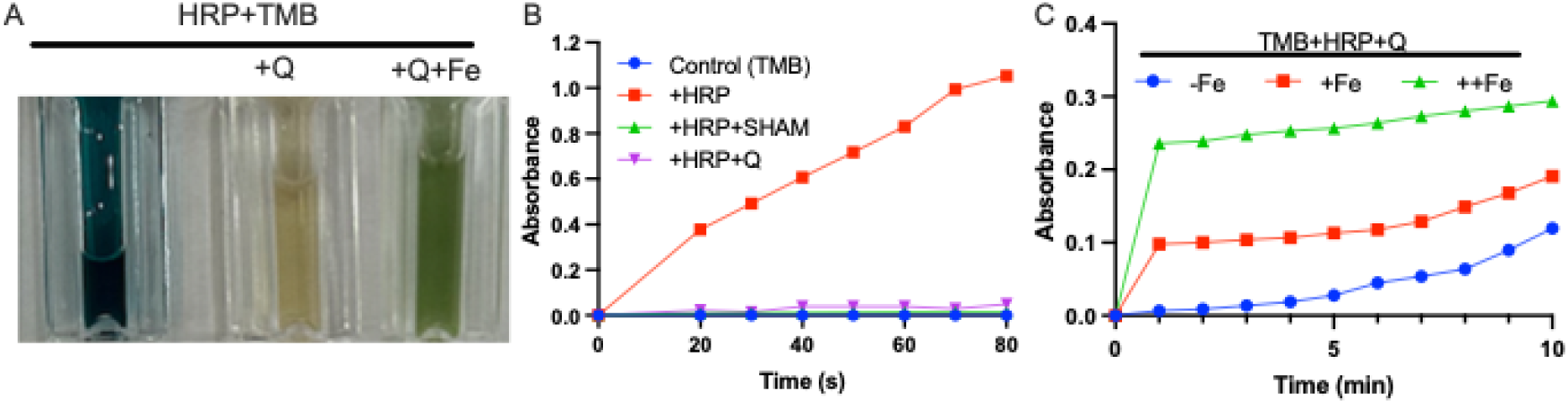
Quercetin inhibits peroxidase activity *in vitro* and this inhibition is relieved by Fe chelation. A, qualitative assessment of horseradish peroxidase (HRP) activity using the chromogenic substrate TMB. From left to right: HRP+TMB (positive control showing blue oxidized product); +Q: HRP+TMB supplemented with 100 µM quercetin (Q), showing inhibited color formation; +Q+Fe: the inhibited reaction supplemented with 100 µM Fe-EDTA (Fe), showing restoration of activity. Images were acquired 1 hour after mixing. B, quantitative time course of HRP activity monitored by absorbance of the oxidized TMB product. The reaction with HRP alone (red squares) was compared to reactions inhibited by SHAM (salicylhydroxamic acid, a known peroxidase inhibitor; green triangles) or Q (quercetin; purple inverted triangles). TMB alone (control, blue circles) shows negligible auto-oxidation. C, dose-dependent restoration of quercetin-inhibited HRP activity by Fe-EDTA. The basal inhibited reaction (TMB+HRP+Q, -Fe; blue circles) was supplemented with a low (+Fe; red squares) or high (++Fe; green triangles) concentration of Fe-EDTA, and absorbance was monitored for 10 minutes. Abbreviations: HRP, horseradish peroxidase; SHAM, salicylhydroxamic acid; Q, quercetin; TMB, 3,3’,5,5’-tetramethylbenzidine; Fe, Fe-EDTA.

We further validated this antagonistic mechanism *in situ* by specifically targeting the Fe-induced PRX17 activity in the seed coat. Using chloronaphthol staining, we observed robust peroxidase activity in the chalaza of Fe-treated seeds. However, the simultaneous application of exogenous quercetin effectively repressed this Fe-induced activity (Figure S11). Collectively, these data establish a direct biochemical mechanism: quercetin acts as a potent inhibitor of PRX17, and Fe functions as a derepressor by sequestering the flavonoid, thereby allowing PRX17 to degrade the cuticle.

### Fe-triggered germination phenotype is relevant during prolonged floods

In waterlogged or flooded soils, oxygen depletion drives the reduction of insoluble Fe(III) to soluble Fe(II), a process often accelerated by soil microbes engaging in Fe respiration, which elevates Fe to levels toxic to plants^43,44^ We hypothesized that the Fe-triggered germination phenotype can be reconstituted in flooded soils. We submerged soil in water for a few days to simulate flooding while monitoring Fe availability. As expected, Fe^2+^ levels continuously increased during submergence (Figure S1). Wild type seeds sown on soil that had been flooded for 3 days (prolonged flood) germinated faster than seeds sown on 0 days flooded soil (acute flood); however, the mutants that lack Fe-triggered germination phenotype in agar (i.e., *tt7* and *prx*) germinated regardless of the duration of flooding (Figure 7). Prolongation of flooding of soil reduced the time required to reach 50% germination by approximately 6.5 hours (61 hours in prolonged flooded soil vs. 67 hours in acute flooded soil).

**Figure 7:**
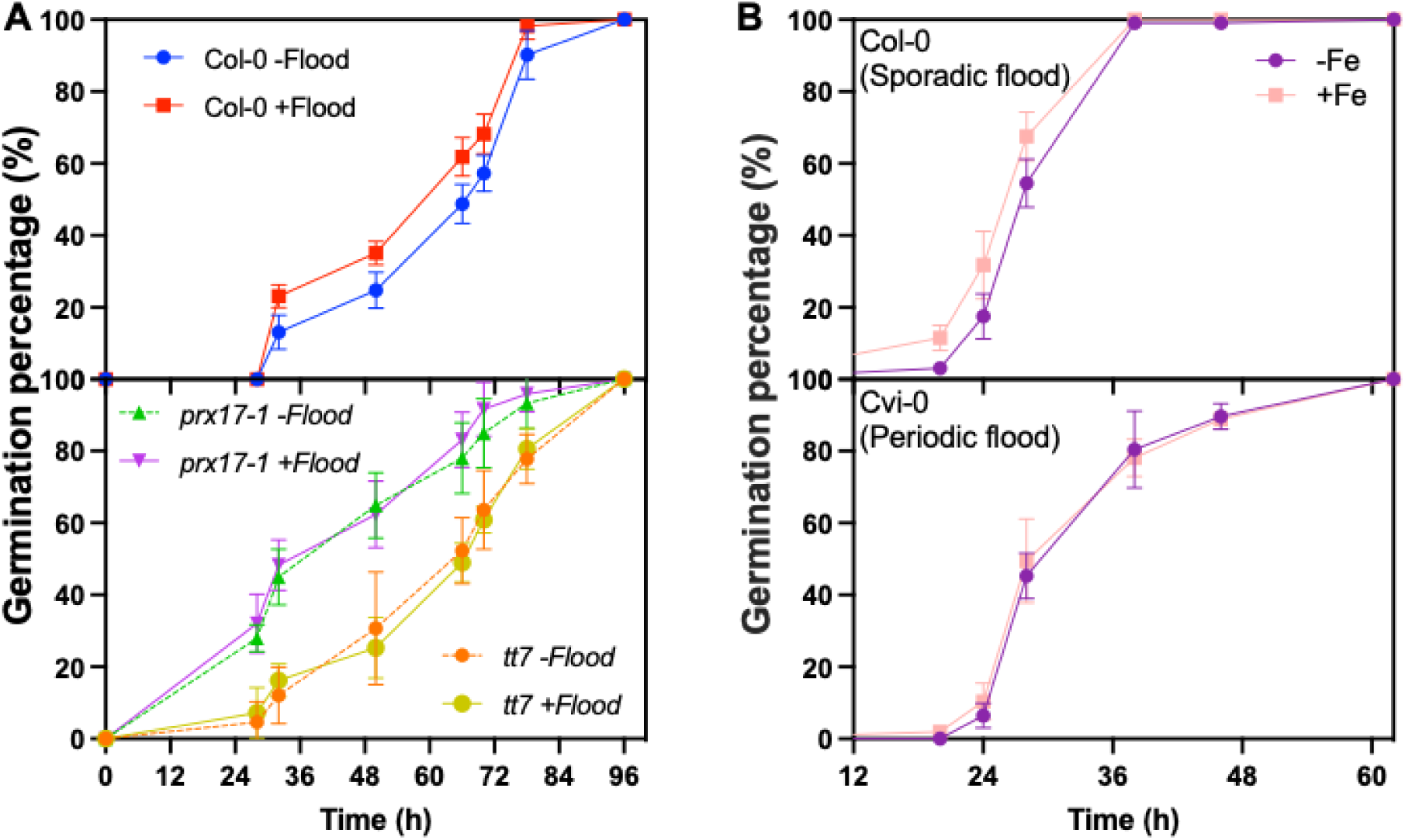
Fe-triggered germination is ecologically relevant in environments subject to sporadic flooding. A, germination time course of Col-0 and mutant seeds (*prx17-1* and *tt7*) on soil subjected to acute (-flood, 0 days) or prolonged (+flood, 3 days) flooding. Unfertilized soil was mixed with powdered dry plant material and incubated submerged in a growth chamber for 0 or 3 days. Seeds were placed on top of the sludge. Error bars represent ± SEM (n = 3 independent replicates of 30 seeds). Two-way ANOVA revealed a significant main effect of flood treatment only for Col-0 (*p* = 0.0116), whereas mutants remain insensitive to flooding. The experiment was conducted three times independently with similar results. B, germination time course of Col-0 and Cvi-0 seeds supplemented with or without 250 µM Fe-EDTA. Note that, compared to Col-0, which is adapted to a climate of continuous rainfall with sporadic floods, Cvi-0 is adapted to a climate with seasonal rainfall and continuous floods during the rainy season^4445^. The germination test was performed on agar medium. Error bars represent ± SEM (n = 3 independent replicates of 30 seeds). Two-way ANOVA revealed a significant main effect of Fe treatment only for Col-0 (*p* = 0.0113), whereas Cvi-0 remains insensitive to Fe supplementation. Experiment was conducted twice with similar results.

**Figure 8:**
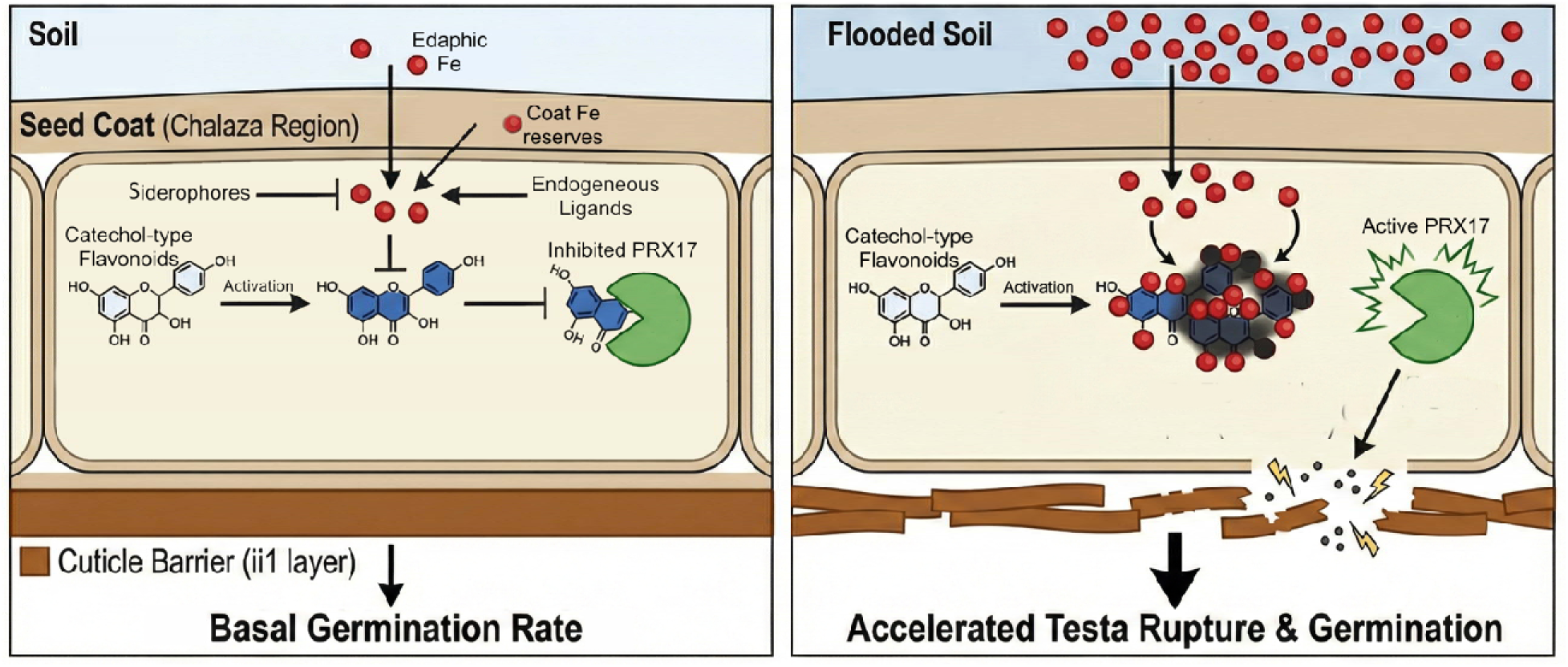
Proposed model for the Fe-sensing mechanism controlling germination timing. A, under low Fe availability (e.g., aerated soil), catechol-type flavonoids accumulate in the chalazal seed coat and bind PRX17 and inhibit its activity. The degree of inhibition is determined by the local balance of Fe, which is modulated by endogenous ligands (e.g., nicotianamine), maternal seed coat stores, and sequestration by microbial siderophores (e.g., desferoxamine). Consequently, the endosperm-associated cuticle (ii1 layer) remains intact, acting as a hydrophobic barrier that limits permeability and maintains a basal germination rate. B, under high Fe availability (e.g., flooding-induced hypoxia), solubilized Fe penetrates the seed coat and chelates the catechol-type flavonoids. This relieves the inhibition of PRX17 (derepression). The activated PRX17 catalyzes the remodeling of the cuticle barrier, increasing seed coat permeability and accelerating testa rupture to facilitate rapid seedling establishment upon receding water levels.

A mechanism that continuously represses germination may have evolved to take advantage of the post-flood environment, where floods occur sporadically. We hypothesized that such an adaptation is unnecessary where floods occur periodically. To test this hypothesis, we examined the presence of Fe-triggered germination phenotype in another Arabidopsis accession, Cvi-0. In the Cape Verde Islands, where Cvi-0 originates, precipitation is seasonal and limited between June and December^45^. Strikingly, Cvi-0 germination was not induced by Fe (Figure 7).

## Discussion

The precise timing of germination is a critical determinant of plant survival. A seed must remain dormant until environmental conditions favor establishment; yet once a promising window is detected, it must germinate rapidly to outcompete neighbors. Consequently, seeds have evolved sophisticated sensing mechanisms to recognize specific environmental cues, such as smoke-derived karrikins that signal a post-fire landscape, characterized by low competition and high light availability^46^. Sporadic floods, especially when prolonged, have been documented to damage existing vegetation and shift community composition through mechanisms including oxygen deprivation of the roots^47–49^. Here, we report a parallel sensing mechanism by which seeds detect flooding, which signals lower competition and continuous water availability during seedling establishment. This mechanism depends on hypoxia-induced Fe solubilization that occurs during flooding.

### PRX17-mediated cuticle remodeling controls seed permeability

Dry seed coats function as reservoirs for a diverse array of proteins, including numerous class III peroxidases, most of which remain uncharacterized^5^. Class III peroxidases have been associated with the *formation* of apoplastic barriers—hydrophobic layers, like lignin and suberin, that are deposited in the the cell wall^50^. For instance, PRX2 and PRX25 are known to catalyze the polymerization of aromatics to reinforce the seed coat, with their absence leading to increased permeability^14^.

By contrast, our data demonstrate that PRX17—a cell-wall localized peroxidase found throughout the developing seed coat and endosperm (Figure S8 and Figure S9)—functions in the remodeling of these barriers. While mature *prx17* mutant plants exhibit decreased lignin deposits and faster bolting^37^, its function in the seed appears fundamentally different. We identified PRX17 as source of the peroxidase activity observed on the endosperm surface during germination (Figure 4). PRX17 is active during early seed development in the seed coat (Figure S8, S9) and is reactivated in the endosperm upon imbibition^15^. Unlike the physical tearing typically associated with germination, this suggests a mechanism of biochemical remodelling. Critically, this process is occurring even when metabolism is suppressed by ABA (Figure S7), regulated post-translationally by Fe-flavonoid interplay (Figure 5, 6).

### The chalazal Fe hotspot acts as a permeability checkpoint

We observed a striking and highly reproducible spatial pattern in seeds prior to testa rupture: a defined “chalazal ring” and a distinct micropylar spot where Fe accumulation, peroxidase activity, and the characteristic Fe-flavonoid blackening colocalize (Figure 5 and Figure S10). This pattern emerges during the earliest phase of imbibition (water uptake), coinciding with the remobilization of Fe stores (Figure 1). Anatomically, these two hotspots correspond to the developmental openings of the seed : the micropyle (the entry point for the pollen tube) and the chalaza (the attachment point of the funiculus, the maternal umbilical cord). The seed coat cuticle is known to be discontinuous or modified at these poles^16,50^. Our data imply that cell wall constituents (i.e., flavonoids, PRX17, and Fe) interact with the solutes of the soil more directly in this region. Previously, these openings have been associated with key areas for water uptake^51^. Our data indicate that these are not passive open doors but act as a gatekeeping system regulated by the remodeling of the cuticle. Recently, in the phloem unloading zone in developing seeds, formation of a termination point that regulates nutrient delivery by modulating callose barriers was reported, with a striking resemblance to the shape of the chalazal ring we observed in mature seeds^52^. Exploring whether the phloem end of the developing seed indeed forms the chalazal ring in the mature seed, thus serving as a post-mortem gate, is a challenging task, given that well-defined seed coat layers during seed development collapse during seed maturation^53^.

### Bioconversion of coat flavonoids to sense environmental Fe

Flavonoids represent one of the most chemically diverse classes of plant metabolites, yet their specific functions in the seed coat have remained obscure. While seed coats accumulate both proanthocyanidins (PAs) and flavonols, their roles appear distinct: PAs are primarily associated with dormancy^54^ , and kaempferol has recently been linked to seed longevity^55^. Surprisingly, quercetin—often the most abundant flavonol in the seed coat and associated with UV shielding, ROS scavenging and pathogen defense^56^—has lacked a defined biological function in the seed coat until now.

Catechol-type flavonoids (including quercetin derivatives and possibly PAs) are stored as inactive forms^41^ and upon imbibition, are bioconverted into their active forms^57^ creating the distinctive blackening reaction we observed (Figure 5), which is specific to flavonoids containing a catechol moiety (a di-hydroxylated B-ring)^42^. The Fe-induced blackening reaction therefore serves as a tool to track deglycosylation. The conversion of quercitrin to quercetin most likely requires a rhamnosidase. While rhamnosidases are ubiquitous in bacteria and fungi, surprisingly a plant rhamnosidase has not yet been definitively identified^17,58^. Our functional data suggests that such an enzyme must be active in the imbibed seed coat, marking a critical target for future genetic identification.

Finally, we dissected the genetic requirements for this sensing mechanism using transparent testa (*tt*) mutants. The blackening reaction is completely abolished in *tt7* mutants. Since *TT7* encodes the enzyme F3’H (Flavonoid 3’-hydroxylase), its loss prevents the synthesis of both quercetin and downstream catechins, while leading to an overaccumulation of kaempferol. The absence of blackening in *tt7*, combined with our finding that *tt4* (total flavonoid null) seeds can regain the blackening response when supplemented *in situ* with quercitrin and Fe (Figure 5), argues in favor of Fe-quercetin interaction as the core mechanism. However, we note that due to the structural redundancy of flavonoids, catechins and others may also participate in this interplay.

### Cuticle remodeling links edaphic Fe levels to germination timing

Physiologically, a seed behaves like a sponge during imbibition, passively taking up water until saturation. However, once saturated, seeds lack a direct mechanism to assess the *amount* or *persistence* of the water in their vicinity. Such an information would be crucial for survival and fitness in which light rains falsely signal continuous water availability. We propose that the seed solves this problem by sensing edaphic Fe levels. In waterlogged or flooded soils, oxygen depletion drives the reduction of insoluble Fe(III) to soluble Fe(II) (Figure S12), a process often accelerated by soil microbes engaging in anoxic respiration, thereby elevating Fe to levels toxic to plants^43,44^. Thus, rising soluble Fe levels serve as a reliable chemical proxy for prolonged hydration or flooding. This allows the seed to fuel rapid germination immediately as water levels recede (Figure 7A). Consequently, we speculated that seeds may be unresponsive to Fe and minimally accumulate catechol-type flavonoids, such as quercetin, where predicting floods is not advantageous. Strikingly, Cvi-0, which originated from a climate characterized by periodic floods, contains the lowest amount of seed quercetin among the tested accessions^59^ and does not respond to Fe during germination (Figure 7B).

Our synchrotron X-ray fluorescence analysis reveals a striking spatial dichotomy: while the chalazal ring is Fe-rich, the remainder of the seed coat is remarkably Fe-poor (Figure 1). This pattern suggests that although the mother plant provisions enzymes and precursors throughout the entire coat, it restricts Fe availability at the surface. The seed coat functions as an incomplete circuit, strictly dependent on Fe supply to trigger widespread cuticle remodeling. We conclude that in Arabidopsis, the Fe-flavonoid-PRX17 module operates as a hybrid system: a conserved, maternally provisioned trigger in the chalaza, but functioning primarily as a monitoring system for flooding both in the chalaza and across the rest of the seed coat.

