## Supplemental table S1 for "Iron□Flavonoid□Peroxidase Signaling Triggers Seed□Coat Remodeling for Rapid Germination"

**Supplemental Figures**

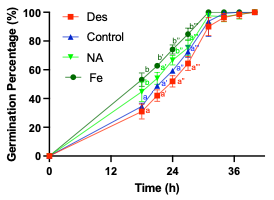

**Figure S1: Impact of Fe and Fe chelators on germination of *vit1*.** Germination time-course of vit1 seeds imbibed on filter paper buffered with MES (pH 5.5) containing nicotianamine (NA, 5 µM), desferoxamine (Des, 200 µM), or Fe-EDTA (250 µM). Statistics: Data represent means ±SEM (n = 4 replicates of 30 seeds). Letters indicate significant differences between the control and treatments (NA, Fe) at each time point; no significant difference was observed for Des treatment (Two-way ANOVA with Tukey’s post-hoc test, p ≤ 0.05).

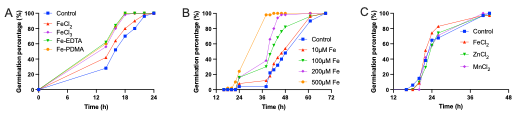

**Figure S2: Various Fe forms accelerate germination in a concentration-dependent manner.** A, Germination time-course of seeds imbibed with diverse Fe sources (FeCl_2_, FeCl_3_, Fe-EDTA, and Fe-PDMA) supplied at a concentration of 100 µM. B, Dose-response analysis of germination speed in the presence of increasing concentrations of Fe-EDTA (10–500 µM). C, Specificity of the response to Fe compared to other divalent cations. Germination kinetics were monitored in the presence of 100 µM FeCl_2_, ZnCl_2_, or MnCl_2_. In all experiments, germination percentages were calculated from a population of 50 seeds per treatment.

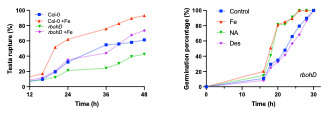

Figure S3: Fe mediated germination speed is not through RBOH driven hydroxyl radical production. Left, time-course of testa rupture in wild type (Col-0) and rbohD mutants imbibed with or without 250 µM Fe-EDTA (+Fe). Right, germination time-course of rbohD seeds in the presence of Fe availability modulators. Seeds were imbibed on filter paper wetted with Fe-EDTA (Fe, 250 µM), nicotianamine (NA, 10 µM), or Desferoxamine (Des, 200 µM). Germination and testa rupture were scored for at least 50 seeds per treatment.

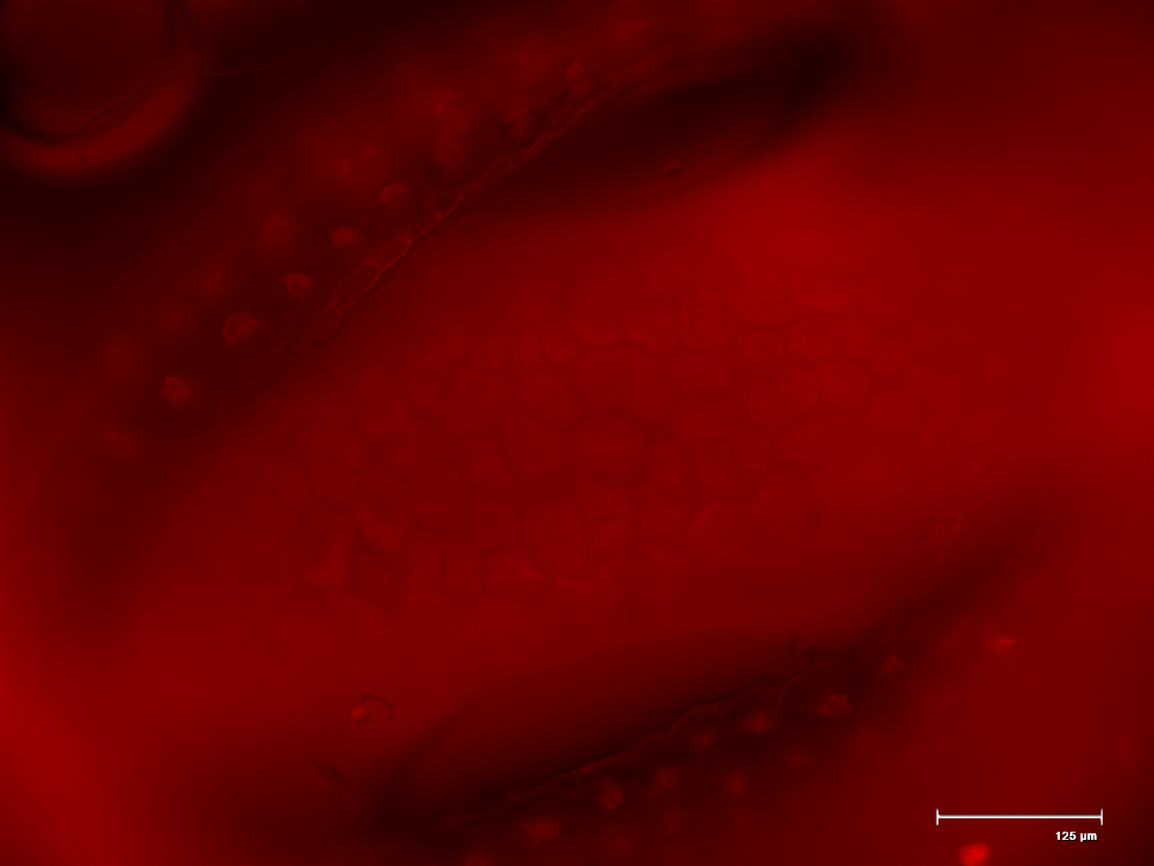

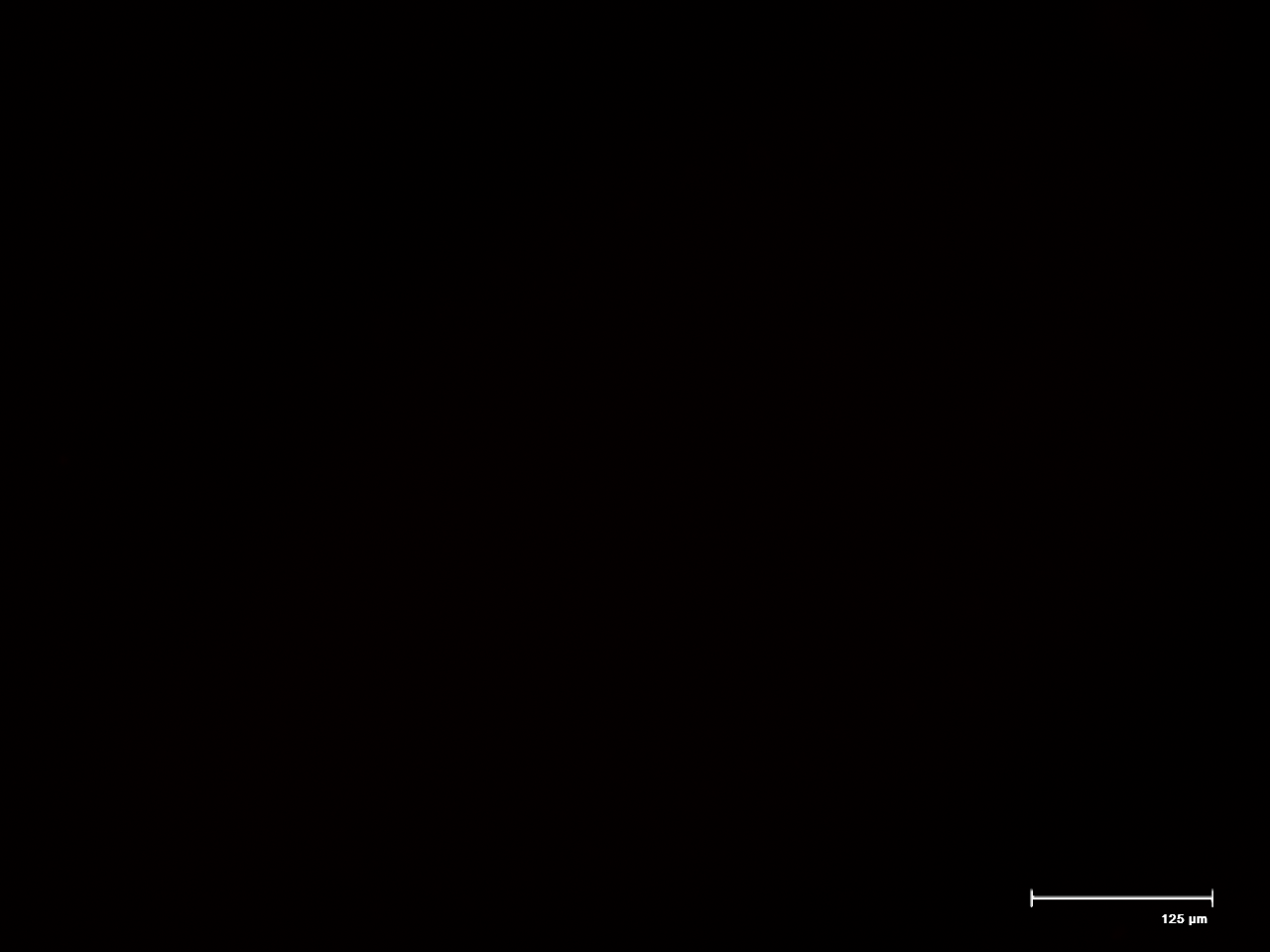

Nile Red (Suberin)

**C**

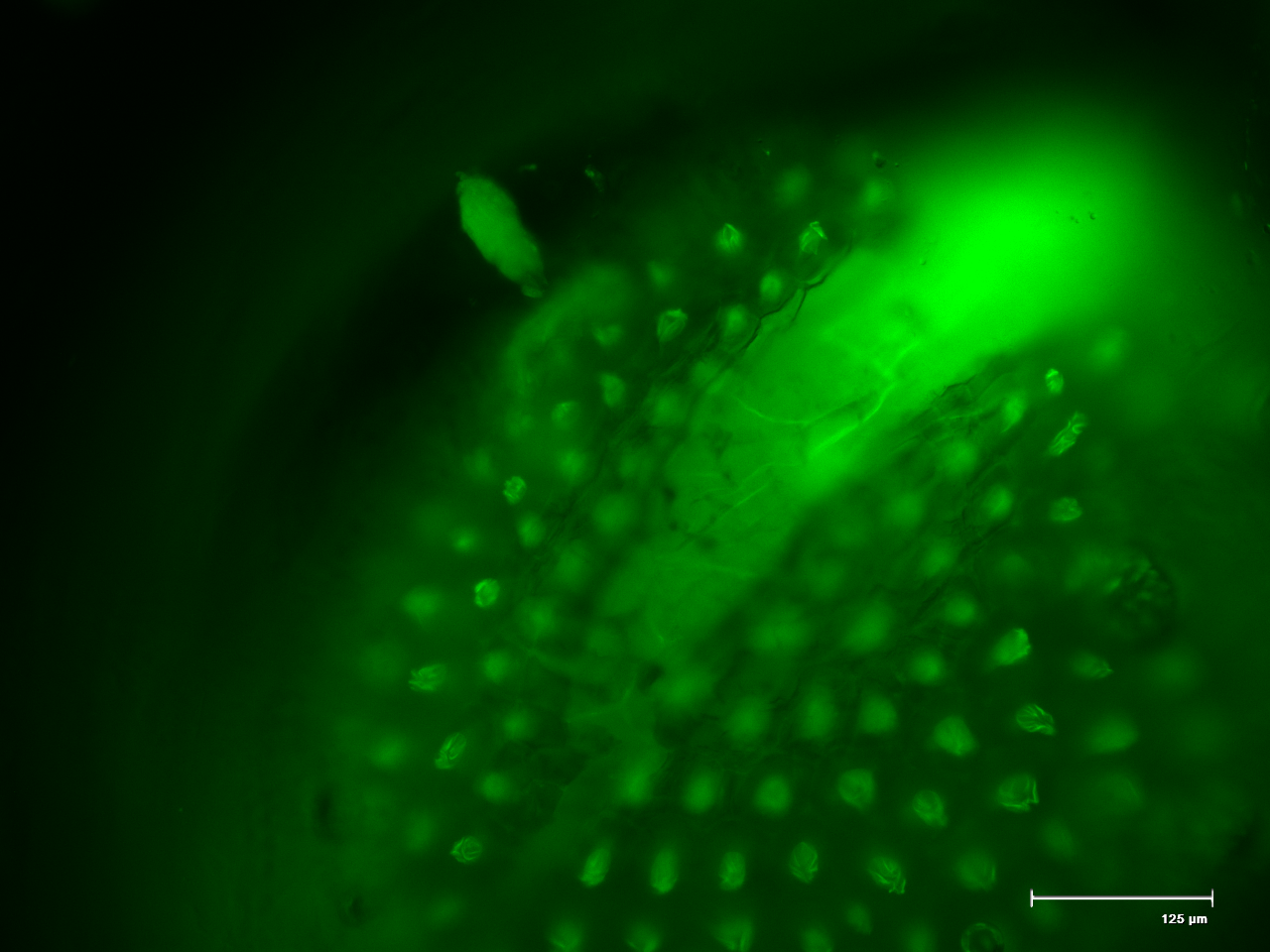

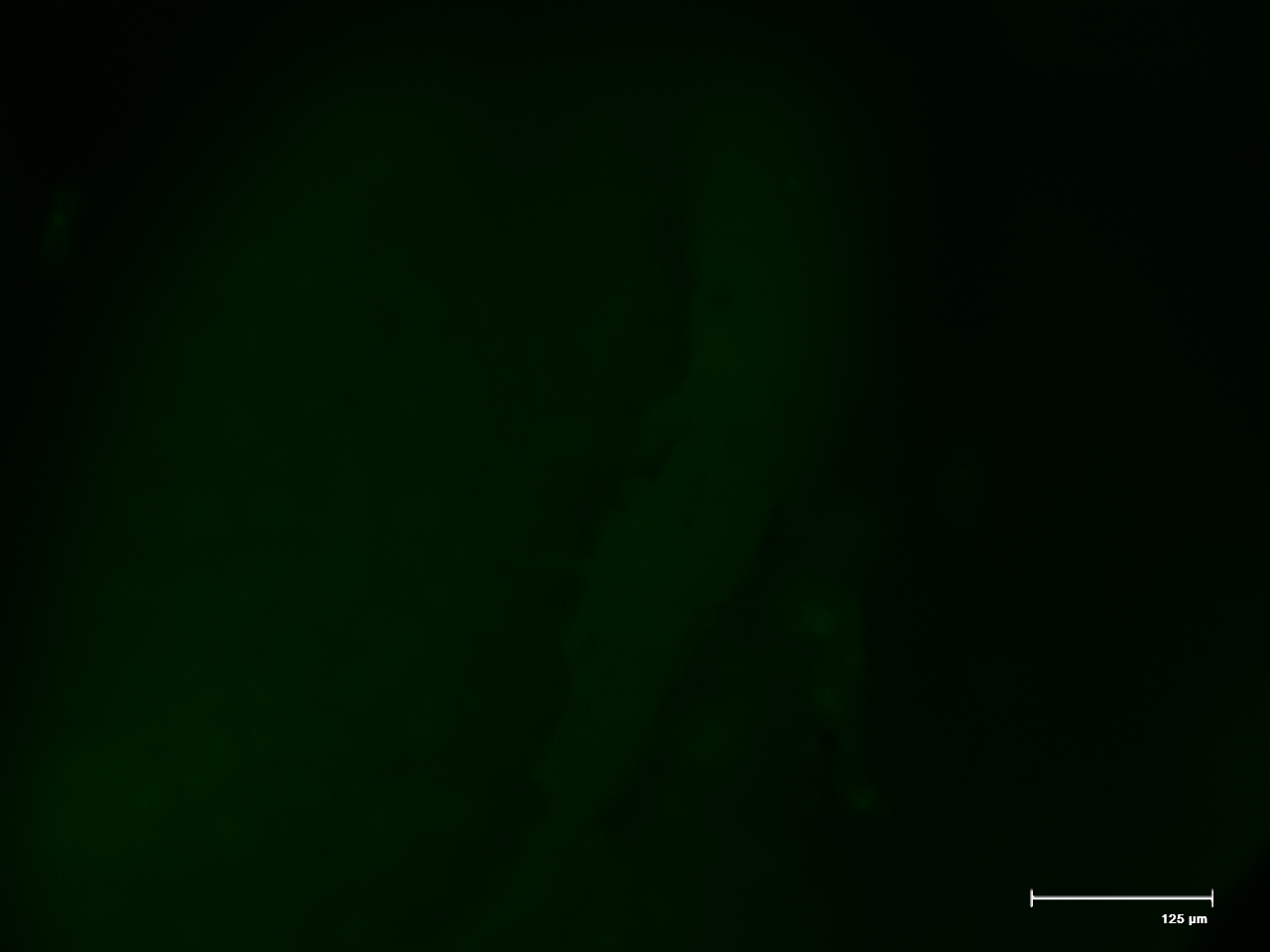

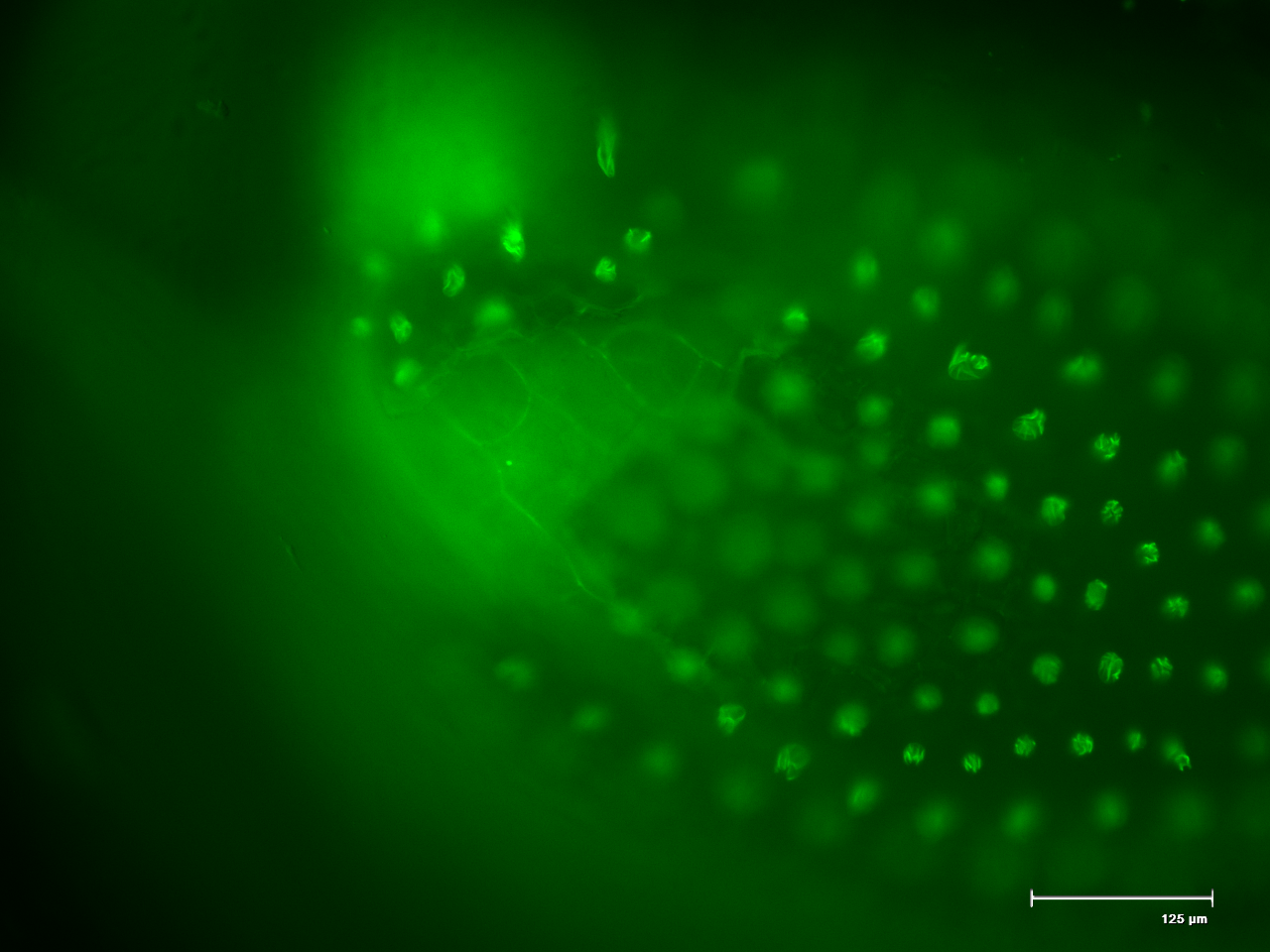

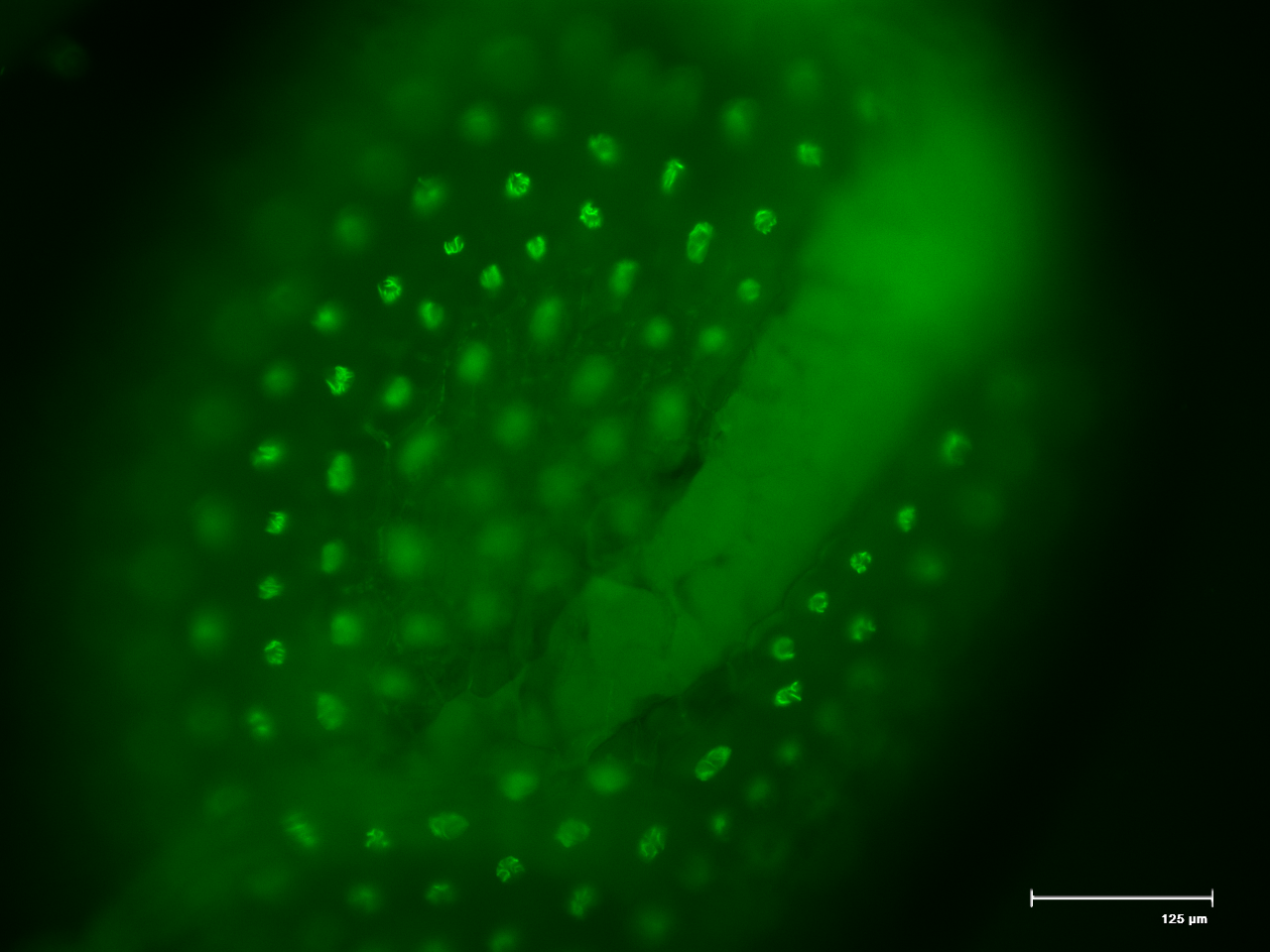

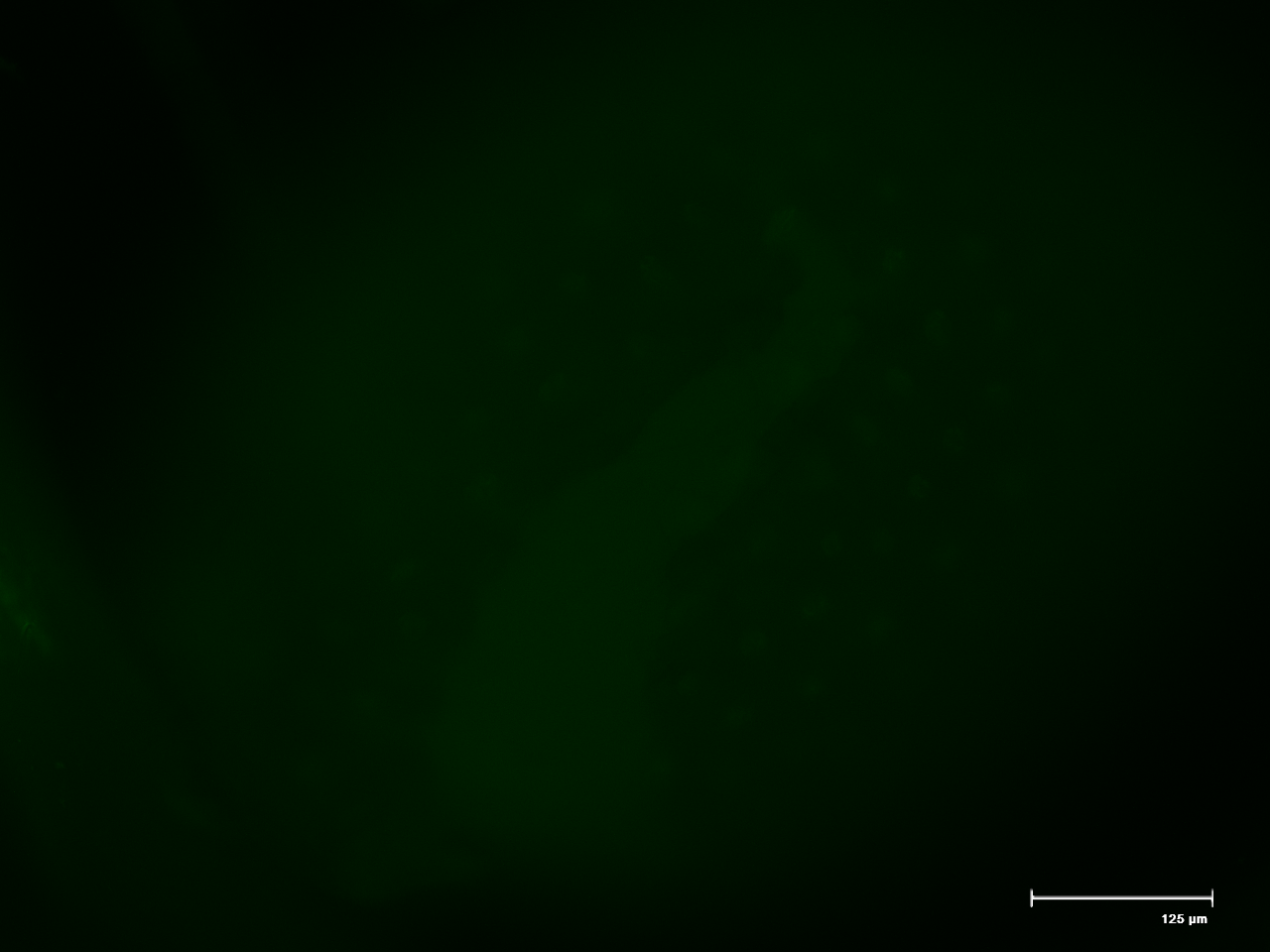

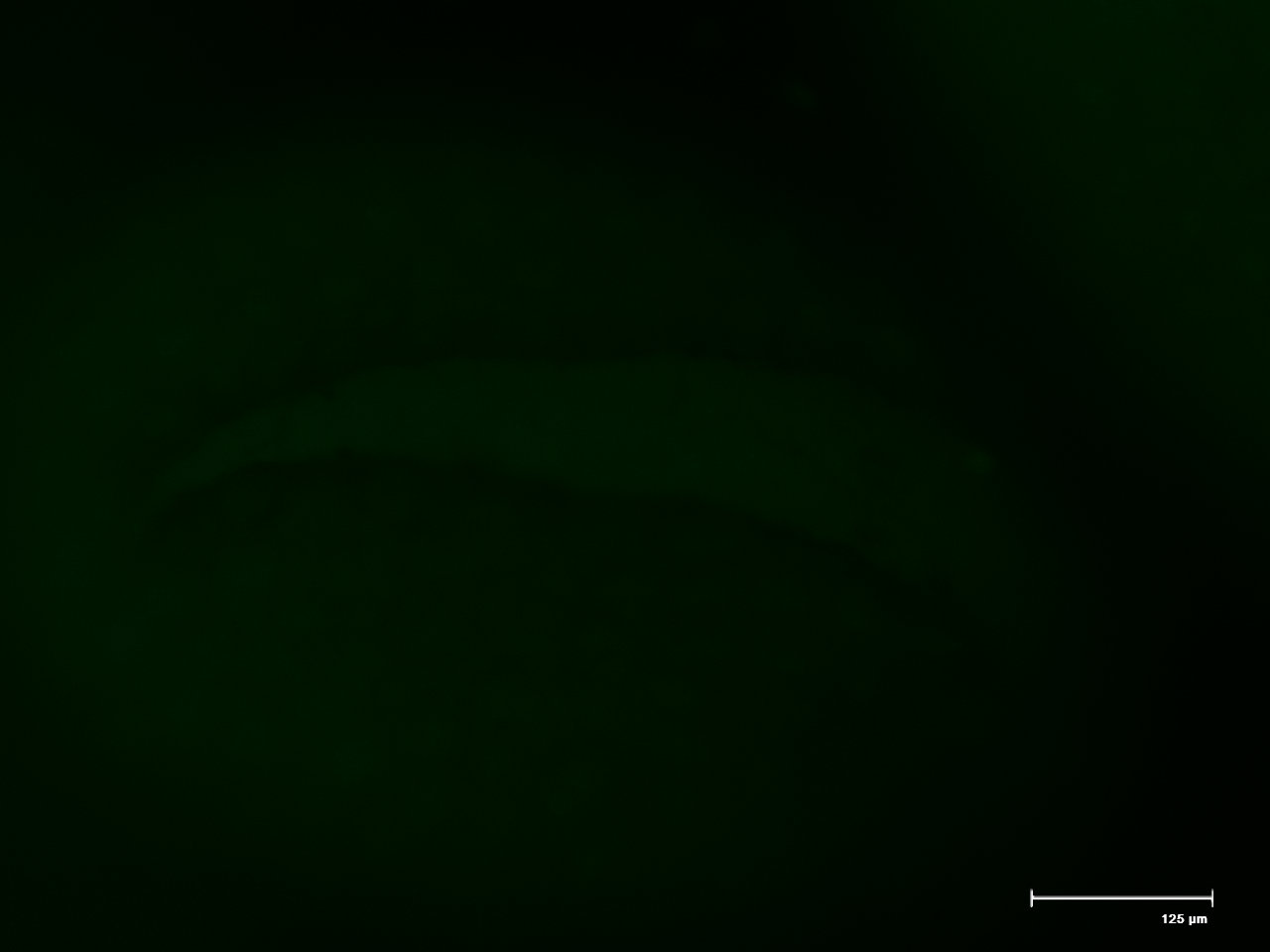

Auramine o

(Cutin, Lignin, Suberin)

-Fe	+Fe	+Fe+Met

**A**

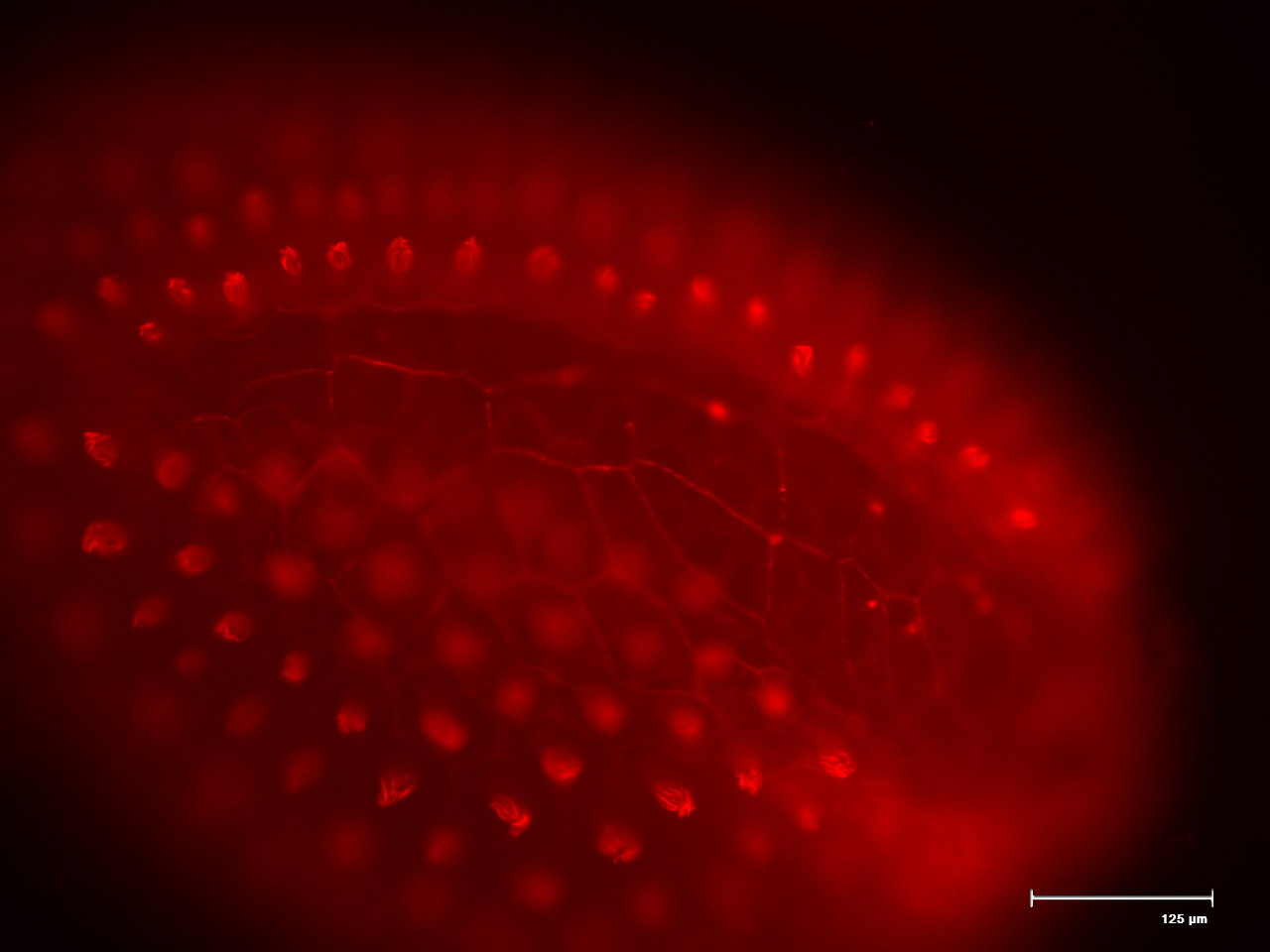

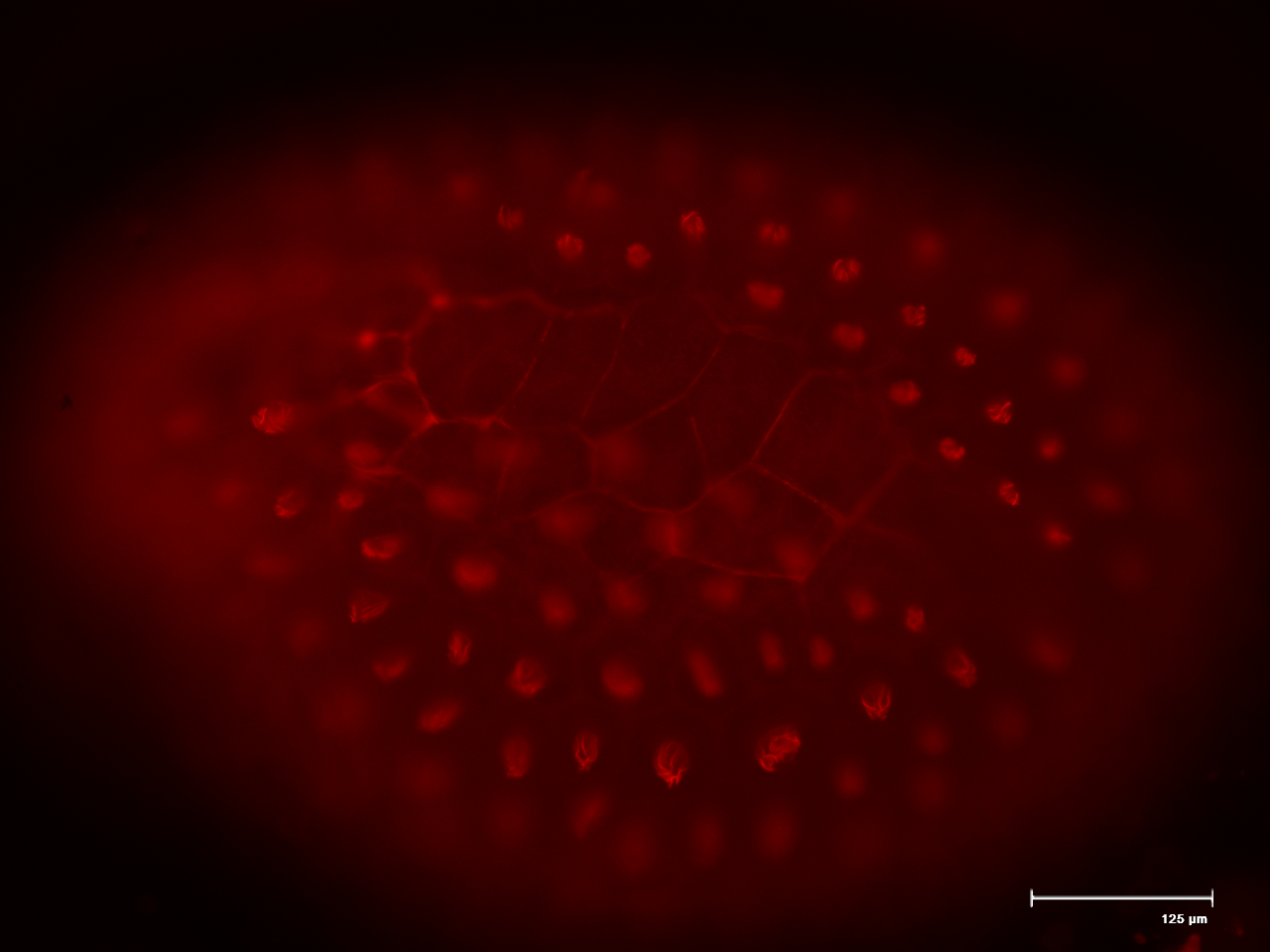

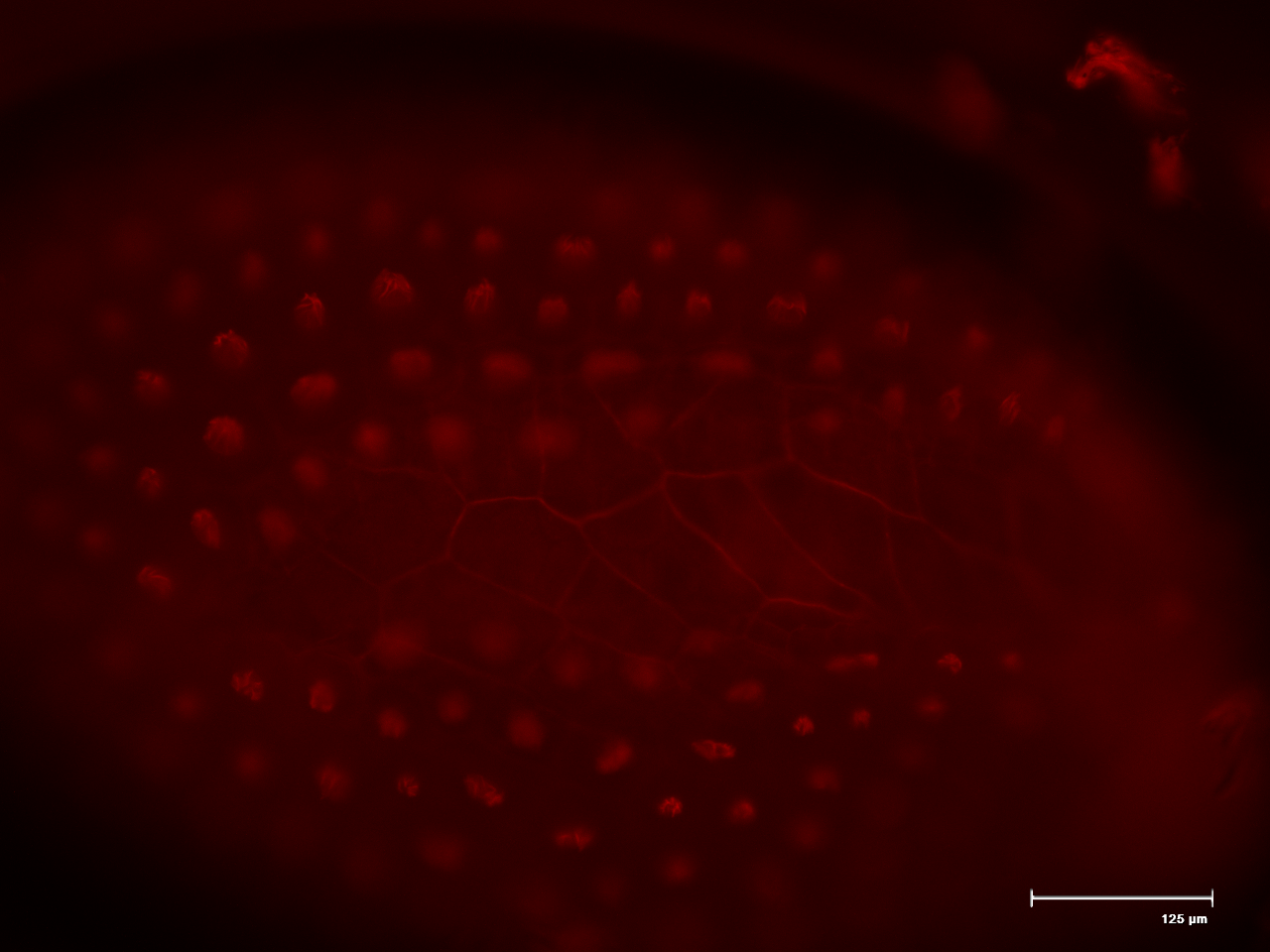

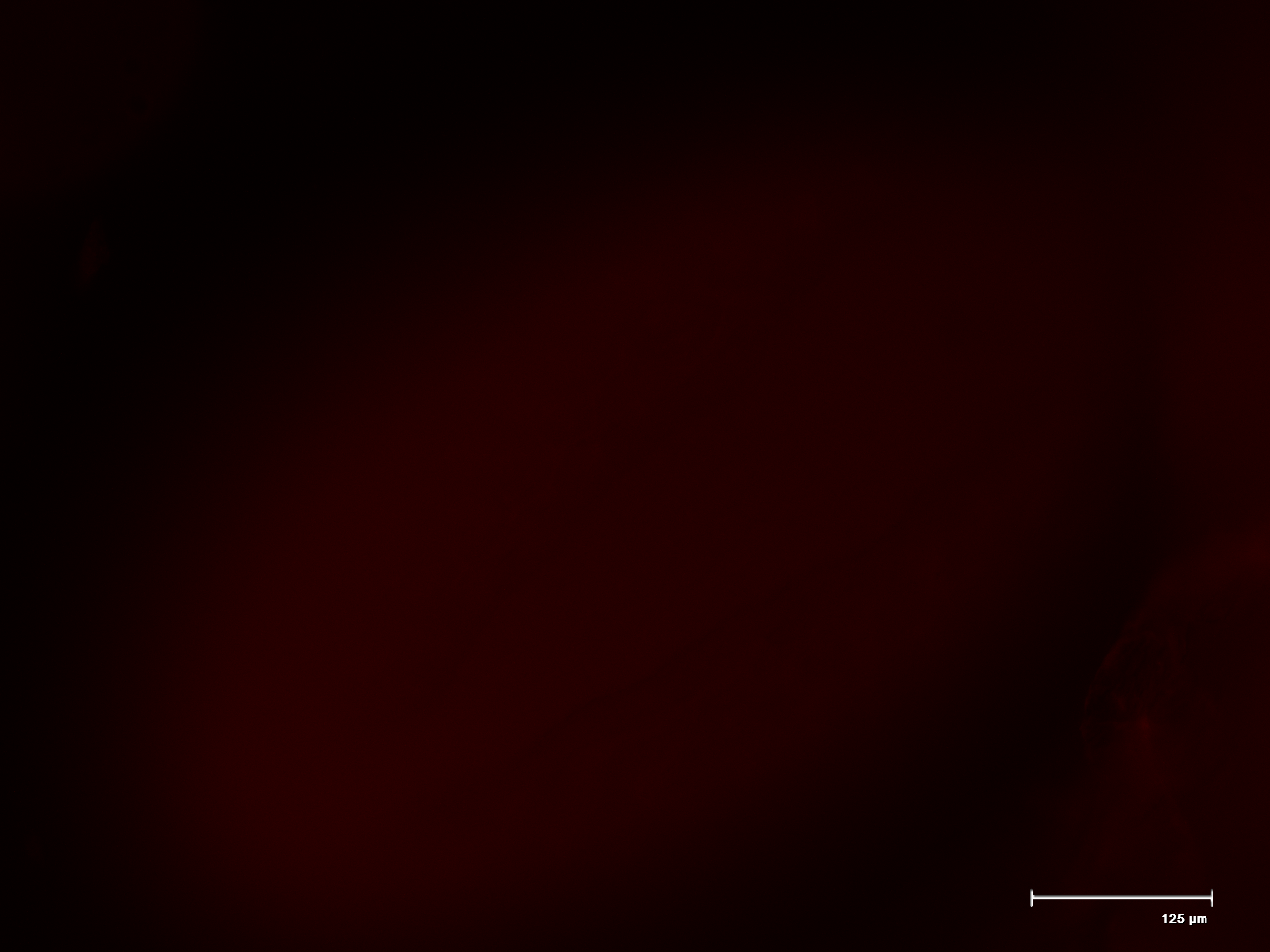

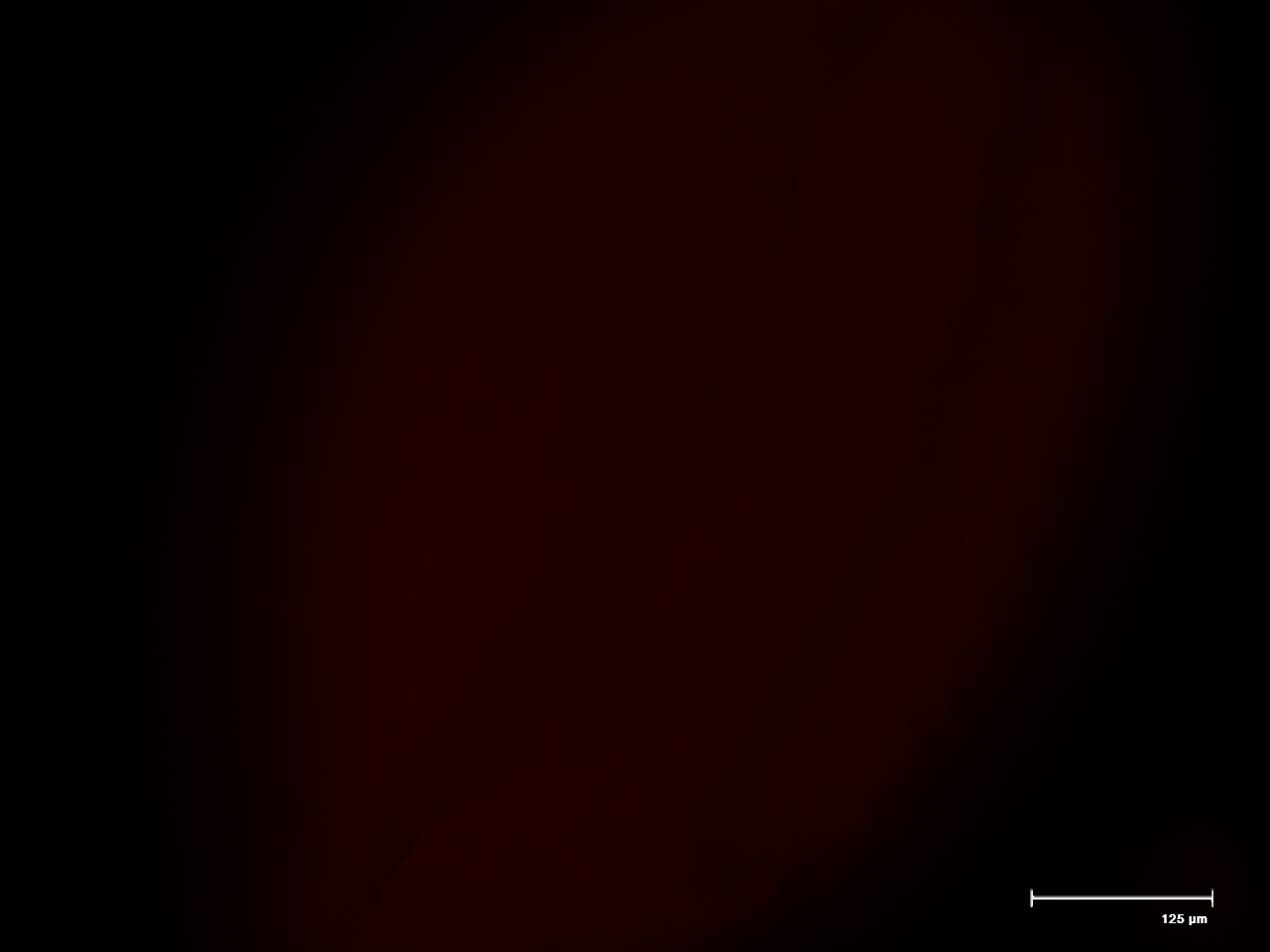

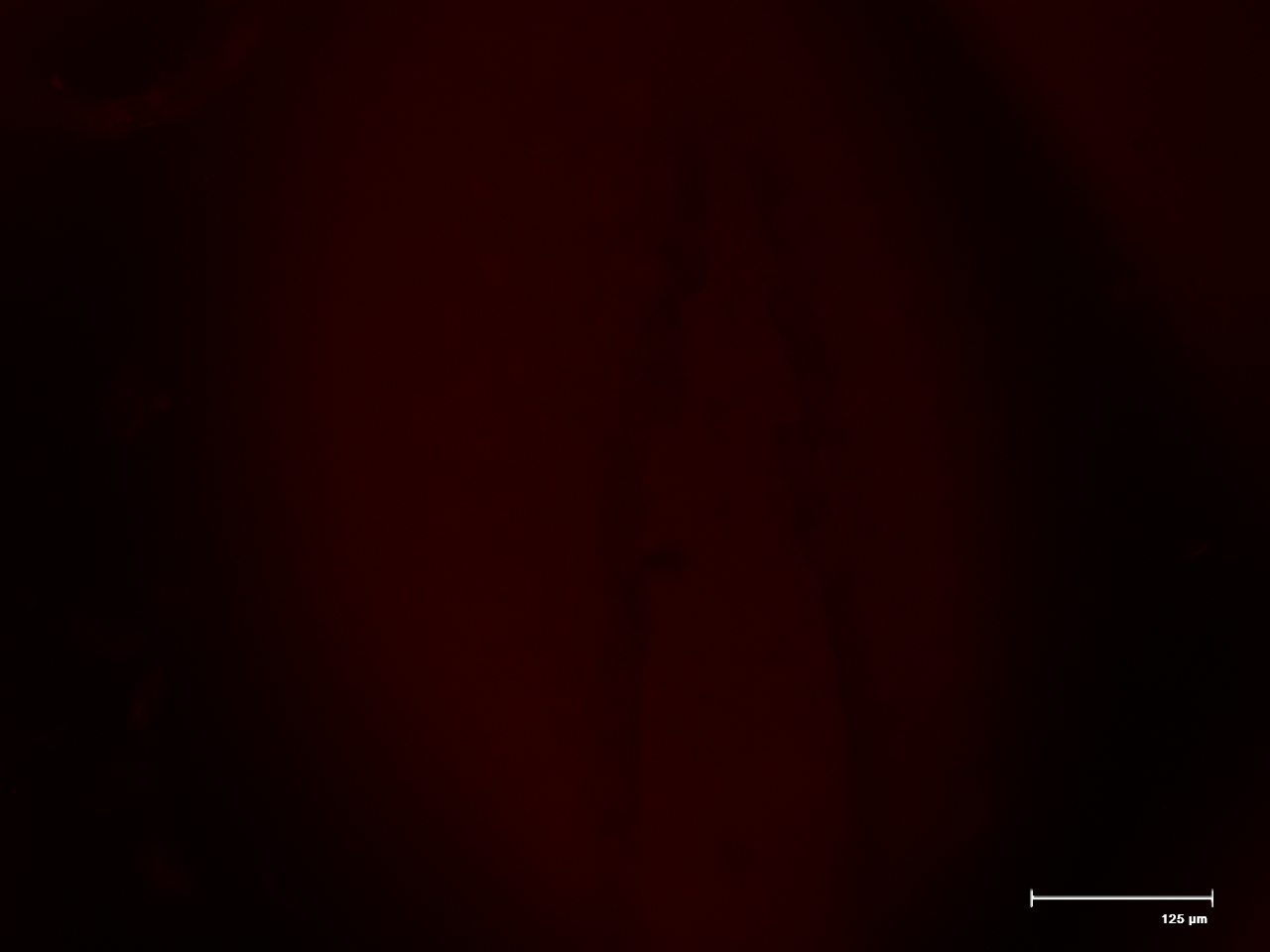

Basic Fuchsin (Lignin)

**B**

**Figure S4:** **The Fe-sensitive seed coat barrier is composed of cutin, not lignin or suberin.** A, epifluorescence microscopy of auramine O (general lipid/lignin stain) in testa-ruptured seeds. Seeds were germinated on 30 µM ABA, then transferred for 24 hours to media containing ABA alone (-Fe), 500 µM Fe-EDTA (+Fe), or 500 µM Fe-EDTA + 100 µM methimazole (+Fe+Met). White arrowheads indicate the continuous lipid barrier in control and Met-treated seeds, which is degraded in the presence of Fe. B, basic fuchsin (lignin-specific stain) signal in seeds treated as in (A). Note that the lignin signal remains intact across all treatments, indicating that the barrier degraded by Fe is not lignin. C, nile red (suberin-specific stain) signal in control seeds. The absence of a defined barrier signal suggests suberin is not a major component of this layer. Insets: No-stain controls showing background autofluorescence. Sample size: Representative images selected from 3–4 independent biological replicates.

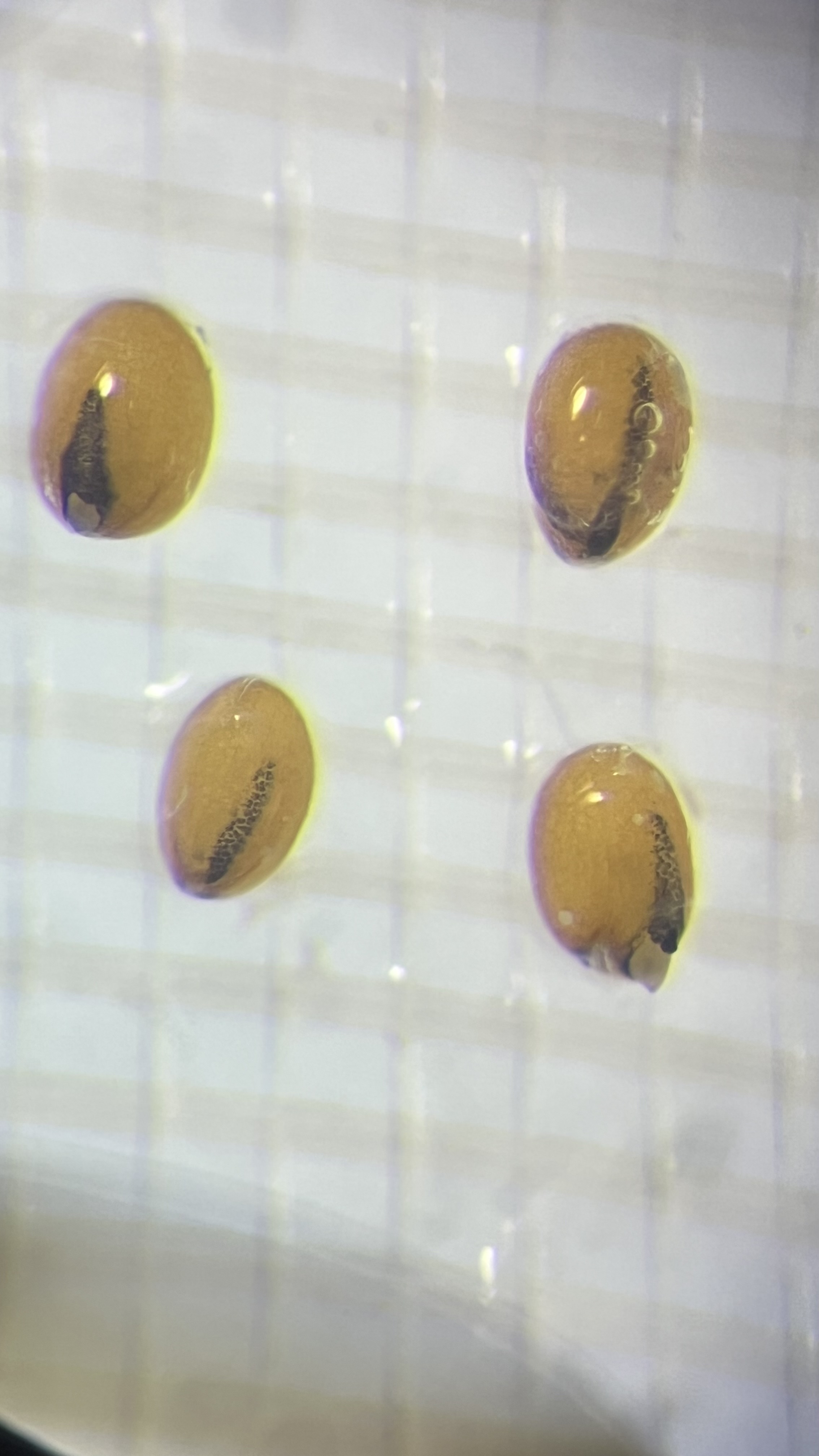

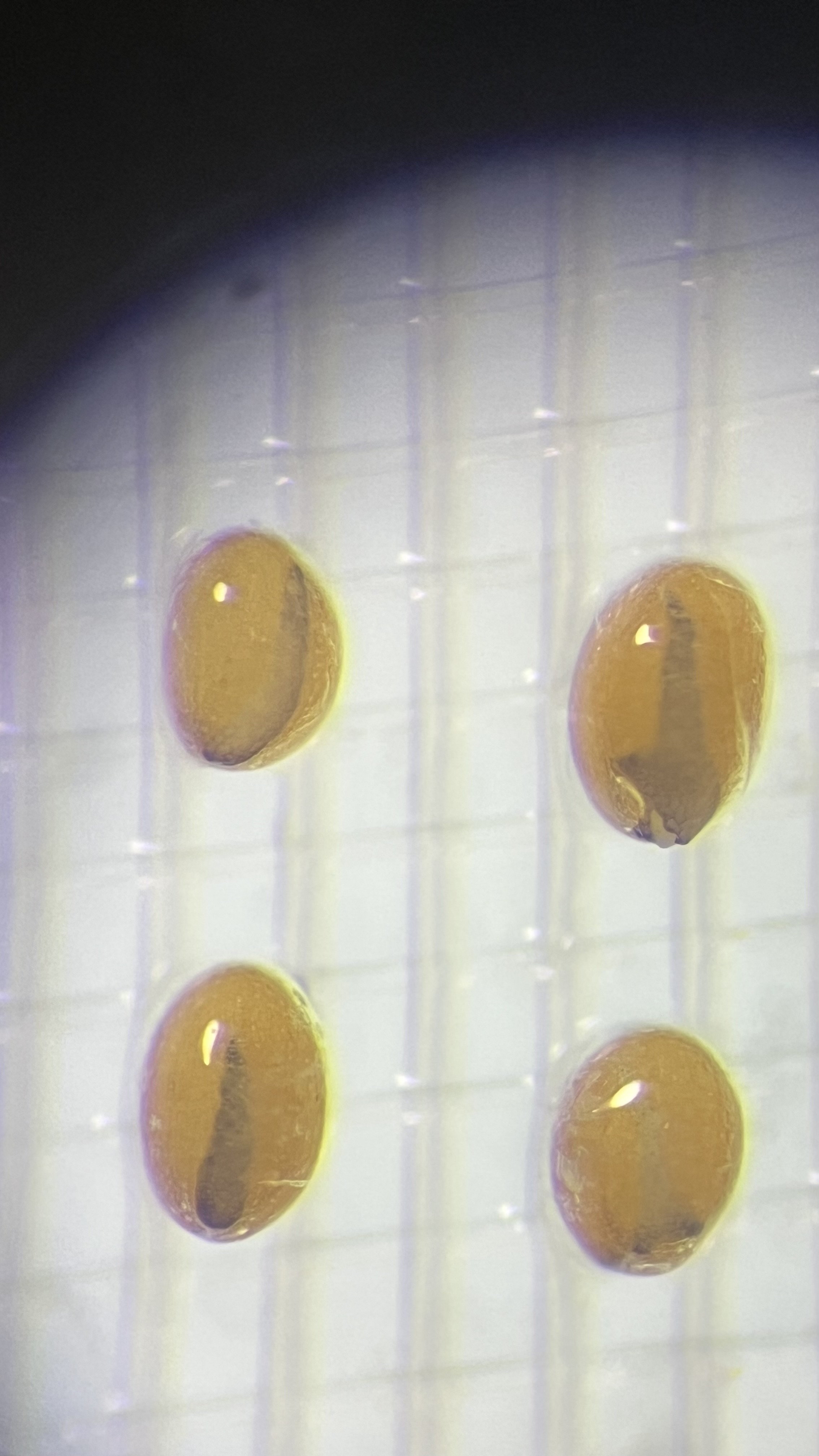

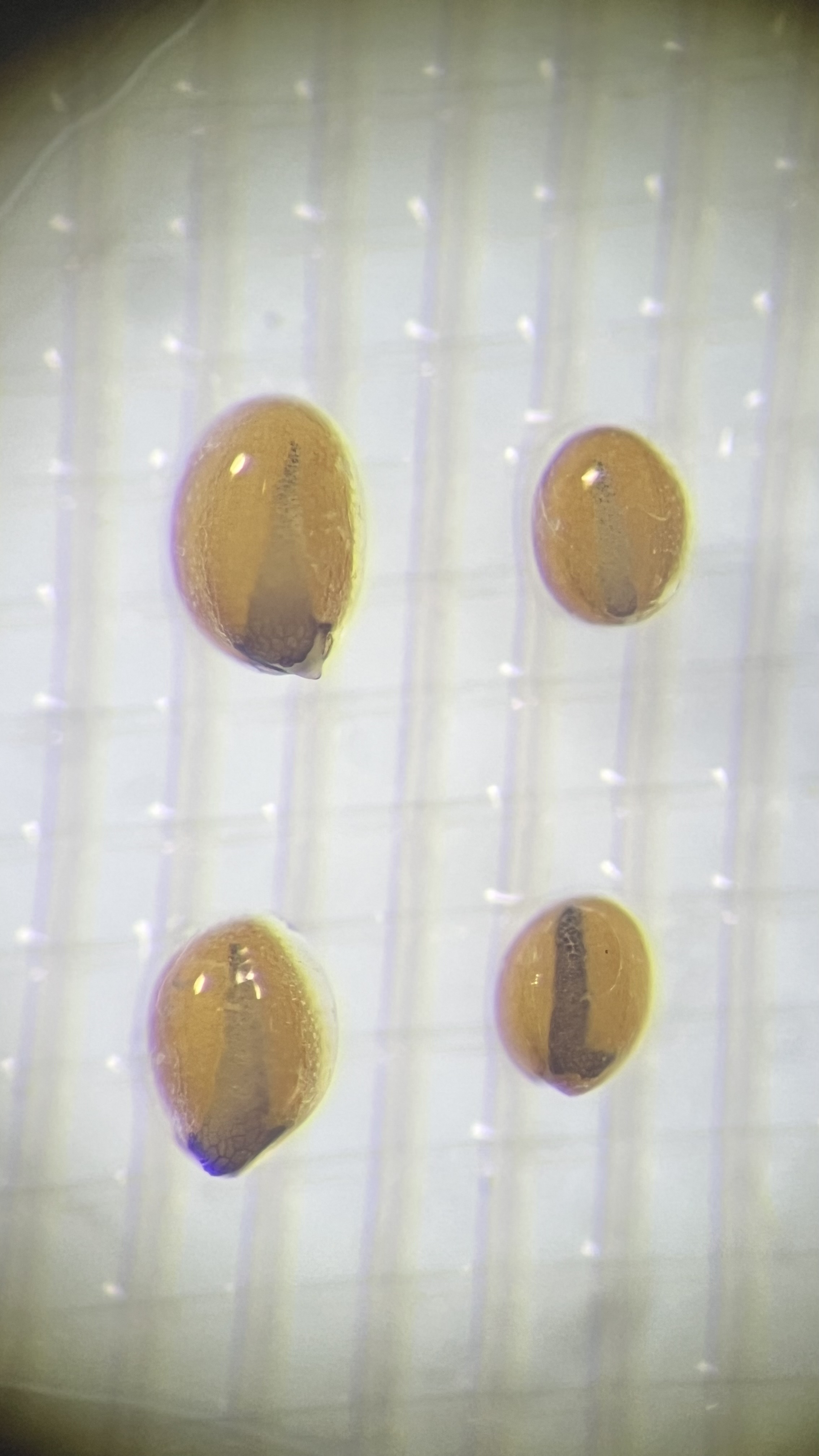

Col-0

*prx17-1	prx17-2*

Figure S5: Fe-induced peroxidase activity is suppressed in *prx17* mutants. Seeds were imbibed for 24 hours on filter paper saturated with 250 µM Fe-EDTA (without ABA). Peroxidase activity was visualized using chloronaphthol staining. Representative images were selected from at least 10 examined seeds per genotype.

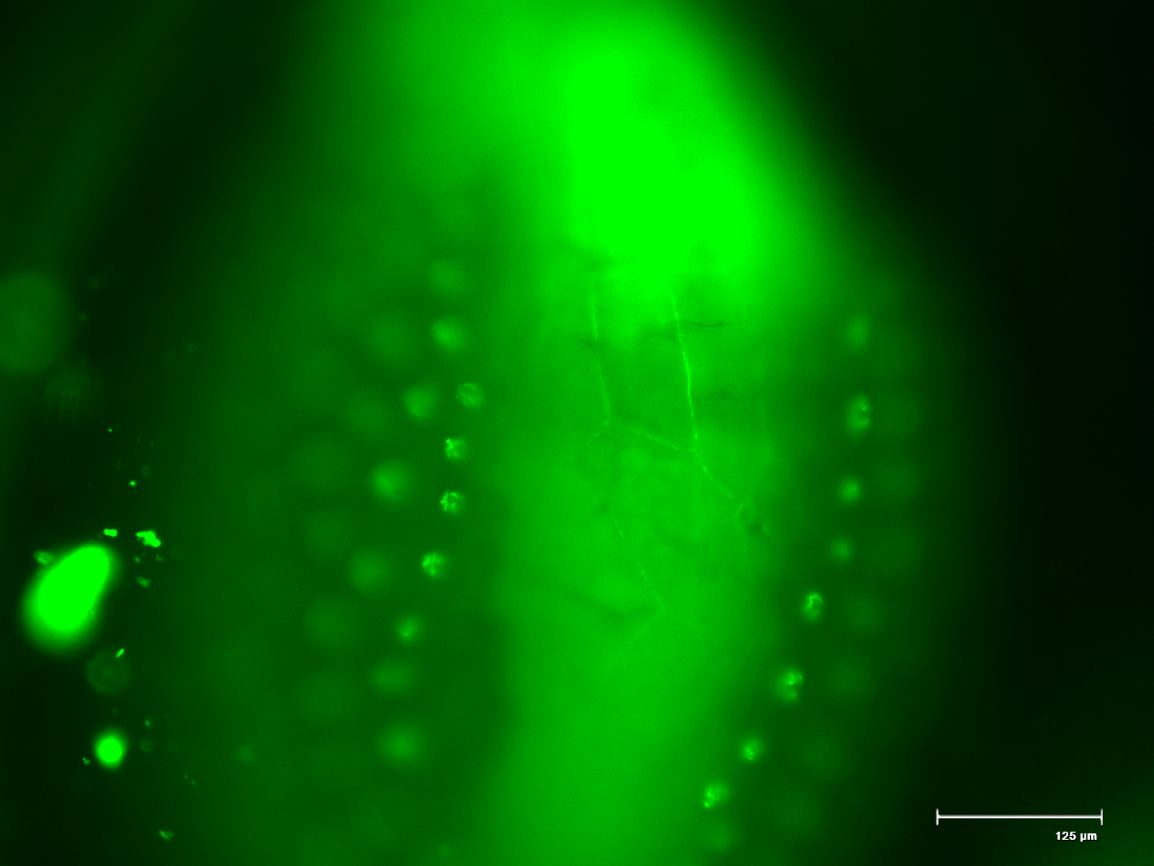

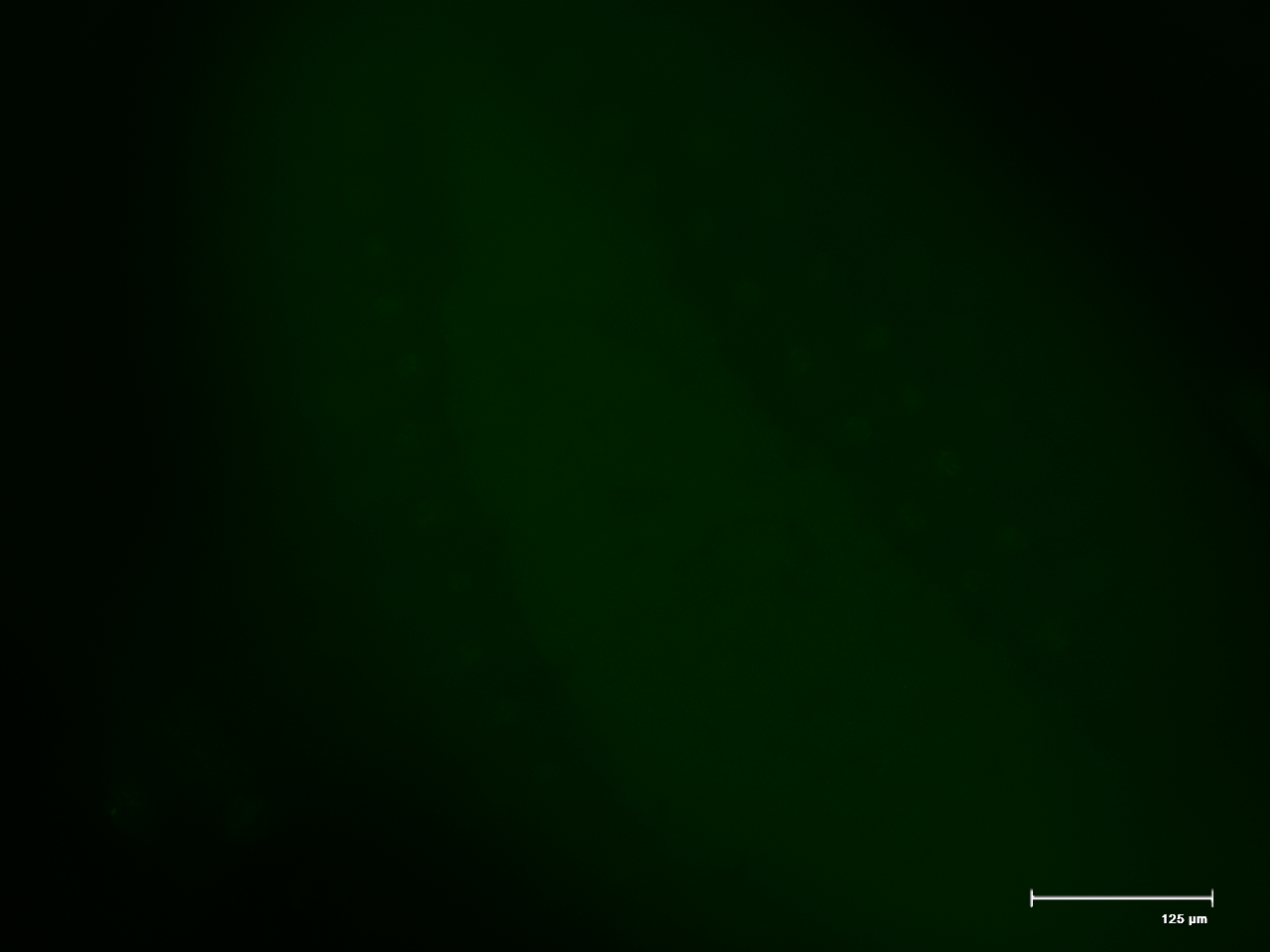

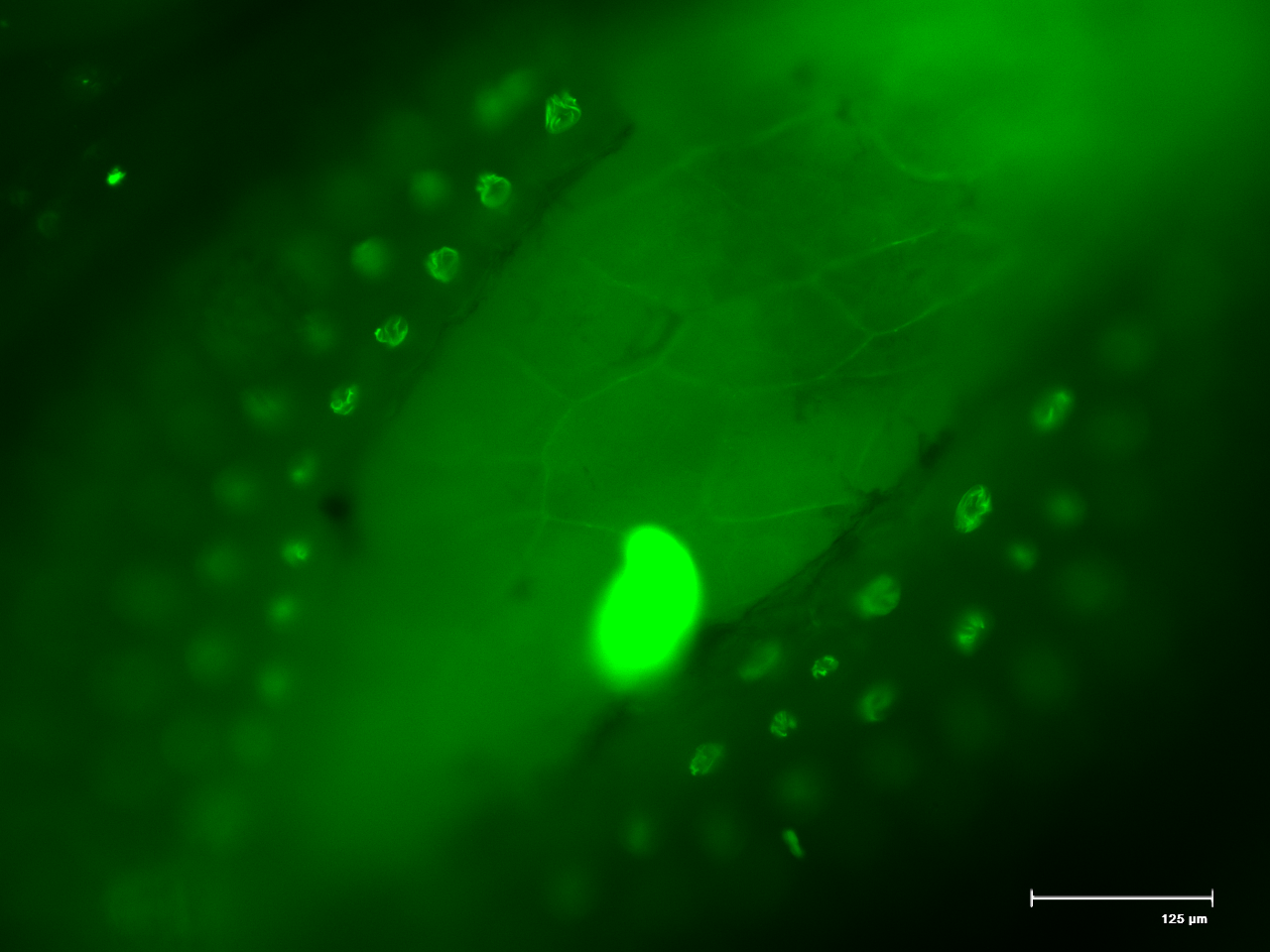

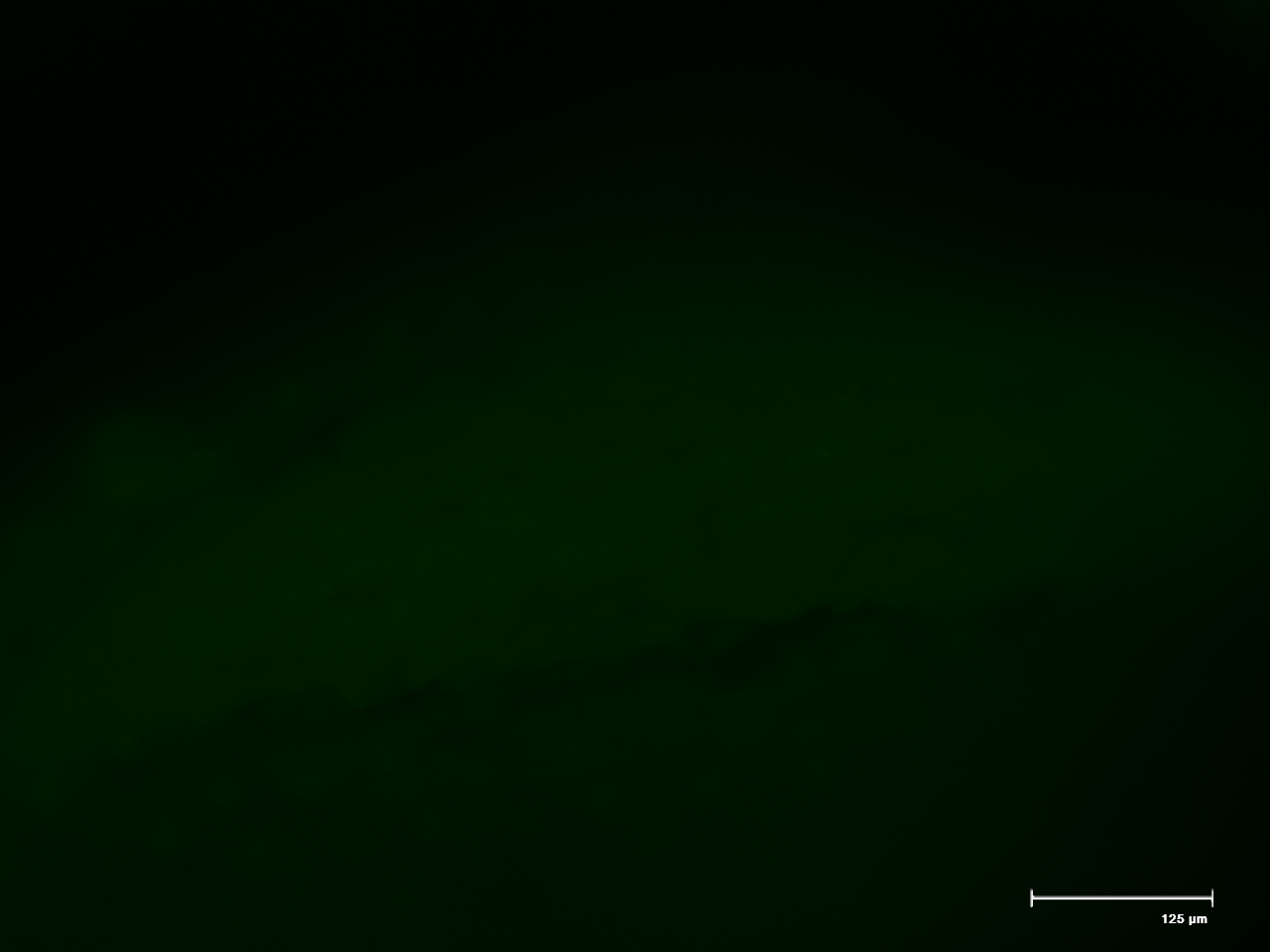

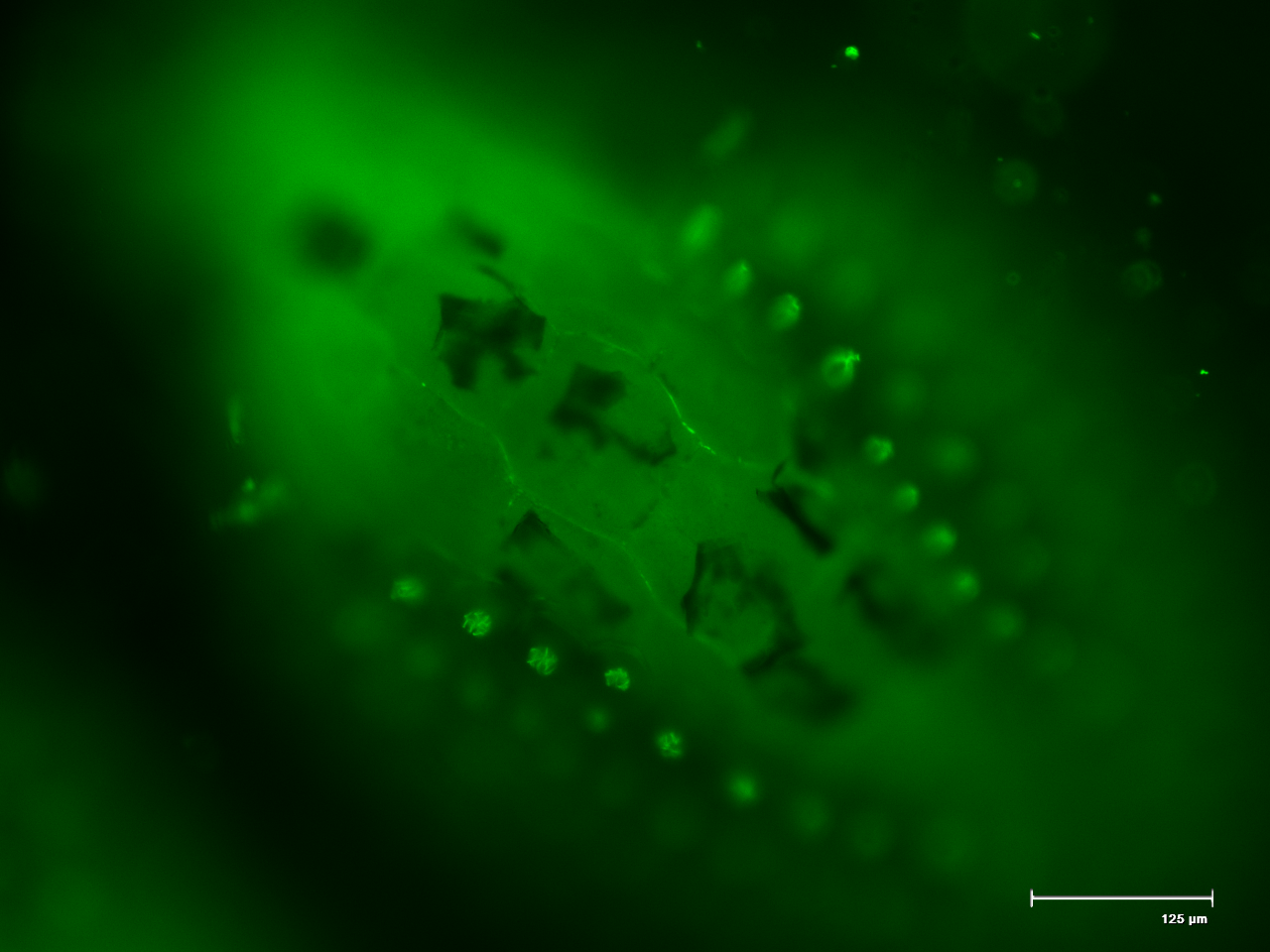

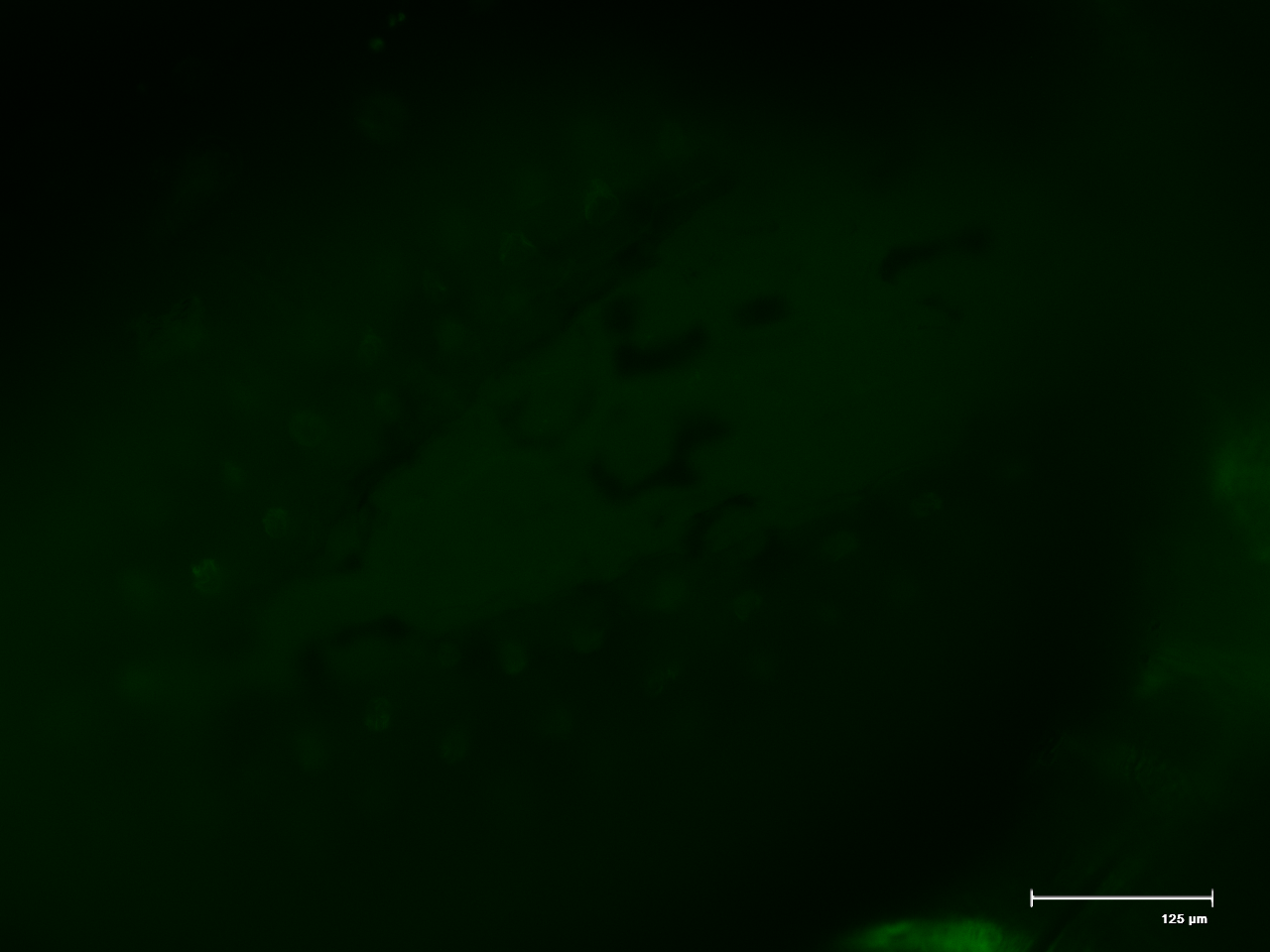

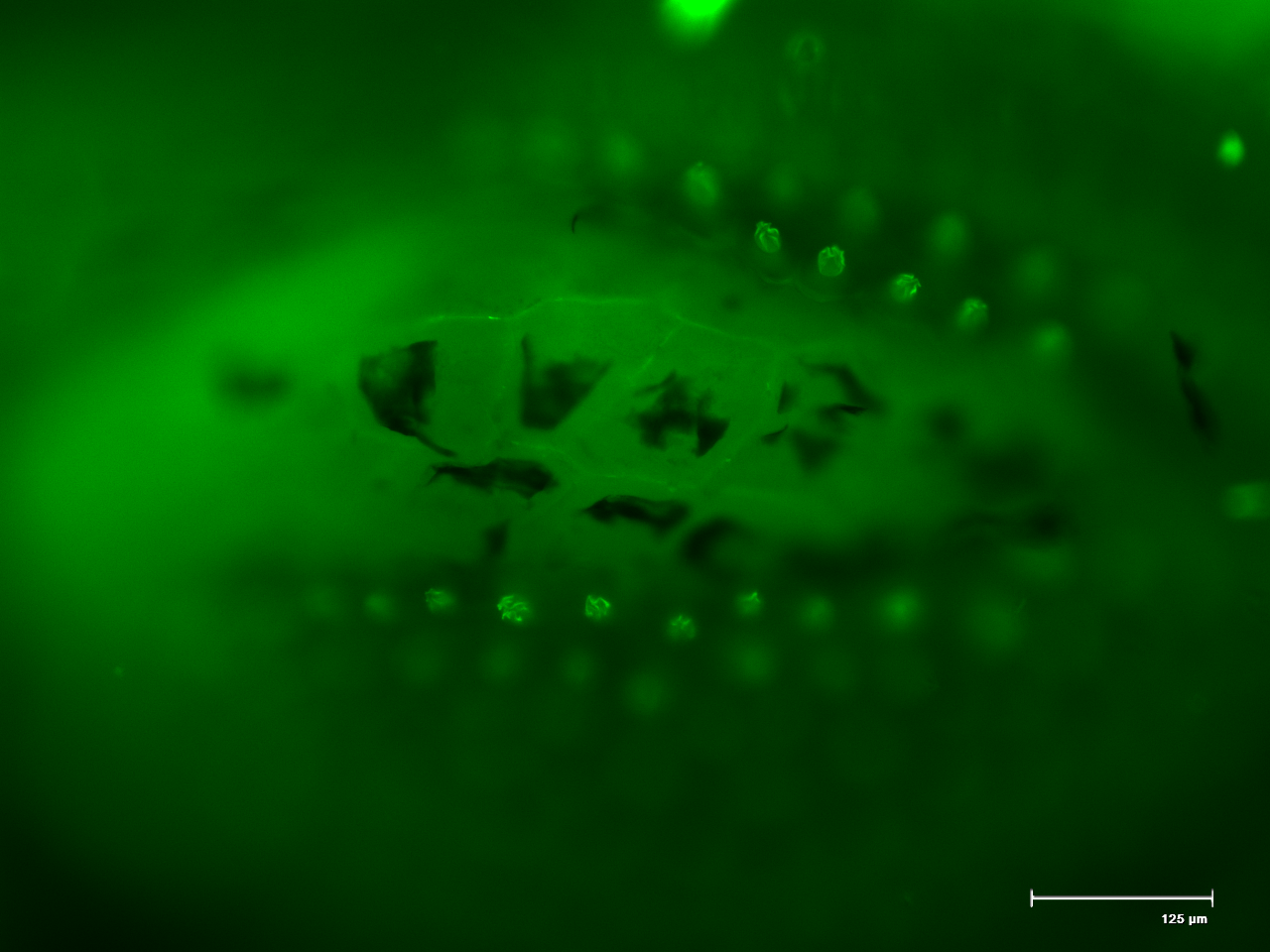

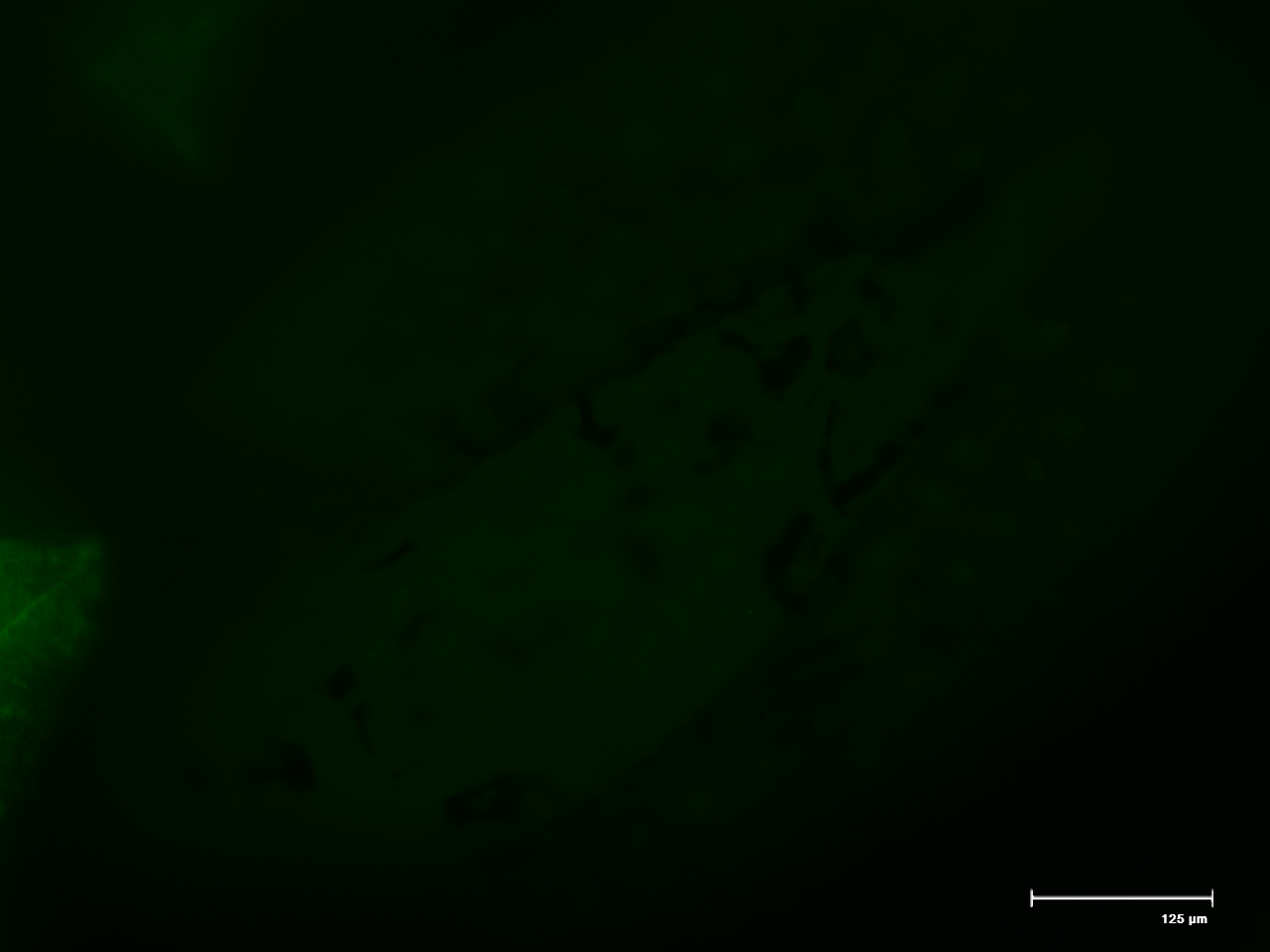

*prx17-1*

*prx17-2*

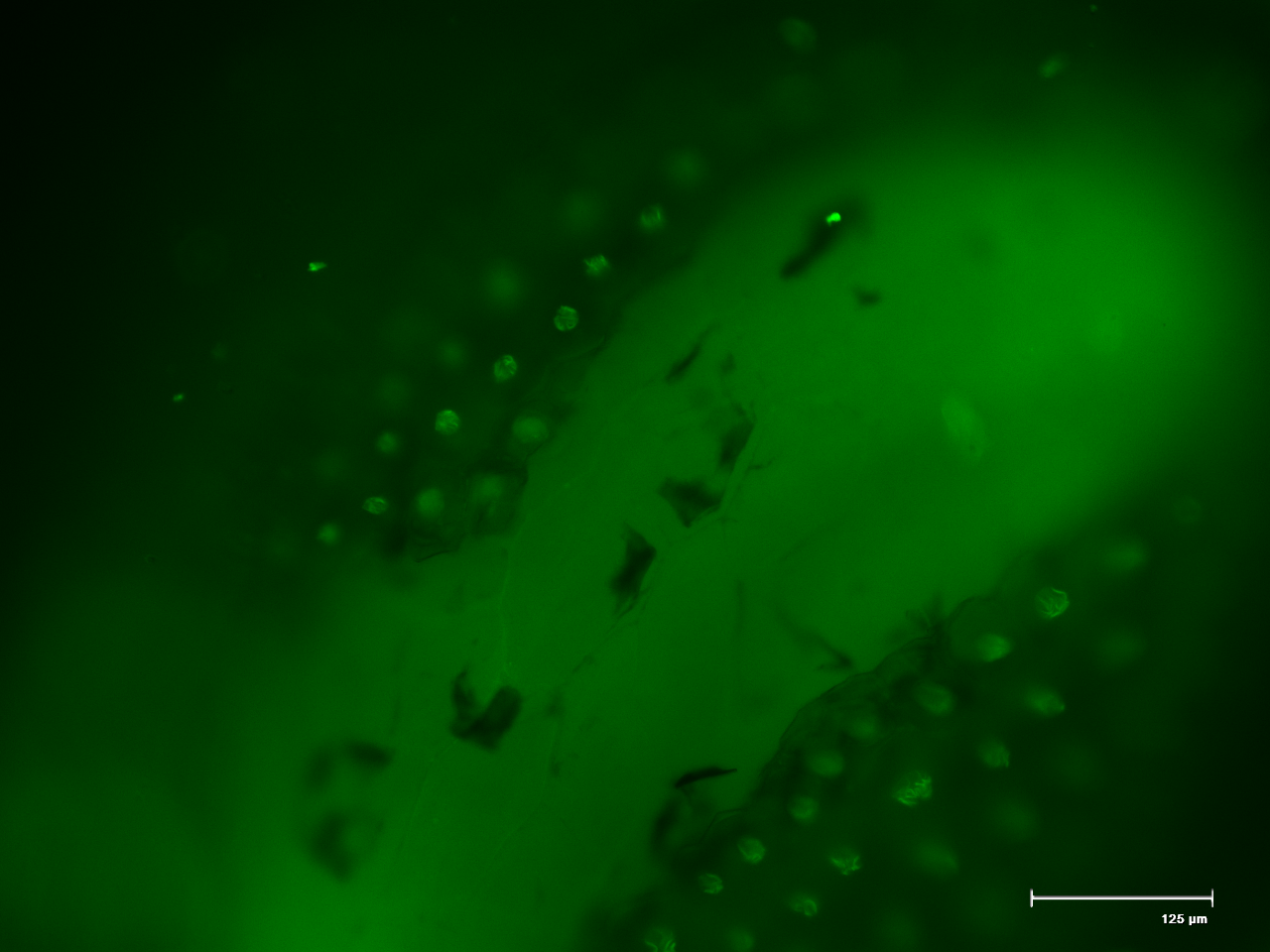

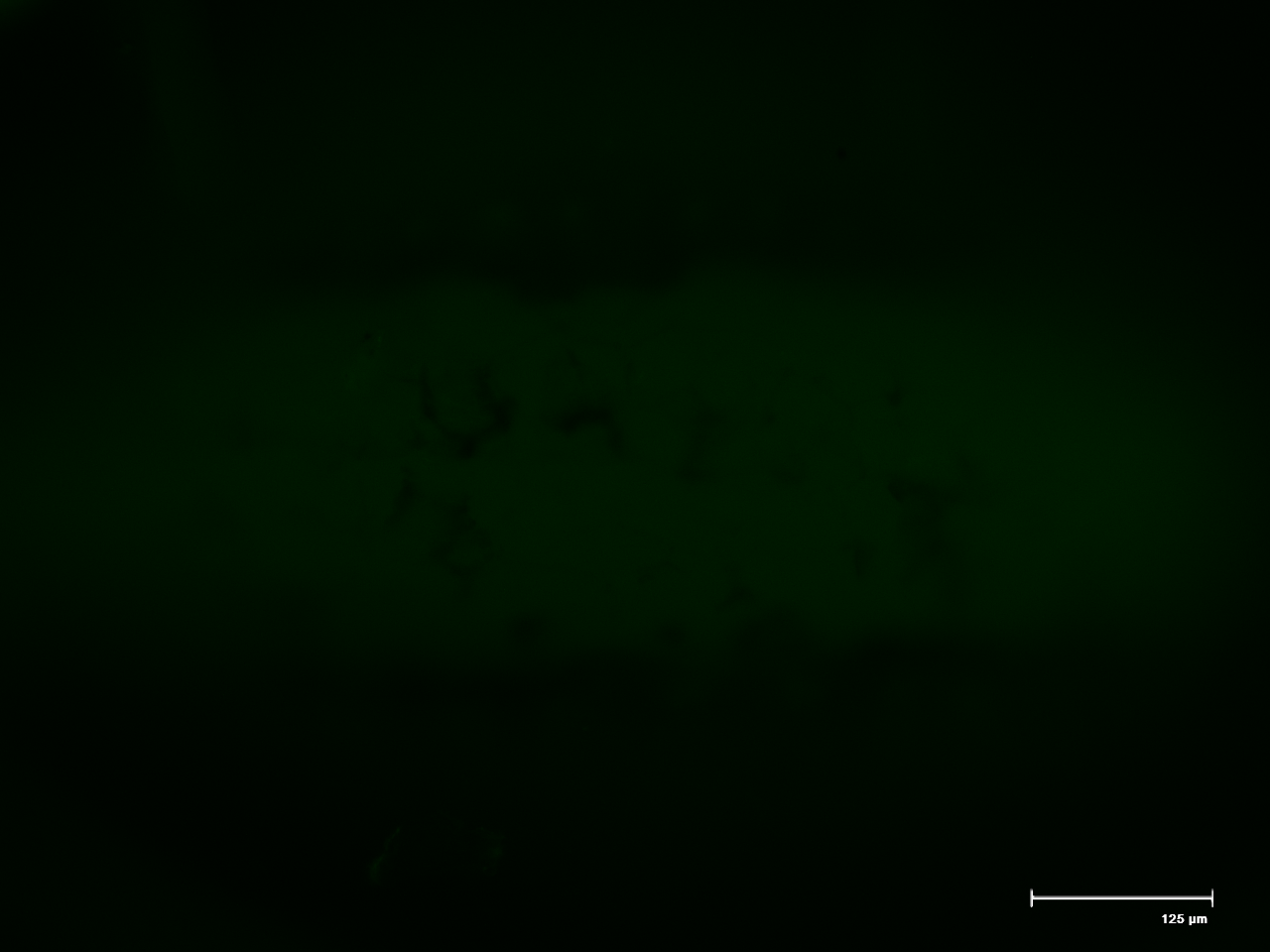

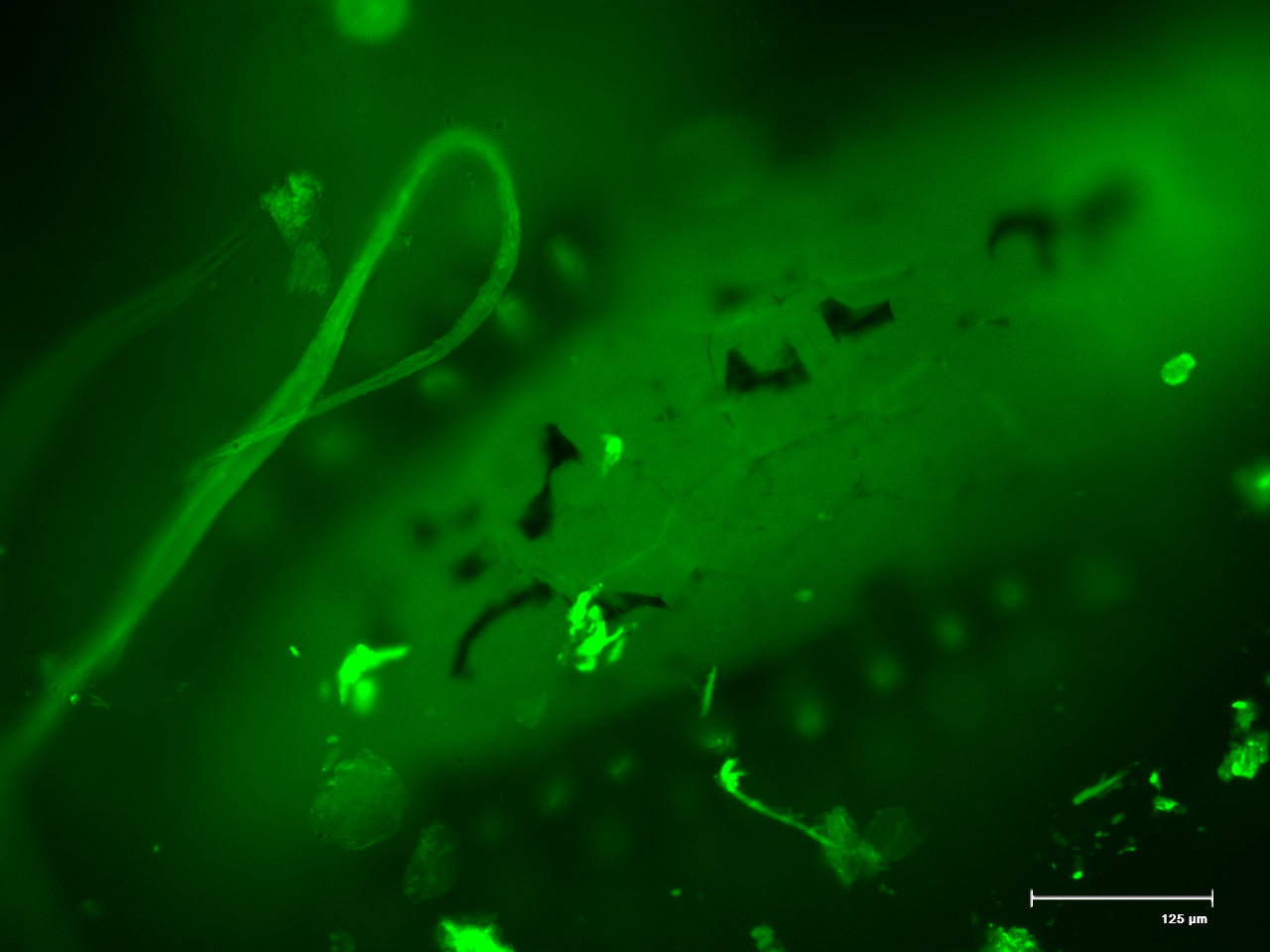

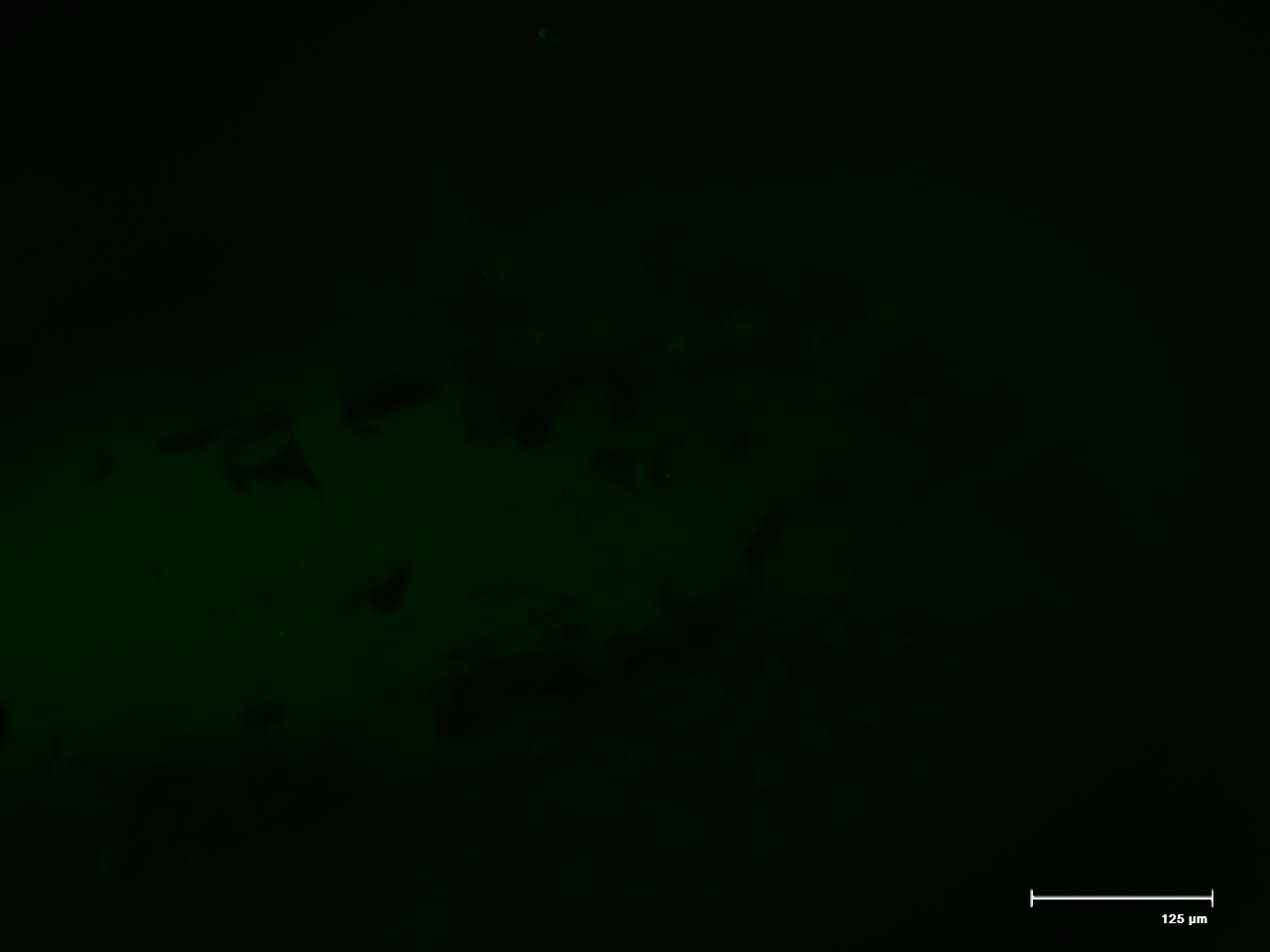

**A**

Col-0

-Fe

+Fe

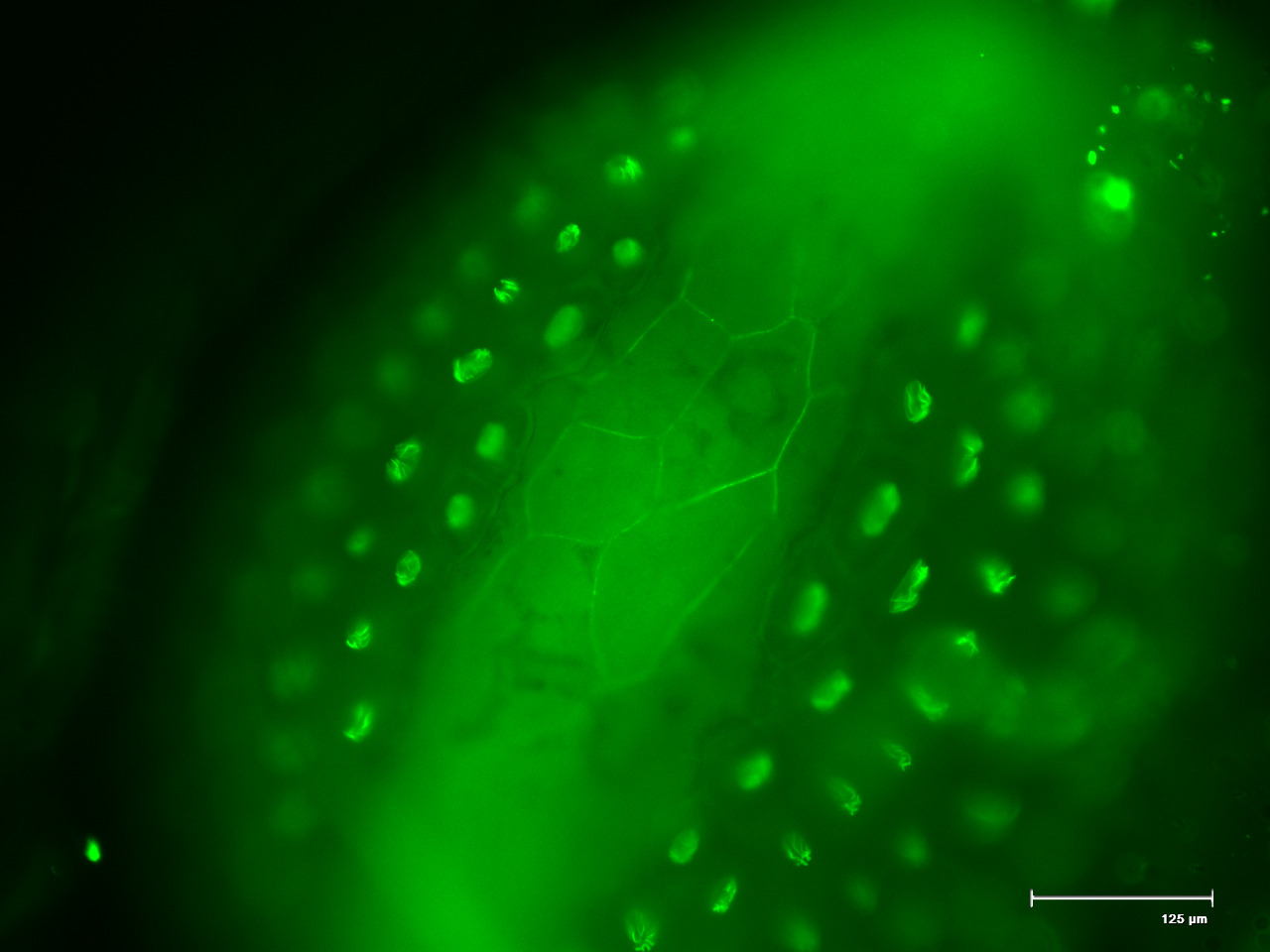

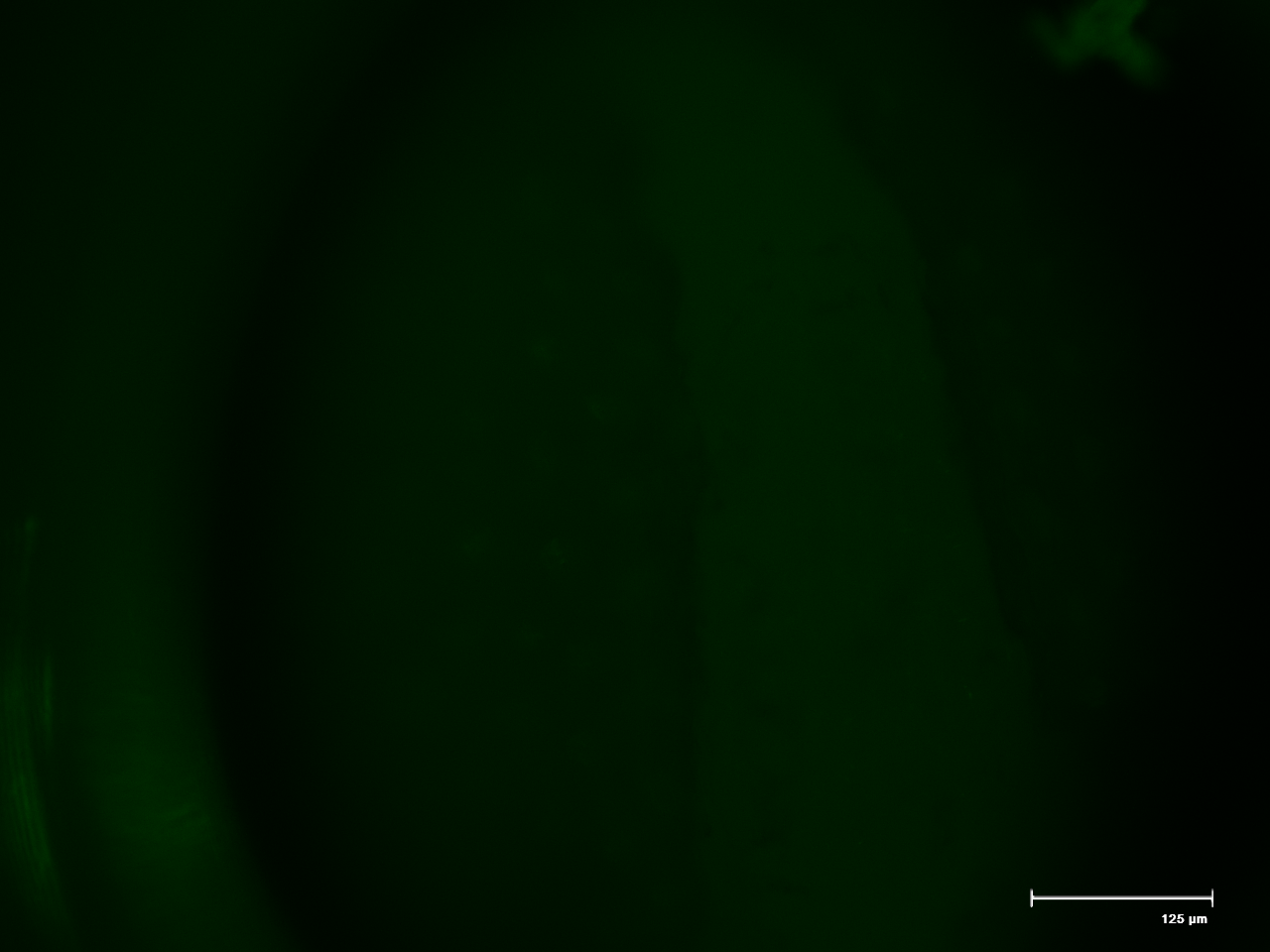

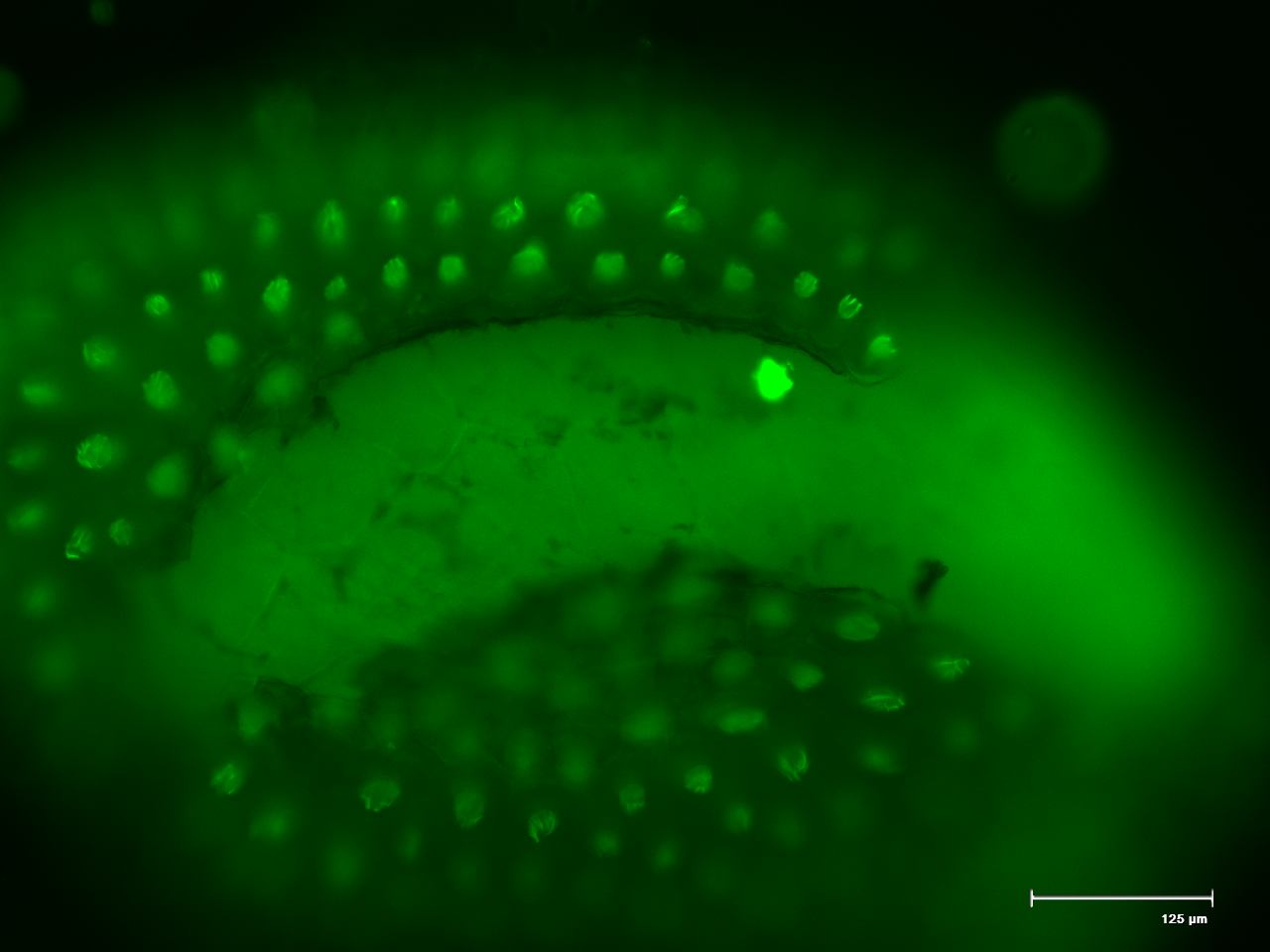

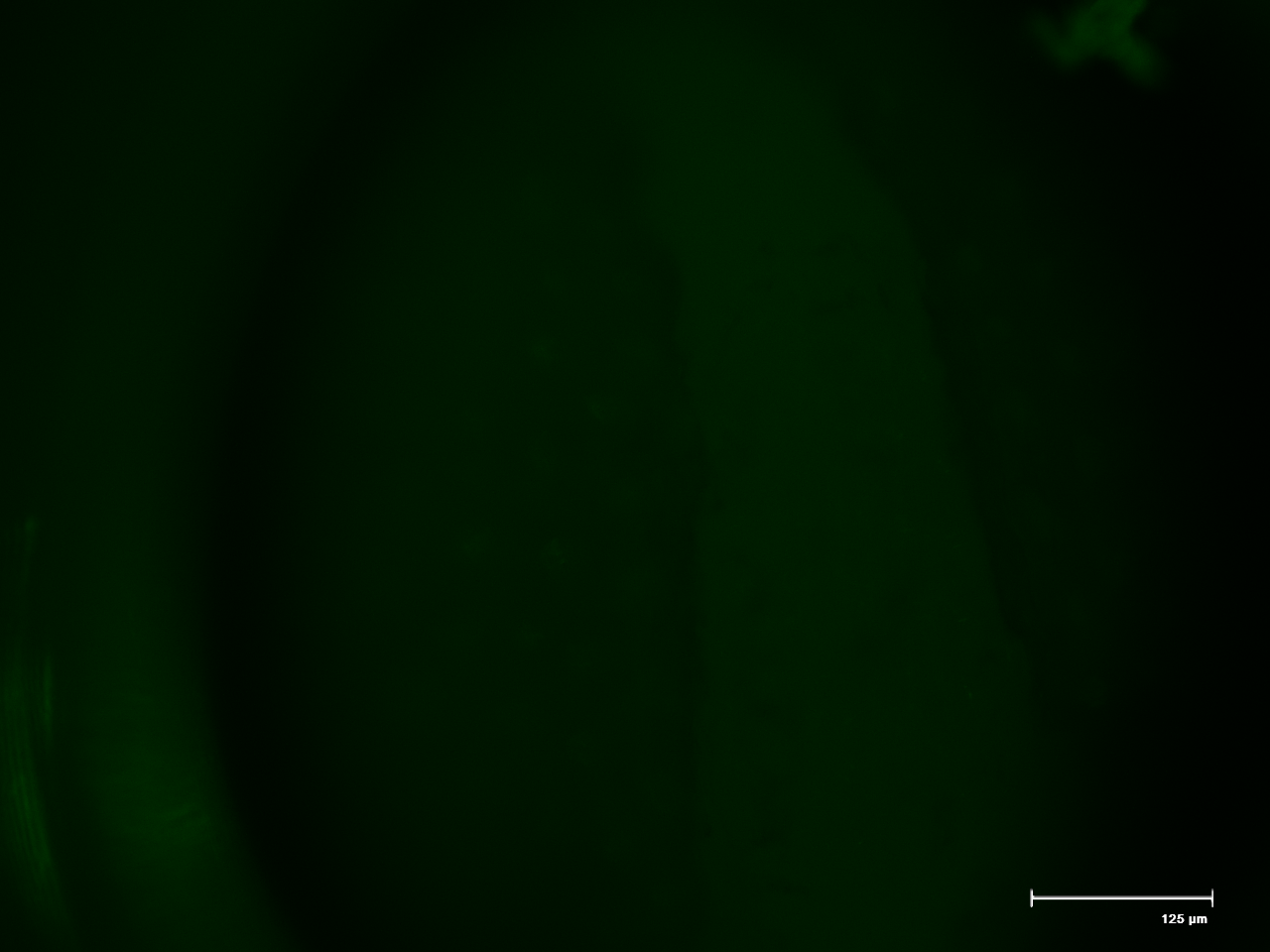

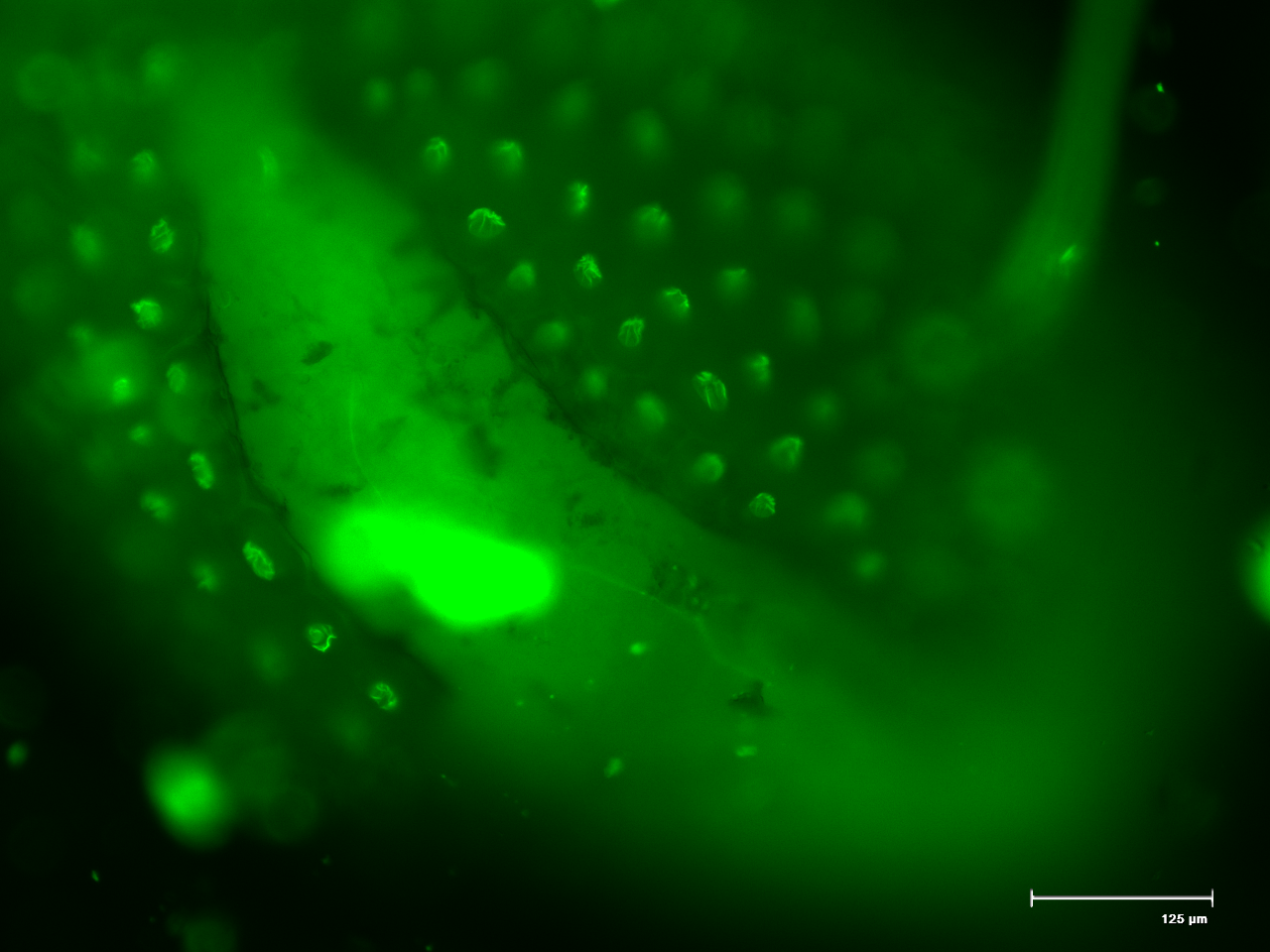

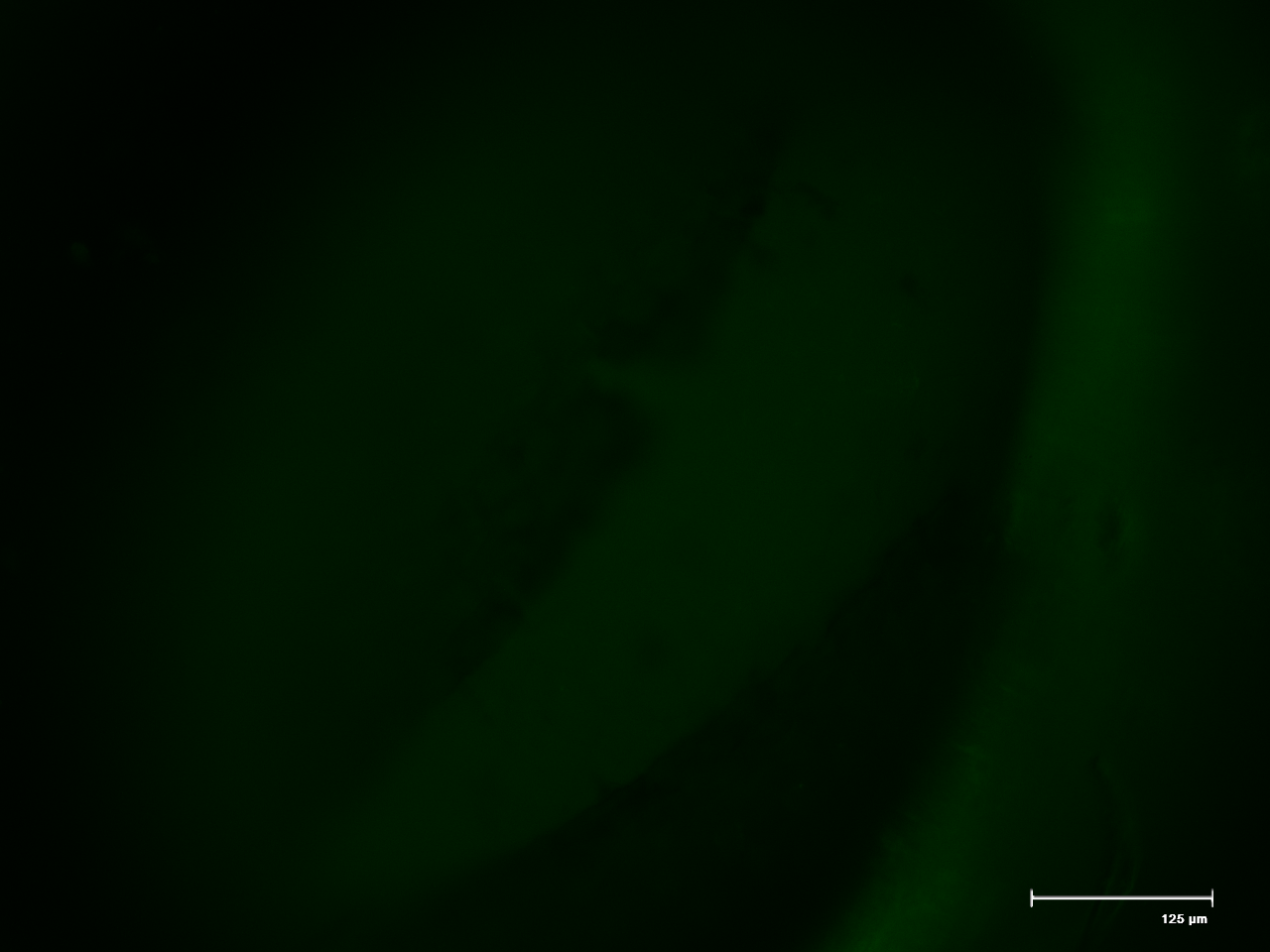

+Fe +HRP

Figure S6: PRX17 is essential for Fe triggered cuticle remodelling. A, epifluorescence microscopy of auramine O stained testa-ruptured seeds from wild type (Col-0) and *prx17* mutants. Seeds were induced to rupture on agar containing 30 µM ABA and 10 mM KNO_3_, then transferred for 24 hours to media supplemented with 500 µM Fe-EDTA (+Fe) or 500 µM Fe-EDTA + 500 ng/mL horseradish peroxidase (+Fe+HRP). White arrowheads indicate the continuous lipid barrier. Insets show background autofluorescence.

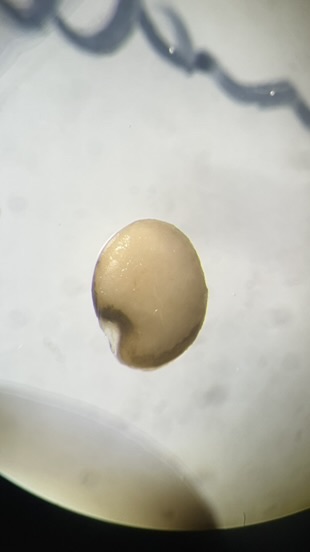

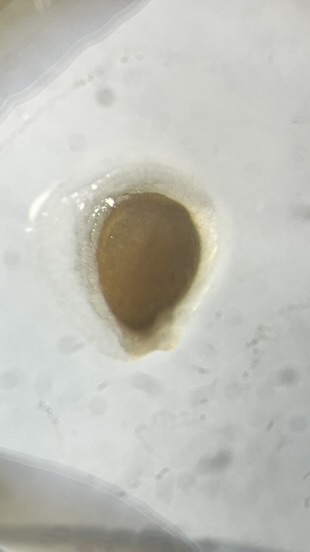

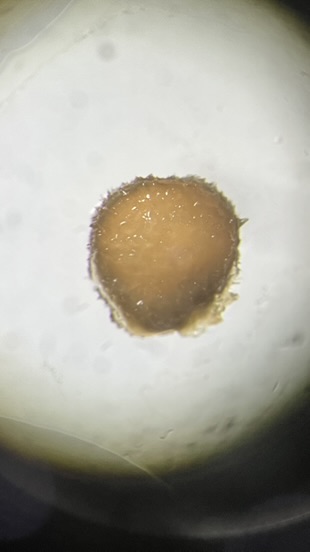

**Candella**

**Cobra**

**Moneymaker**

**48h	24h	0h**

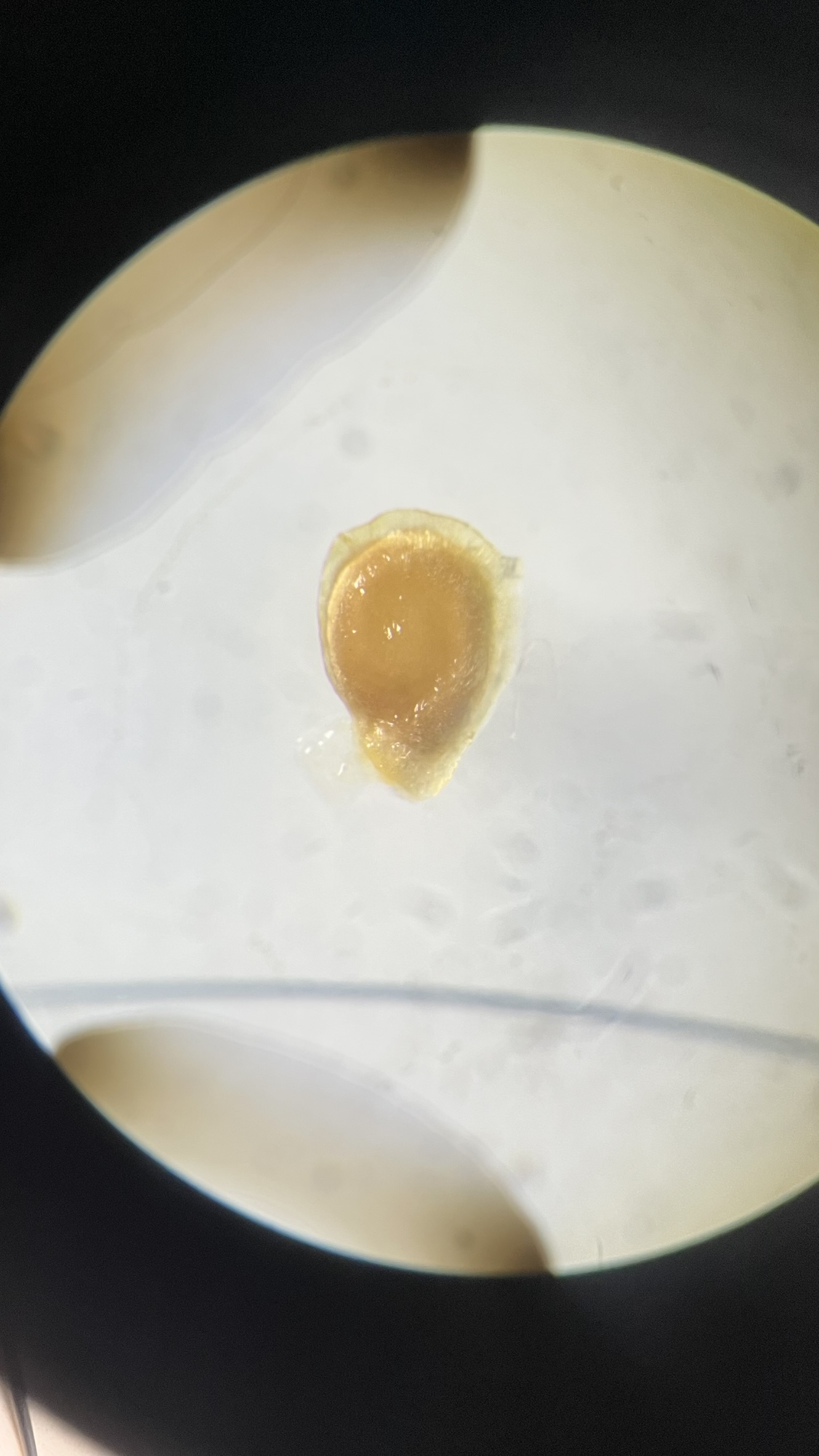

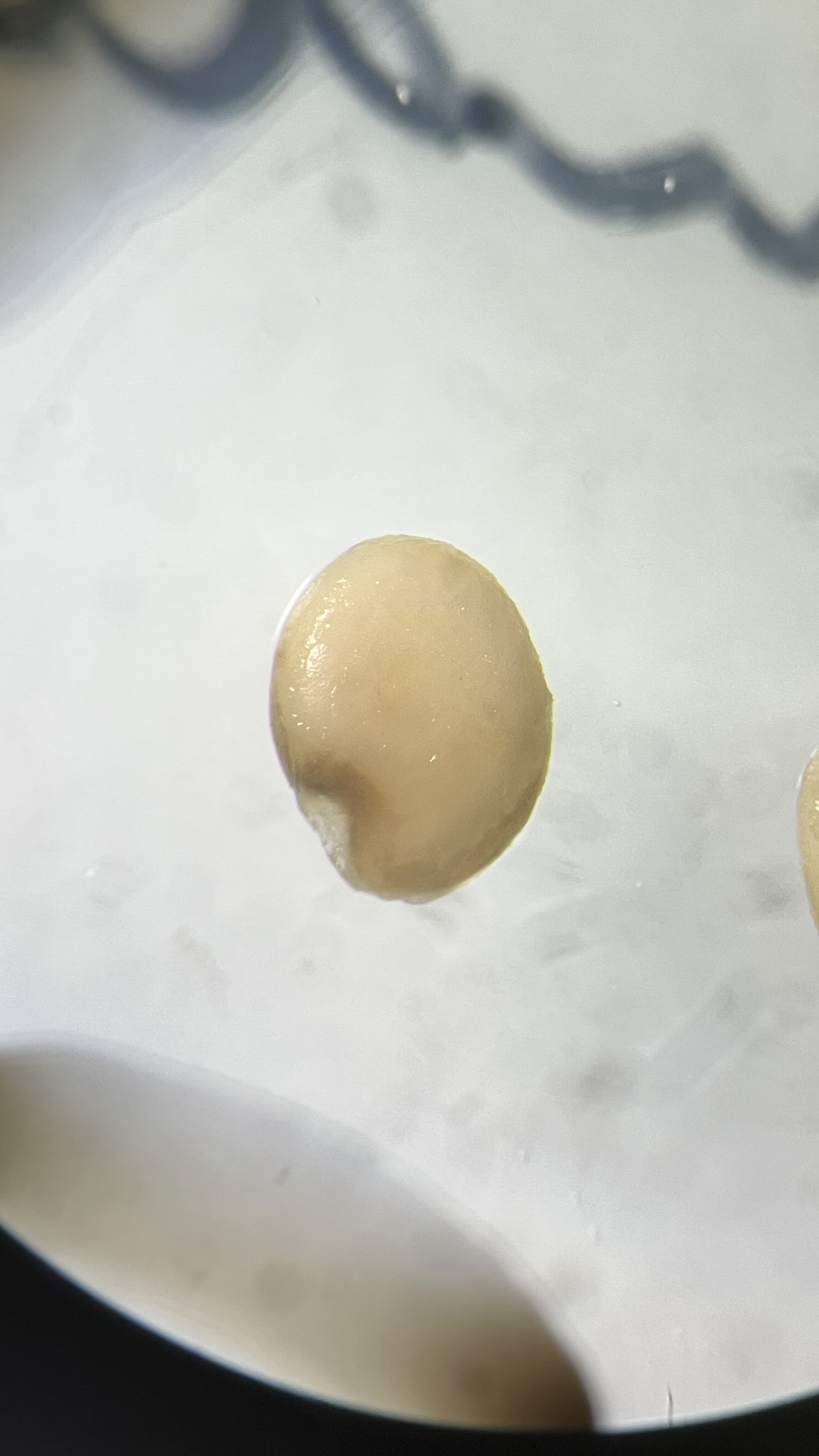

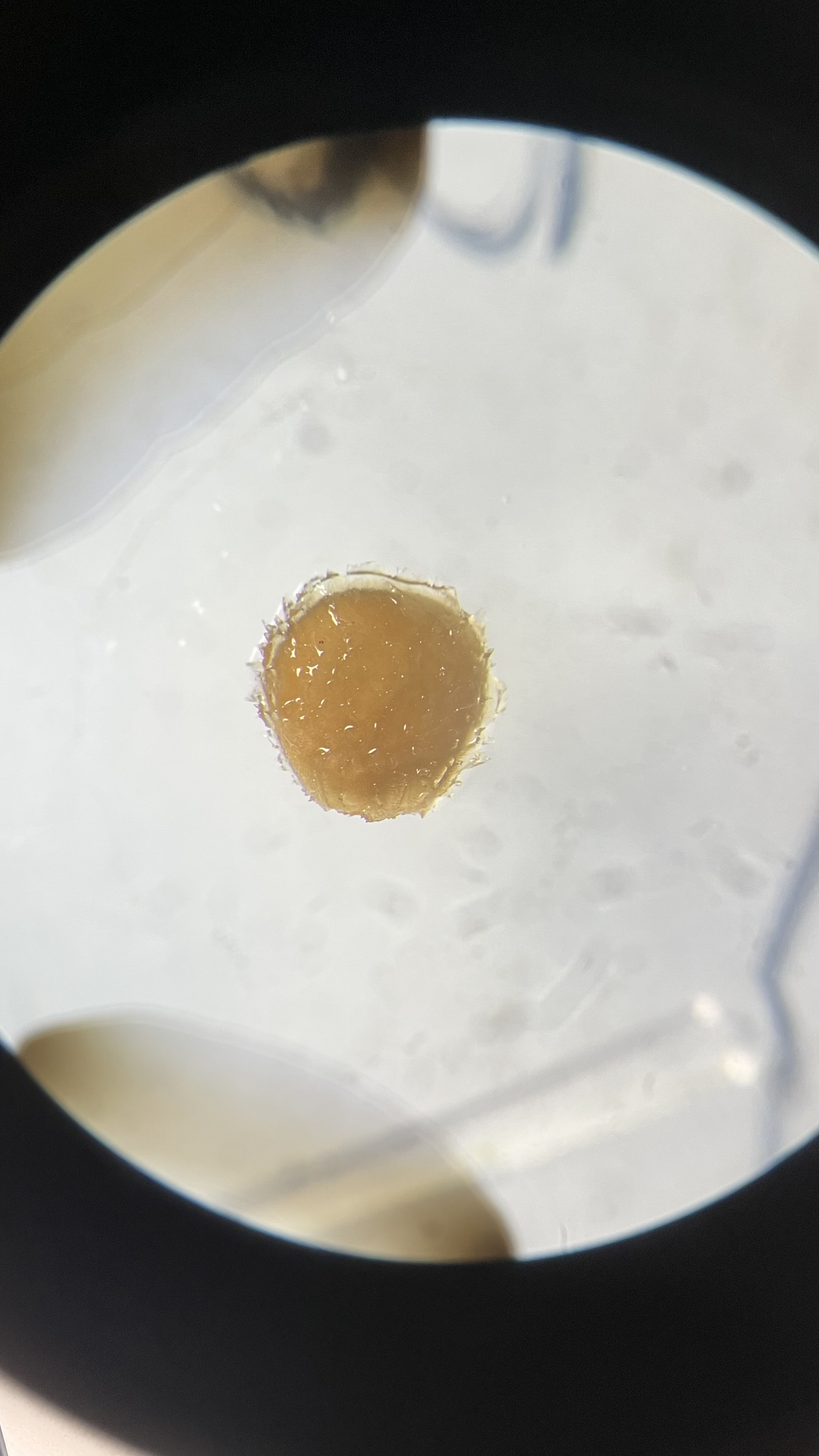

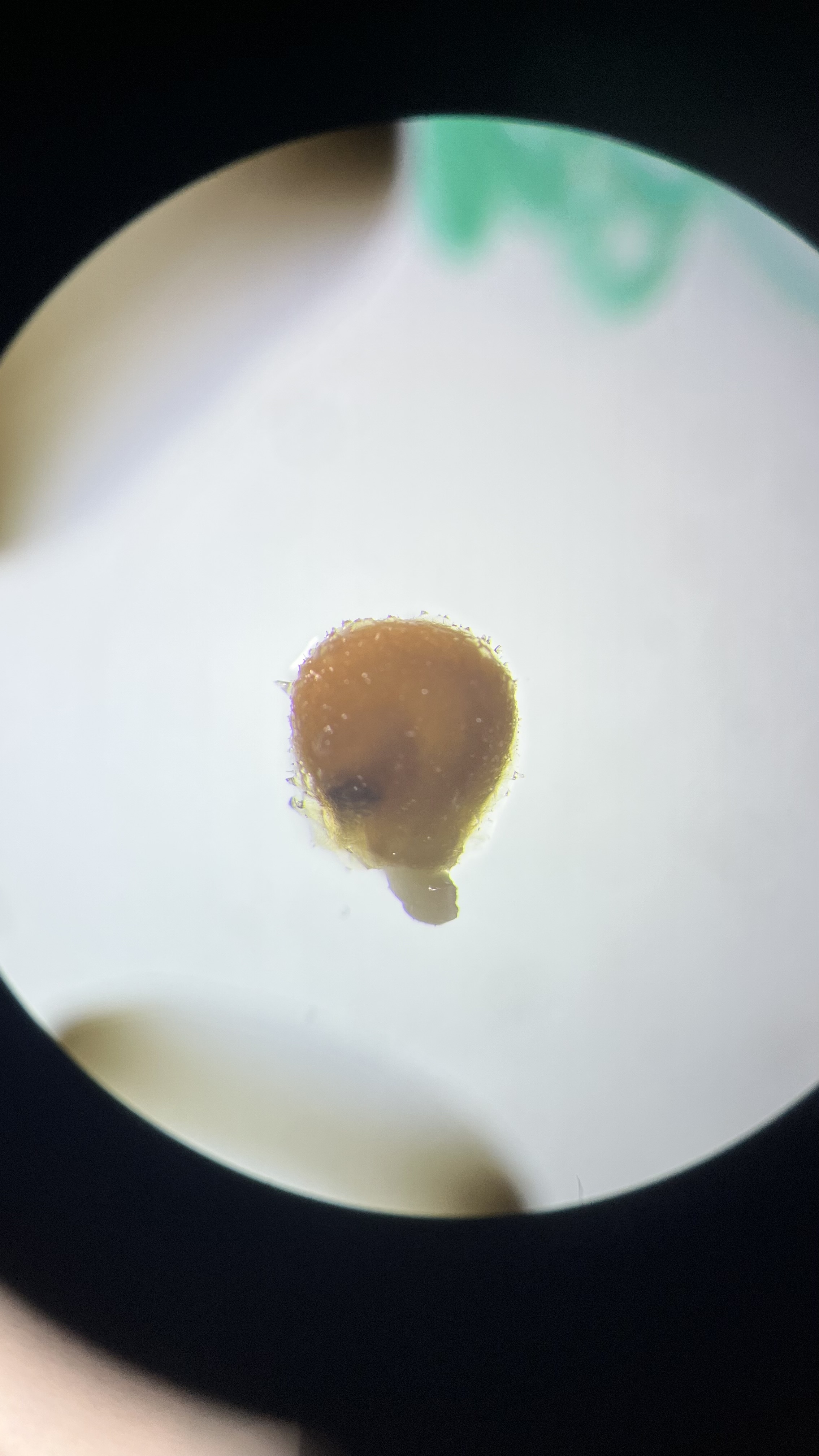

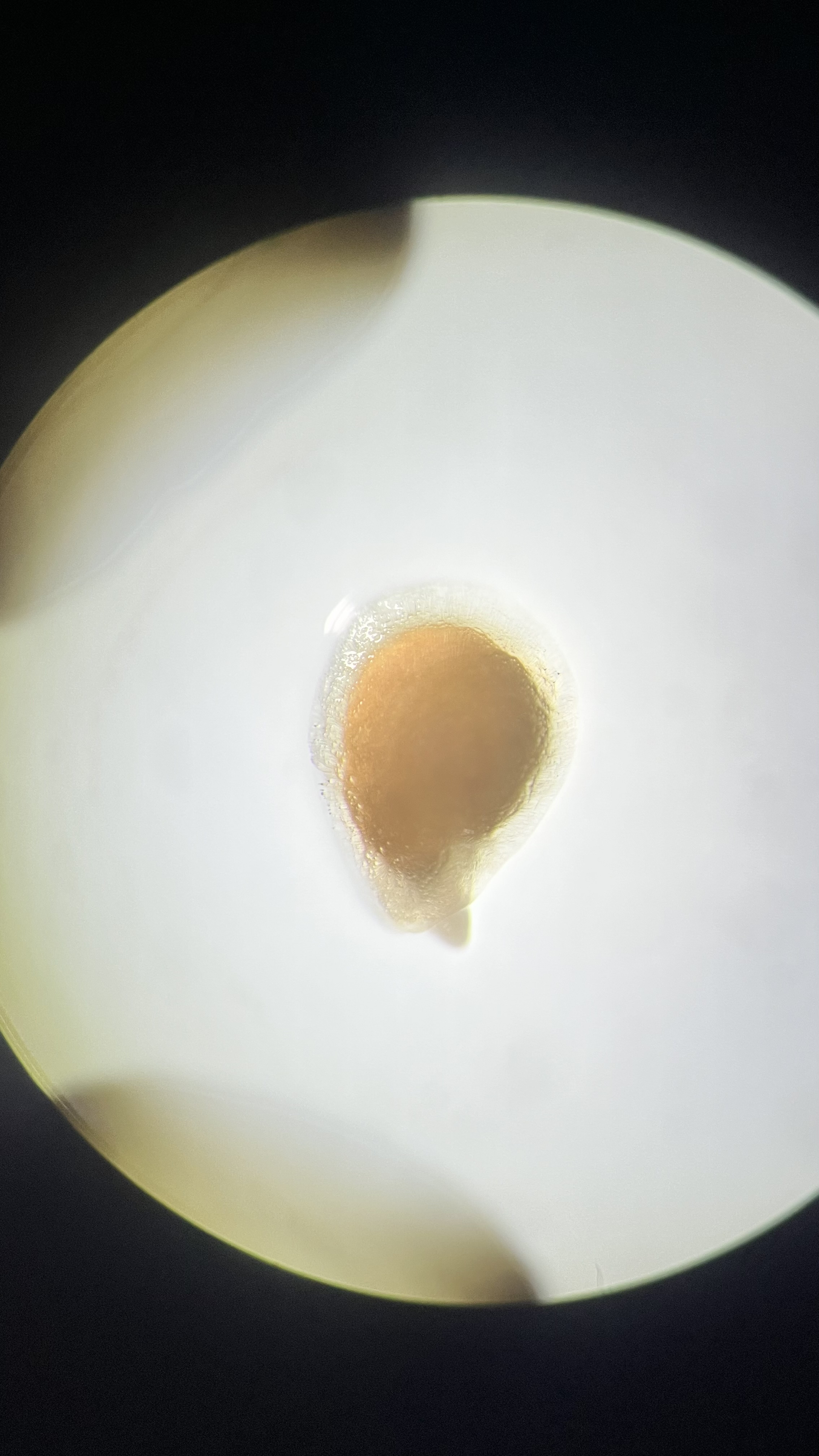

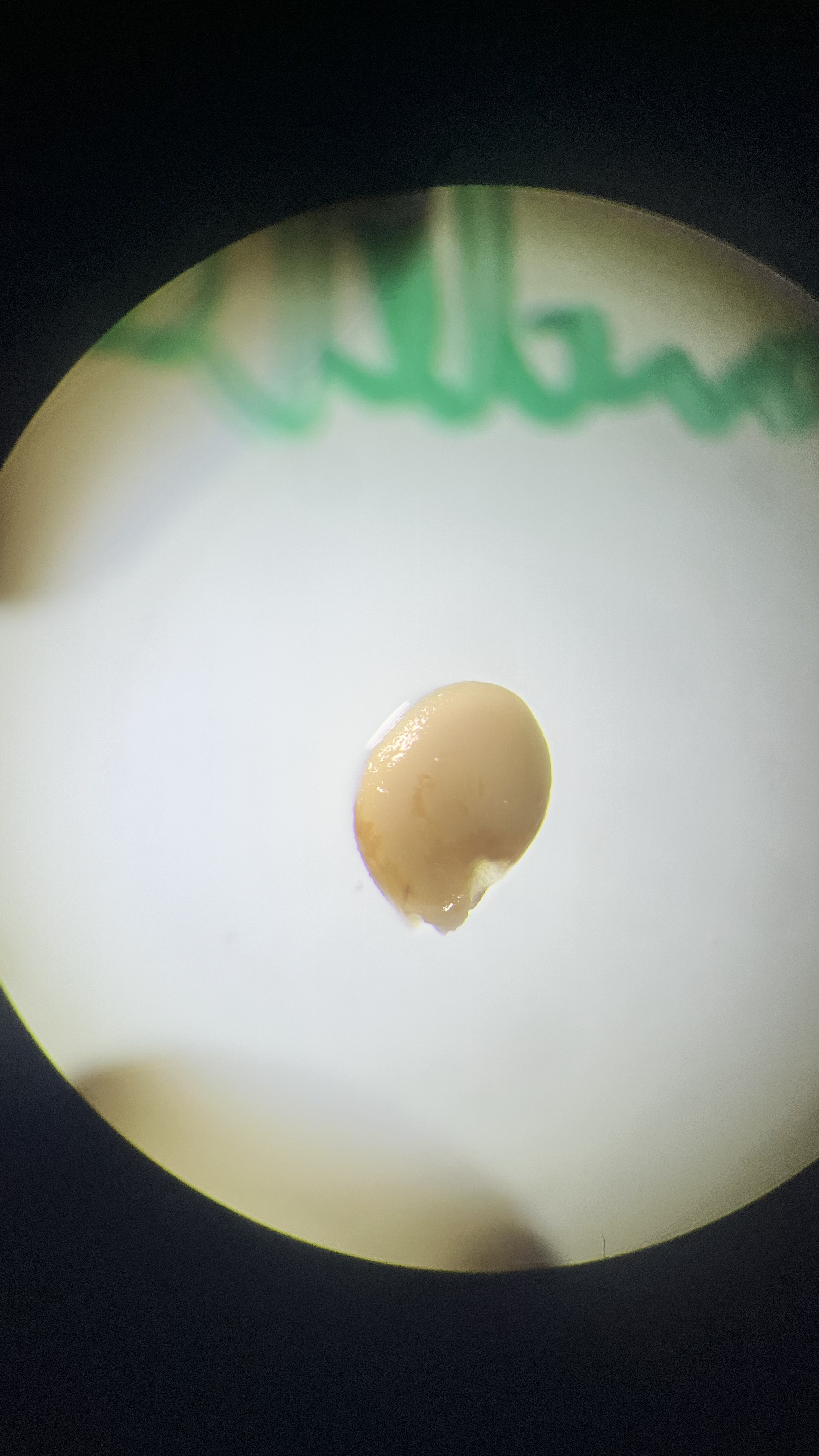

**EDTA	FeCl_2_**

Figure S7: The Fe-induced blackening response is conserved in tomato and is reversible. Time-course of pigmentation changes in seeds of three tomato cultivars (Candella, Moneymaker, Cobra). Seeds were imbibed on filter paper saturated with 250 µM FeCl_2_ for 24 hours (middle row), followed by transfer to 20 mM EDTA for an additional 24 hours (bottom row). Note that the dark pigmentation induced by Fe is chemically reversible upon chelation. Representative images were selected from at least 15 seeds tested per cultivar.

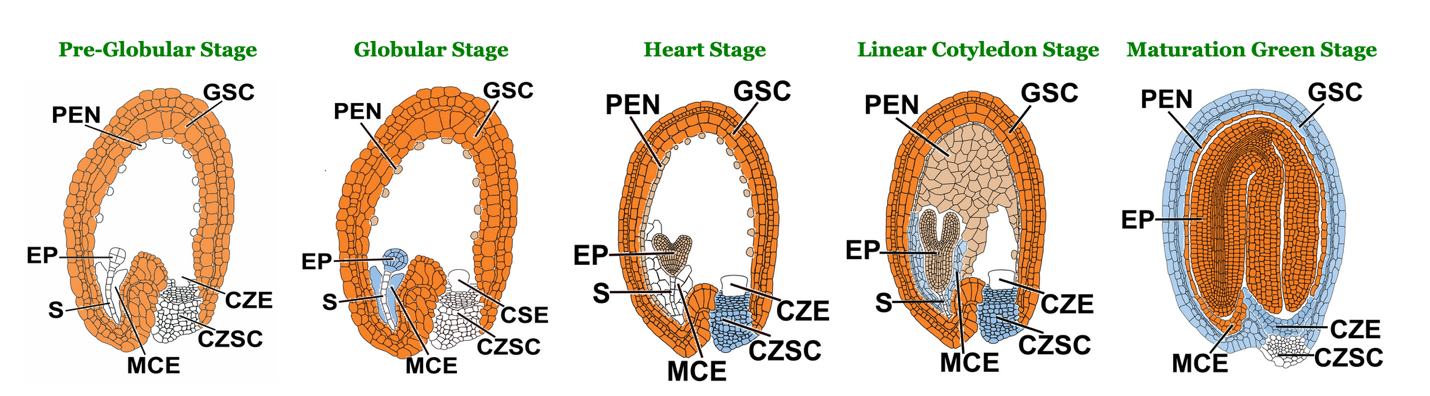

*FLS1*

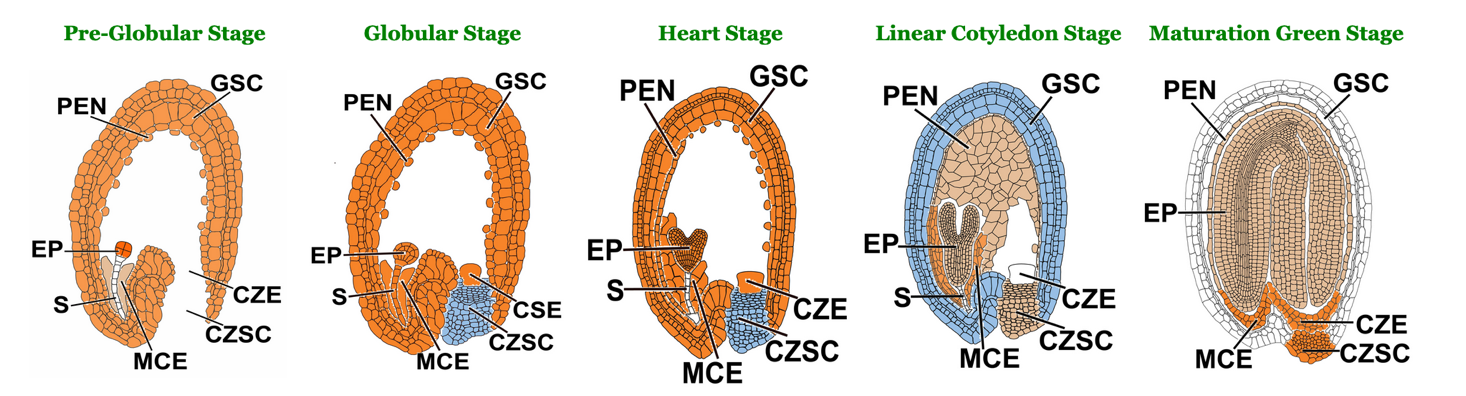

*PRX17*

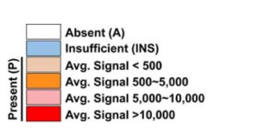

Figure S8: Expression of *FLS1* and *PRX17* during seed development. Data were retrieved from www.seedgenenetwork.net. Abbreviations: tissue/compartment: CZE, chalazal endosperm; CZSC, chalazal seed coat; EP, embryo proper; GSC, general seed coat; MCE, micropylar endosperm; PEN, peripheral endosperm; S, suspensor; WS, whole seed.

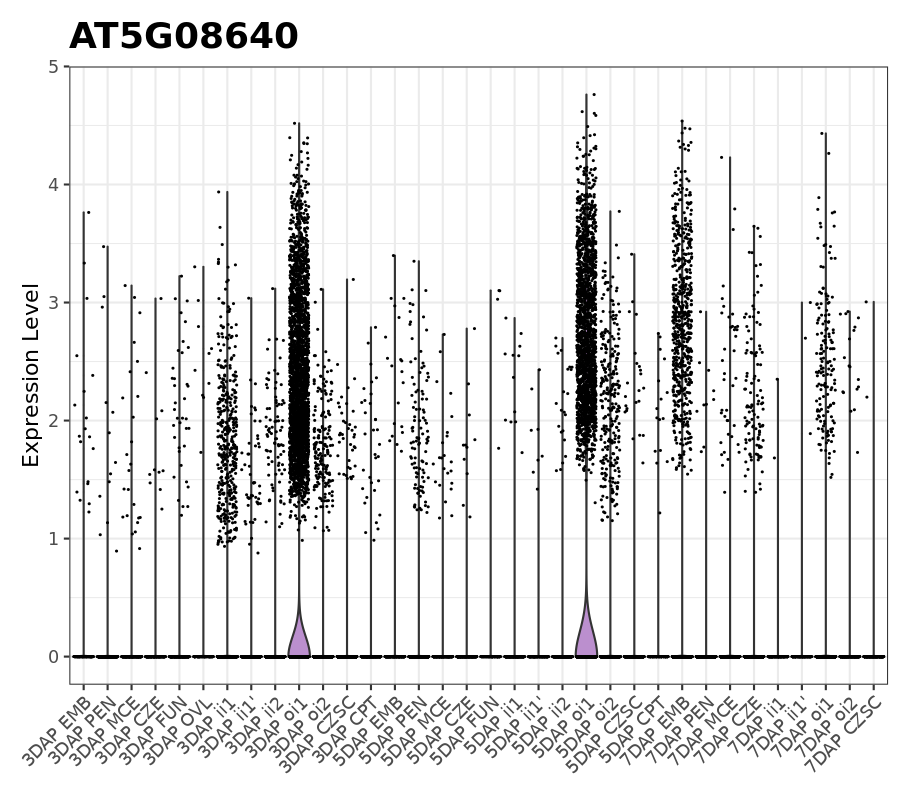

*FLS1*

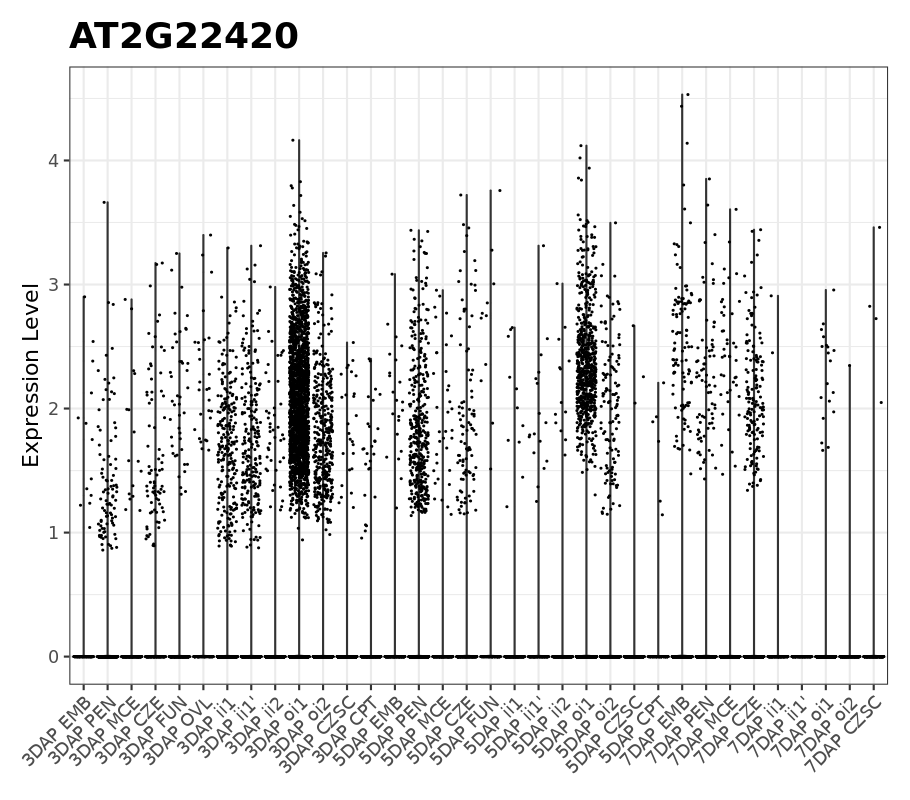

*PRX17*

Figure S9: *FLS1* and *PRX17* expressions temporally and spatially overlap during early seed coat development. *in silico* expression analysis of *FLS1 (Flavonol Synthase 1)* and *PRX17 (Peroxidase 17)* during seed development. Data were retrieved from the Arabidopsis Seed Gene Expression Atlas (https://seedatlas.wi.mit.edu). Note the concurrent high expression levels of both genes in the outer integument 1 (oi1) and inner integument 1 (ii1) layers between 3 and 5 days after pollination (DAP), supporting the hypothesis that flavonols and peroxidase are co-regulated during early seed coat formation.

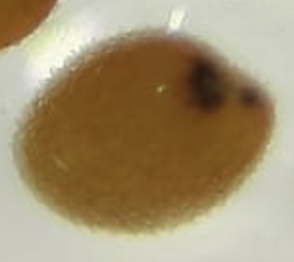

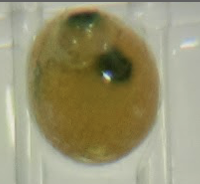

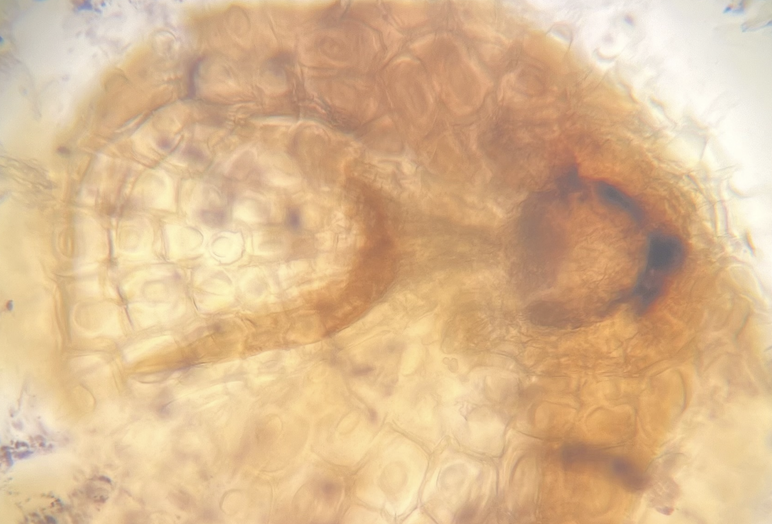

Figure S10: Spatial colocalization of flavonoid-Fe complexation, peroxidase activity, and Fe in the chalazal ring. Left, *in situ* visualization of the Fe-flavonoid complex. Seeds were imbibed in 250 µM FeCl_2_; the specific black pigmentation indicates the presence of flavonoid-Fe complexation. Middle, localization of peroxidase activity using TMB staining. Seeds were imbibed in 250 µM Fe-EDTA, and the blue signal indicates peroxidase activity. Right, localization of internal Fe stores. Dissected seed coat caps were stained with Perls/DAB, where dark precipitate indicates Fe accumulation. White arrowheads indicate the conserved chalazal ring structure observed across all three markers. Representative images were selected from at least 10 examined seeds per treatment.

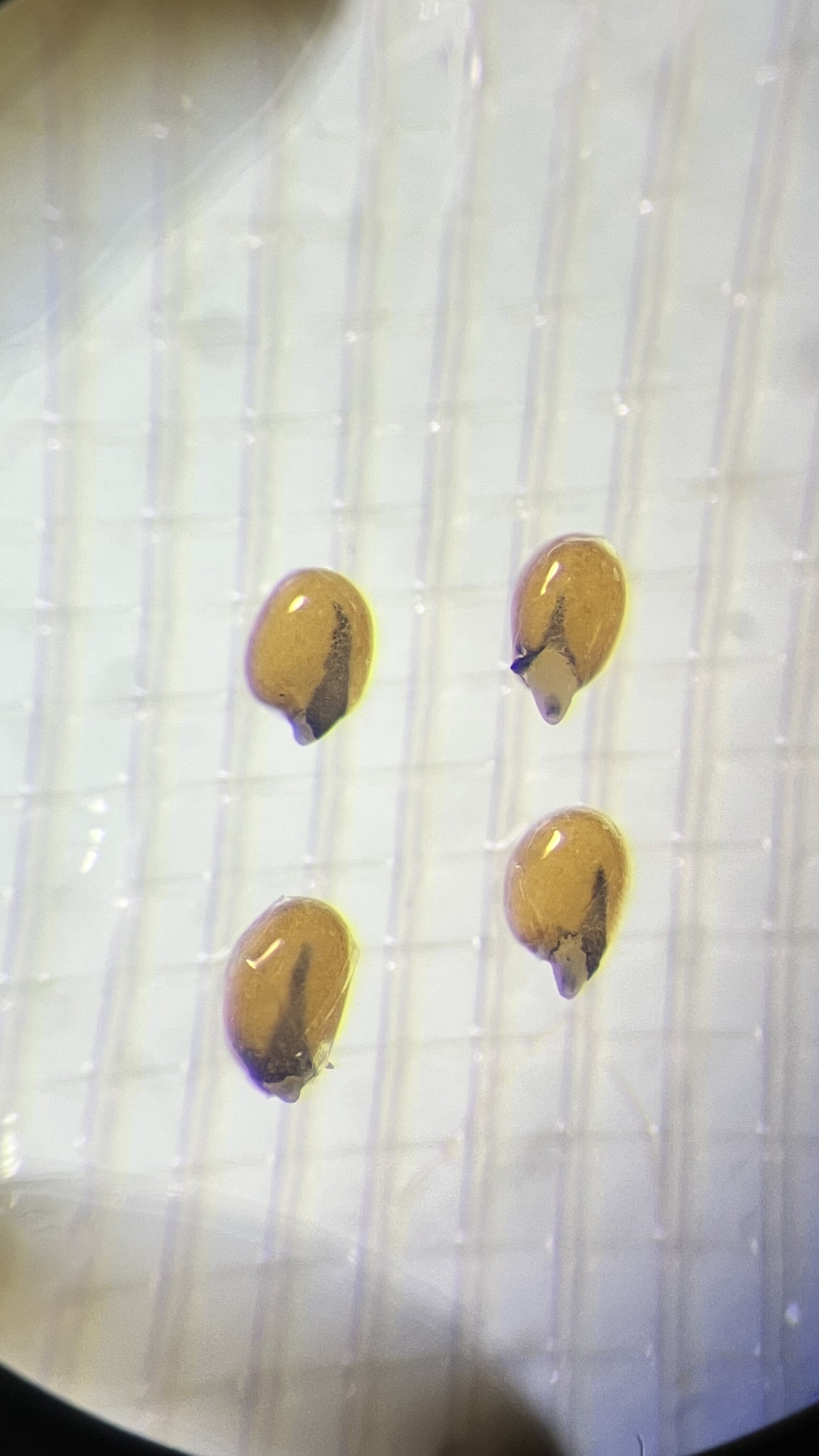

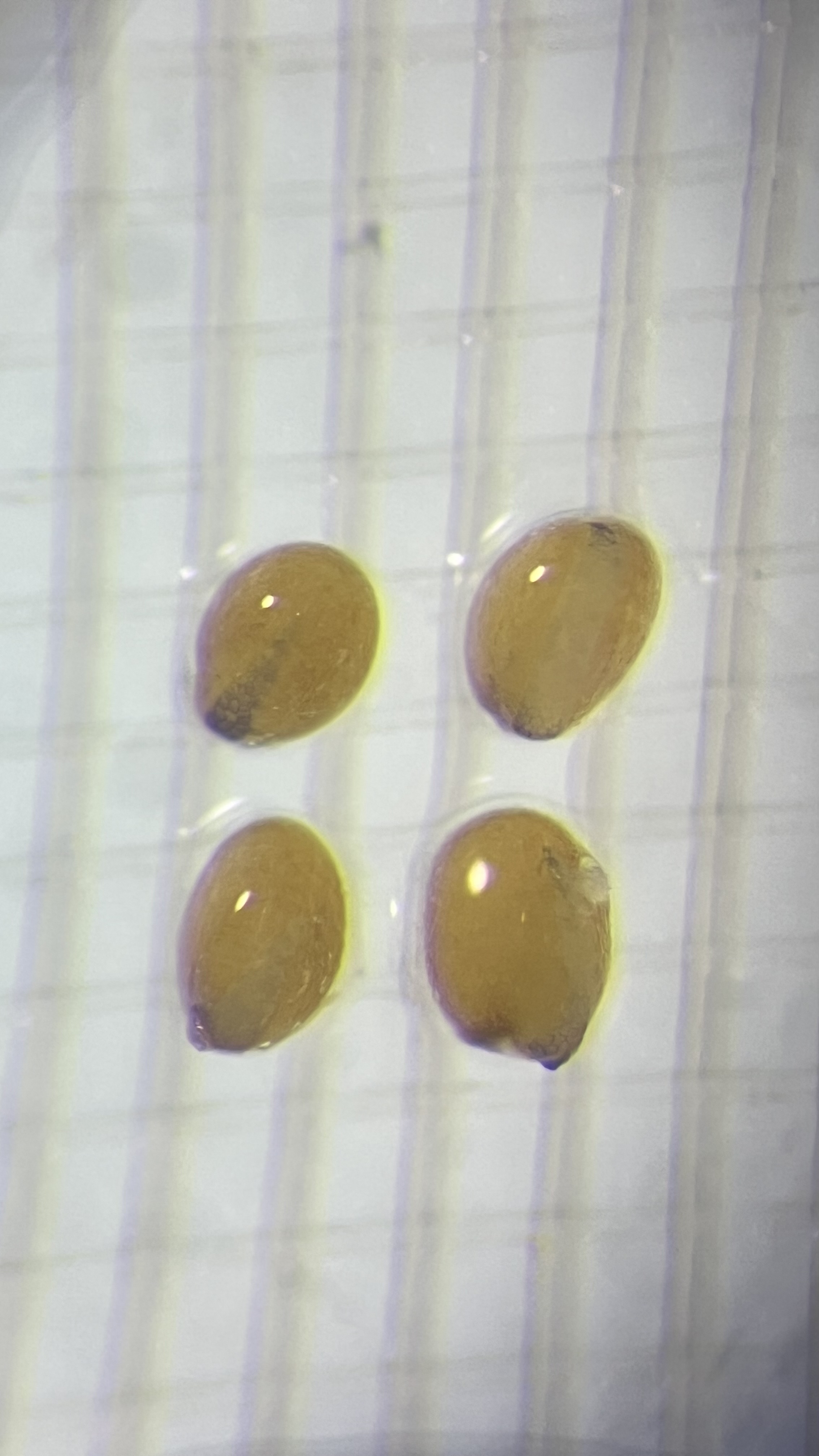

-Q	+Q

Figure S11: Exogenous quercetin suppresses Fe-induced peroxidase activity in the seed coat and endosperm. Visualization of peroxidase activity in seeds imbibed with 30 µM ABA and 250 µM Fe-EDTA, either without (-Q) or with (+Q) 100 µM quercetin. Peroxidase activity was detected using chloronaphthol staining. Representative images were selected from at least 10 examined seeds per treatment.

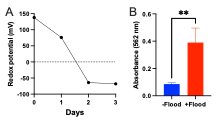

Figure S12 Flooding induces a reducing environment and increases bioavailable Fe²⁺ in soil. A, Impact of flooding on soil redox potential. Soil was flooded, and an oxidation-reduction potential (ORP) meter was used to record the redox potential (mV) over three days. The decrease in redox potential indicates a shift toward a reducing environment, driven by anaerobic microbial activity, which facilitates the reduction of metals. Data shown are representative of three independent experiments. B, Impact of flooding on Fe²⁺ availability. Soil was either left unflooded (-Flood, day 0) or flooded for 3 days (+Flood) in a growth chamber. Samples were extracted, mixed with ferrozine, and absorbance was measured at 562 nm to quantify relative Fe²⁺ levels. Error bars represent ± SEM. Asterisks indicate statistical significance (*p*=0.08, Student’s t-test;  n=4).
